# The role of ANGIOGENIN, or lack thereof, in the generation of stress-induced tRNA halves and of smaller tRNA fragments that enter Argonaute complexes

**DOI:** 10.1101/737114

**Authors:** Zhangli Su, Canan Kuscu, Asrar Malik, Etsuko Shibata, Anindya Dutta

## Abstract

Overexpressed Angiogenin (ANG) cleaves tRNA anticodons to produce tRNA halves similar to those produced in response to stress, but it is not clear whether endogenous ANG is essential for producing the stress-induced tRNA halves. It is also not clear whether smaller tRNA fragments (tRFs) are generated from the tRNA halves. Global short RNAseq experiments reveal that ANG over-expression selectively cleaves a subset of tRNAs (tRNA^Glu^, tRNA^Gly^, tRNA^Lys^, tRNA^Val^, tRNA^His^, tRNA^Asp^ and tRNA^SeC^) to produce tRNA halves and 26-30 bases long tRF-5s. Surprisingly, knockout of ANG reveals that the majority of stress-induced tRNA halves except 5’ half from tRNA^HisGTG^ and 3’ half from tRNA^AspGTC^ are ANG-independent, suggesting there are other RNases that can produce tRNA halves. The 17-25 bases long tRF-3s and tRF-5s that could enter into Argonaute complexes are not induced by ANG overexpression, suggesting that they are generated independently from tRNA halves. Consistent with this, knockout of ANG did not decrease tRF-3 levels or gene-silencing activity. Therefore ANG cleaves specific tRNAs, is not the only RNAse that creates tRNA halves and the shorter tRFs are not generated from the tRNA halves or from independent tRNA cleavage by ANG.

---

In the last decade, advances in next-generational sequencing have sparked the discovery of groups of previously unnoticed small non-coding RNAs (1,2). One among these groups includes small RNAs derived from transfer RNAs (tRNAs), so-called tRNA-derived RNA fragments (tRFs, tDRs) (3) or tRNA-derived small RNAs (tsRNAs) (4). tRFs have been discovered in multiple organisms and tissues (5–10). Based on the current knowledge about tRNA fragments, there are six major groups (Fig. 1A): (a) 5’ tRNA halves, (b) 3’ tRNA halves, (c) tRF-3s, (d) tRF-5s, (e) tRF-1s and (f) misc-tRFs. (a, b) The tRNA halves are also called tRNA-derived stress-induced RNAs (tiRs) that are produced by cleavage at anticodon arm of the mature tRNAs, linked with various stress response in diverse organisms (11–17). The two halves produced by tRNA cleavage are named respectively as 5’ tRNA halves and 3’ tRNA halves. 5’ tRNA halves have been shown to inhibit global translation and promote stress granule assembly (18,19), whereas 3’ tRNA halves interact with cytochrome C and protect cells from stress-induced apoptosis (20). (c) Similar to 3’ tRNA halves, tRF-3s also end at the 3’ end of mature tRNAs, but start by cleavage within the T-loop. Although these 17-25 nt (nucleotide) tRF-3s (major peaks at 18 nt and 22 nt) could load into Argonaute complexes to perform microRNA-like gene silencing activities (5,21,22), they are still generated in Dicer and Drosha deficient cells (5,21). tRF-3s have also been implicated in retrotransposon inhibition (23) and ribosome biogenesis (24). (d) tRF-5s start at the 5’ end of mature tRNAs and end before the anticodon loop. 17-25 nt (“short”) tRF-5s have been shown to inhibit protein translation (25–27), and also suggested to play potential microRNA-like functions (5). Another distinct group of 26-30 nt (“long”) tRF-5s, also called tRNA-derived Piwi-interacting RNAs (td-piRNAs), could enter PIWI complex to regulate downstream pathways (28–30). (e) tRF-1s or 3’U tRFs correspond to the 3’ trailer from the precursor tRNAs during tRNA maturation process. The 5’ end of tRF-1 results from RNaseZ cleavage and the 3’ end correlates with termination by PolIII transcription. tRF-1s have been linked with cancer (3,31,32). (f) Misc-tRFs include the other types of tRFs that are not represented by the above groups, such as internal tRFs (i-tRFs, also called tRF-2s) that start and end within the mature tRNAs. Internal tRFs that start at D-loop and end at variable loop or T-loop displace YBX1 from 3’UTR of oncogenic transcripts to suppress breast cancer progression (33). Overall there is a growing interest in the functions of tRNA fragments, and evidence so far suggests they might play divergent roles in different biological pathways via specific mechanisms.

**Figure 1.**
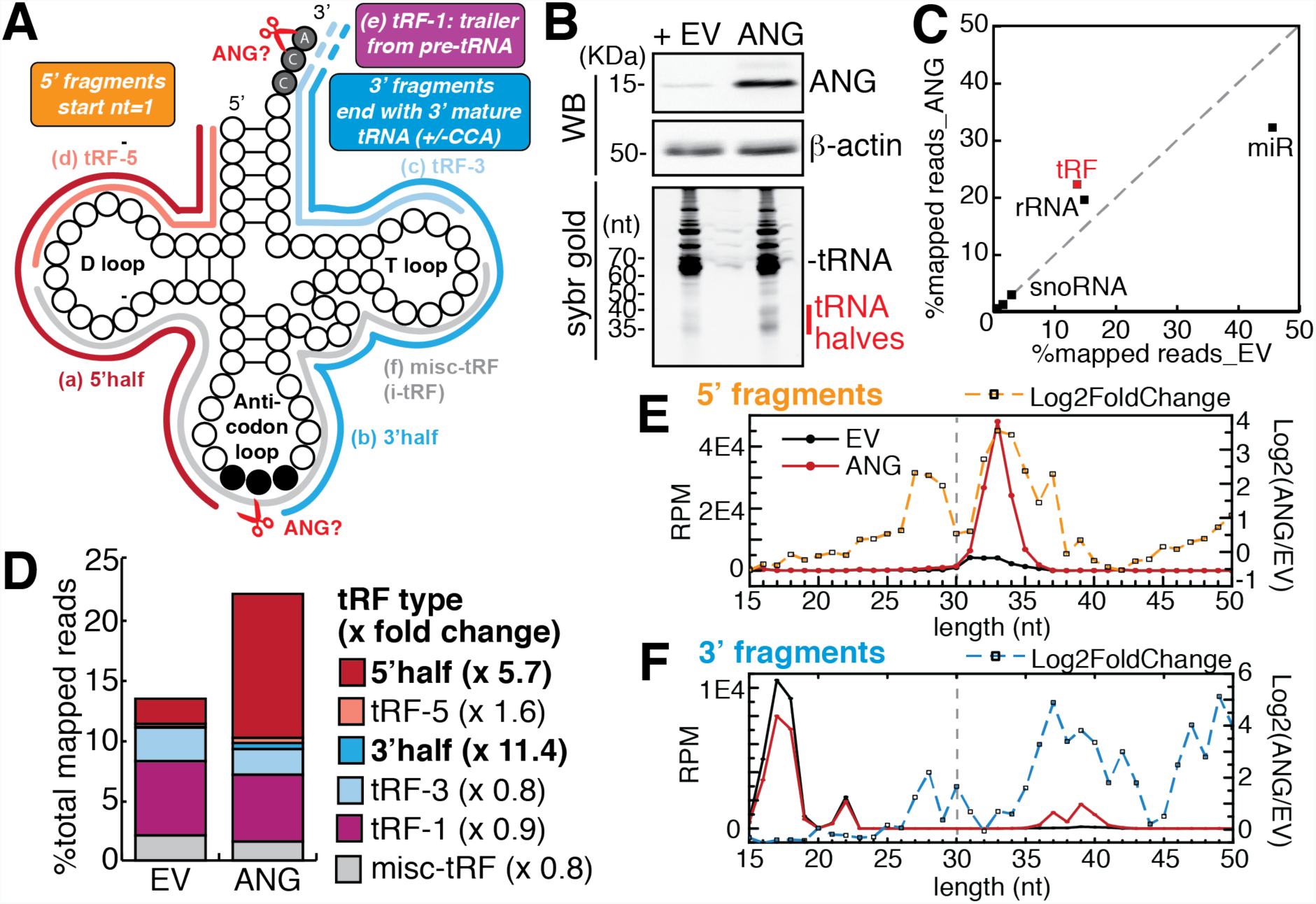
Angiogenin overexpression increases tRNA halves. (A) Different types of tRNA fragments. Refer to introduction for the description of each type. Potential cleavage sites by ANG are indicated by scissors. (B) Over-expression of ANG in HEK293T cells (EV: empty vector) induces tRNA half bands. Upper panel (WB: western blot): specificity of ANG antibody was shown in Fig. S1B. Lower panel: total RNA was separated on 15% Urea-PAGE and stained by Sybr gold. (C-F) ANG over-expression specifically increases tRNA halves, shown by small RNA-seq analysis averaged from 3 biological replicates. (C) Scatter plot shows Log 10 of read counts mapped to each category from small RNA-sequencing analysis (X-axis: empty vector control, Y-axis: ANG overexpression). (D) ANG over-expression specifically increases overall reads of 5’halves and 3’halves. Here 5’half refers to 5’ fragments of >= 30 nucleotide in length, and tRF-5 refers to < 30 nucleotide (corresponding to vertical dashed gray line in panel E); similarly 3’half and tRF-3 are separated by length (dashed gray line in panel F). (E-F) Length distribution of all tRF reads that map to 5’ or 3’ end of mature tRNAs by small RNA-seq (Y-axis: reads per mapped million reads, black line: empty vector, red line: ANG overexpression). Dashed line shows log2 fold change upon ANG overexpression for tRFs of specific length (Y-axis on the right).

Despite the emerging interest in tRFs, how specific tRFs are generated still remains largely unknown. Most studies have focused on the biogenesis of the stress-induced tRNA halves. In human cells, stress-induced cleavage in tRNA has been mostly attributed to enzymatic cleavage by angiogenin (ANG or RNase 5), a member of the vertebrate-specific RNase A family (16–20,34–36). ANG is widely expressed in most tissues (37), and has been linked to angiogenesis, hematopoiesis, oncogenesis, inflammation and immunity (38); loss-of-function mutations in *ANG* have been implicated in neurodegenerative diseases such as ALS (amyotrophic lateral sclerosis) and Parkinson’s Disease (38). Under normal conditions, cellular ANG is inactive due to high affinity binding with RNH1 (RNase inhibitor 1) (39). Under stress conditions, cytoplasmic ANG is activated by dissociation from RNH1, which triggers tRNA cleavage (40). Recombinant ANG added directly to culture media increased tRNA anticodon loop cleavage in multiple cell lines (16–20,34–36); interestingly, ANG cleavage did not significantly reduce the overall tRNA abundance. One report also suggests stress-activated ANG could cleave the CCA at the 3’ end of mature tRNAs quickly, even before the cleavage at the anticodon loop (41) (Fig. 1A). Most of these previous studies quantified the effect of ANG cleavage only on a subset of tRNAs by Northern blot analysis, so a global view is still lacking. On the other hand, biogenesis of stress-independent tRFs is poorly known, except a recent report of ANG-dependent tRNA halves in non-stressed sex-hormone positive cells (42). We have shown that most 17-25 nt tRF-3s are generated by uncharacterized Dicer-independent pathways (5,21), and it has not been tested whether tRNA halves could serve as precursors for these and other shorter tRFs. This is a possibility especially considering that ANG could directly generate tRF-3 as suggested by *in vitro* reactions (43) or detach amino acids from tRNA 3’ CCA end (41), which might be a prerequisite for generating a fraction of tRF-3s.

In this report, we systematically characterize angiogenin’s role in tRNA cleavage in non-stressed and stressed human cell lines. We test whether ANG is sufficient and/or required for tRNA cleavage. By small RNA-sequencing, we profiled the global tRNA cleavage pattern by ANG overexpression, which leads to specific increase of both tRNA 5’ halves and 3’ halves. While increase of tRNA 5’ halves correlates with an increase in the long (26-30 nt) tRF-5s, tRF-1s, short (17-25 nt) tRF-5s and tRF-3s were unaffected by ANG cleavage. Surprisingly, the majority of tRFs were unaffected by *ANG* knockout and only a few specific tRNA halves (5’ tRNA^HisGTG^ and 3’ tRNA^AspGTC^) were decreased by loss of ANG. CRISPR knockout of ANG cells still displayed arsenite stress-induced tRNA anticodon cleavage and translational arrest, suggesting other RNase (s) contribute extensively to the overall stress-induced tRNA cleavage. We conclude that human angiogenin is sufficient but not essential for tRNA anticodon cleavage and that the short tRF-3s and tRF-5s are generated by ANG-independent mechanisms.

## Results

### ANG generates tRNA halves

To identify which tRNAs are cleaved and how by ANG, we over-expressed full-length human ANG in HEK293T cells (Fig. 1B and Fig. S1A). Western blot in these cells confirms that a 14-KDa mature ANG protein is overexpressed in the cell lysate (Fig. 1B and S1B). We noted ANG mRNA was over-expressed over 500-fold (Fig. S1A), while ANG protein appears to be over-expressed in cell lysates to ~10-20 fold (Fig. 1B), which could be due to the secretion of ANG protein from the cells. Sybr gold staining of total RNA detects RNA bands around 35 and 40 nucleotides upon ANG overexpression (Fig. 1B), consistent with the tRNA halves in previous reports (17,35,44).

To characterize the tRNA fragmentation pattern induced by ANG overexpression, we performed small RNA-sequencing of 15-50 nucleotide long RNAs from total RNA (workflow shown in Fig. S1C). This size range will capture tRNA fragments but not the full-length tRNAs (Fig. S1D). Sequencing of tRNA fragments is known to be sensitive to both terminal and internal modifications on the RNA molecules. In order to make quantitative comparison of tRNA fragments between two conditions, we assumed the cloning efficiency for each tRF is not changed by different conditions of cell treatment. This is why we compare relative levels of the same tRF across conditions, rather than the levels of different tRFs in a given condition. ANG overexpression specifically increased counts of fragments derived from tRNAs, and fragments from ribosomal RNAs (rRNAs) to a lesser extent (Fig. 1C). The slight increase in rRNA fragment reads is consistent with previous reports on ANG’s role in rRNA transcription and processing (45–47). Despite the relative global decrease in microRNA reads upon ANG overexpression (Fig. 1C), no microRNA showed significant decrease upon ANG over-expression by differential analysis using DESeq2 (48) on the union of microRNAs and tRFs (Table S2), which suggests ANG is unlikely to decrease specific microRNA levels.

Among the different types of tRFs (Fig. 1A), 5’ halves (≥30 nt) increase by 5.7 fold upon ANG overexpression, tRF-5s (<30 nt) are only mildly increased (Fig. 1D). Similarly, 3’ halves (≥30 nt) increase by 11 fold, whereas tRF-3s (<30 nt) are not increased (Fig. 1D). Here we define 3’ fragments irrespective of CCA presence to account for potential cleavage of the 3’ CCA at CA dinucleotide by ANG (41). Despite this precaution, it is worth noting that most of the 3’ fragments contain intact 3’ CCA sequence (examples shown in Fig. S2A). tRF-1s, produced from the trailer sequence through cleavage by RNAse Z, was also unchanged by ANG overexpression (Fig. 1D). When we looked at tRFs by size, ANG overexpression most specifically increases 5’ tRNA halves in the size range of 31-36 nucleotides (Fig. 1E), and 3’ halves of 36-41 nucleotides (Fig. 1F). In contrast, tRF-5s and tRF-3s of 17-25 nt were not much affected by ANG (Fig. 1E and 1F, dashed line represents log2 fold change). Overall, results from ANG overexpression suggest that ANG does not have a role in producing tRF-1s and the short (17-25 nt) tRF-3s and tRF-5s. Interestingly, long tRF-5s of 26-30 nt (also called td-piRNAs) were also increased by ANG overexpression (Fig. 1D), though it is not clear whether they are directly created by cleavage in the anticodon stem, or whether they are further processed from the 3’ end of the 5’ tRNA halves.

### ANG specifically increases tRNA halves from a small subset of parental tRNAs

To analyze whether ANG cleaves particular groups of tRNAs, we grouped tRNA fragments by the parental tRNAs. ANG overexpression up-regulates most of the tRNA halves detected by small RNA-seq (Fig. 2A and Fig. 2B). Interestingly, both 5’ and 3’ halves from certain tRNAs (including tRNA^Glu^, tRNA^Gly^, tRNA^Asp^, tRNA^Val^, tRNA^Lys^, tRNA^Ser^, tRNA^iMet^ and tRNA^SeC/selenocystein^), and 5’ half from tRNA^His^, are up-regulated significantly (adjusted p value less than 0.05) upon ANG overexpression (Fig. 2B). When we sum up the different types of tRFs by their parental tRNAs, a group of tRNAs (tRNA^Gly^, tRNA^Glu^, tRNA^Lys^, tRNA^Val^, tRNA^His^, tRNA^Asp^ and tRNA^SeC^) are clearly more ANG-responsive than the other tRNAs (Fig. 2C). The 5’ tRNA halves have much higher cloning frequency than 3’ halves for most of these tRNAs (e.g. tRNA^Glu^, tRNA^Gly^, tRNA^His^, tRNA^Lys^, tRNA^Val^) in both control cells and ANG over-expression cells (Fig. 2D and Fig. S2B). This could be a result of higher stability or higher clonability of the 5’ halves (further discussed in Discussion). Furthermore, these ANG-responsive tRNAs (tRNA^Gly^, tRNA^Glu^, tRNA^Lys^, tRNA^Val^, tRNA^His^, tRNA^Asp^ and tRNA^SeC^) also have higher relative levels of tRNA halves even without any ANG overexpression (Fig. S2B), corroborating the existence of tRNA halves in non-stressed cells. Why these tRNAs are more responsive to ANG is later discussed and awaits future investigation.

**Figure 2.**
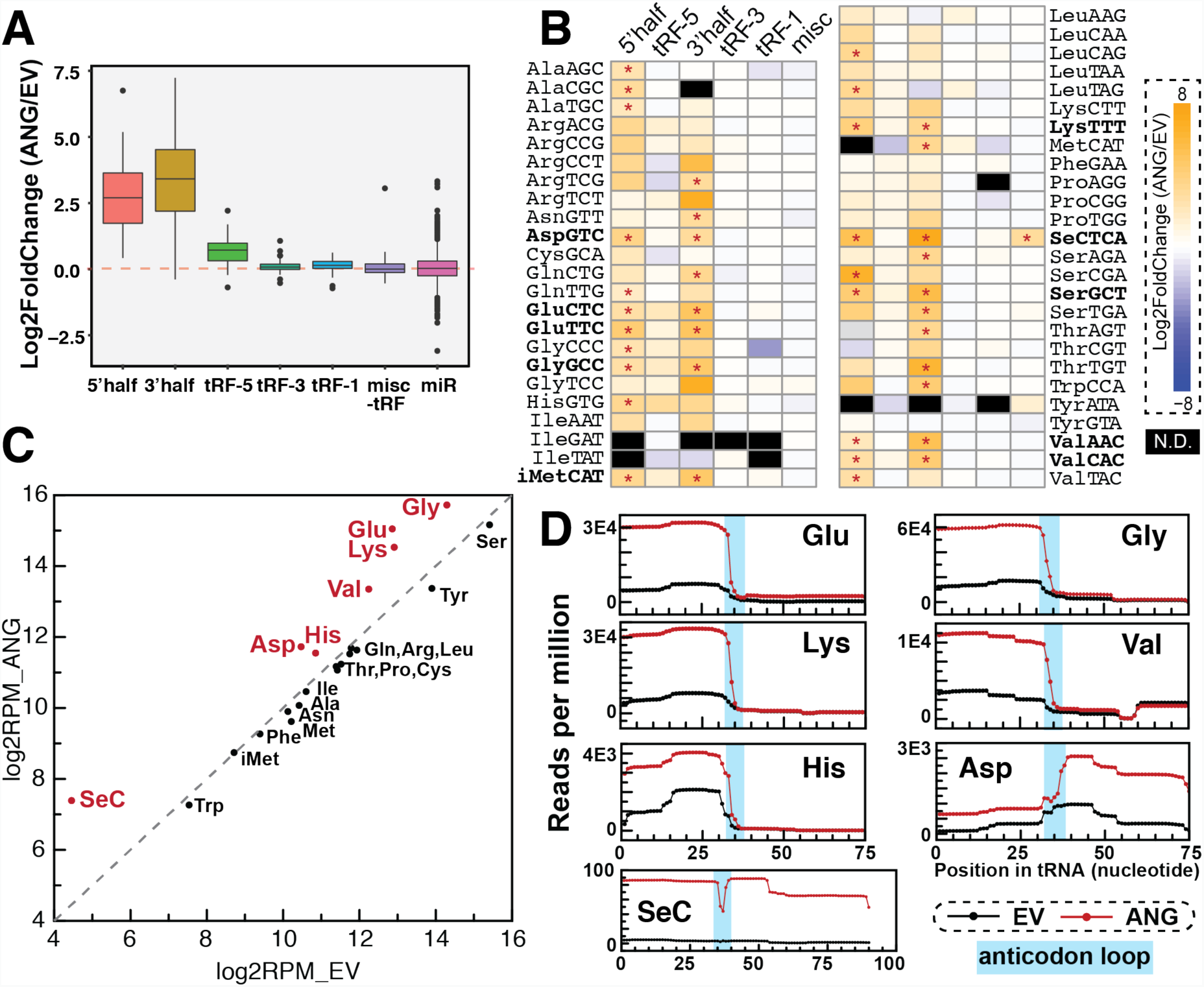
tRNA fragments up-regulated by Angiogenin overexpression. (A-D) tRF changes upon ANG overexpression by small RNA-seq. (A-B) tRNA halves are increased by ANG overexpression. (A) Box-and-whisker plot showing relative fold change of RPM (reads per million mapped reads) on log2 scale of ANG overexpression compared to the empty vector control (averaged from three biological replicates). tRFs are grouped by types and parental tRNAs. Box plot shows median value as middle line, first and third quartile as box outline, and data range (excluding outliers represented by individual dots) as whiskers. (B) Heatmap shows Log2 fold change of tRFs by ANG overexpression. tRFs are grouped by tRF types and parental tRNAs. Significant changes determined by DESeq2 (adjusted p value less than 0.05) are labeled by asterisks. N.D. - not detected. (C-D) ANG overexpression increases tRF reads from certain parental tRNAs, including tRNA^Glu^, tRNA^Gly^, tRNA^Lys^, tRNA^Val^, tRNA^His^, tRNA^Asp^ and tRNA^SeC^. For (C), tRF reads (all types) were summed by parental tRNAs (X-axis: empty vector, Y-axis: ANG overexpression). For (D), coverage plot shows where accumulated tRF reads (excluding tRF-1s) map to the parental mature tRNAs (black line: empty vector control, red line: ANG overexpression). Anticodon loop region for each tRNA (predicted by tRNA covariance model fold in GtRNAdb) is highlighted in cyan.

### Short tRF-3s and tRF-5s are not increased in parallel with the tRNA halves

Short tRFs in the size range compatible with Argonaute loading (17-25 nt) were predicted to play microRNA-like functions due to their direct association with Argonaute proteins by PAR-CLIP (photoactivatable ribonucleoside-enhanced crosslinking and immunoprecipitation) and CLASH (cross-linking, ligation and sequencing of hybrids) experiments (5). Recently we showed three short tRF-3s from tRNA^Leu^ and tRNA^Cys^ can enter RISC (RNA-induced silencing complex) to repress target genes and are generated in a Dicer-independent mechanism (21). To examine whether a tRNA half could be a precursor to a shorter tRF, we plotted 5’ or 3’ fragments by length from the ANG-responsive tRNAs (tRNA^Glu^, tRNA^Gly^, tRNA^Lys^, tRNA^Val^, tRNA^Asp^ and tRNA^His^) and another two tRNAs that produce abundant tRF-3s (tRNA^Ser^ and tRNA^Leu^) (Fig. S2C). For the ANG-responsive tRNAs, the increase of a tRNA half did not lead to an increase of a smaller tRFs from the same tRNA. For example the 23-nt tRF-5s from tRNA^Val^ did not change upon ANG overexpression, despite the significant increase in the corresponding 5’ tRNA half; similarly increase of 3’ half in tRNA^Asp^ did not lead to increase of its 22-nt tRF-3. For the other tRNAs (e.g. tRNA^Ser^ and tRNA^Leu^) that did not produce abundant tRNA halves, shorter tRF-3s were not increased by ANG (Fig. S2C). Together with the above analysis showing global tRF-3 levels unaltered by ANG overexpression (Fig. 1F, Fig. 2A and Fig. S2C), this suggests there is no precursor-product relationship between a tRNA half and a smaller tRF (17-25 nt), which are most likely generated by different biogenesis pathways.

### All tRFs, including tRNA halves, are unchanged in unstressed ANG knock-out cells

To determine whether endogenous ANG is essential for generating tRNA halves, we knocked out the *ANG* gene in HEK293T and U2OS cells by Cas9/CRISPR (clustered regularly interspaced short palindromic repeats) technology. *ANG* (also known as *RNase 5*) and *RNase 4* are transcribed from the same promoter and each protein is translated from one unique exon, therefore we designed guide RNA against the *ANG*-specific exon to deplete ANG specifically (scheme shown in Fig. S3A). Genomic PCR spanning the *ANG*-specific exon shows that we obtained several homozygous clones that have deleted the majority of ANG coding sequence (Fig. 3A), further confirmed by Sanger sequencing (Fig. S3B) and qRT-PCR (Fig. S3C). Immunoblotting by ANG-specific antibody (Fig. S1B) confirmed the complete loss of ANG protein in these clones (Fig. 3B-C). Interestingly, *RNase 4* transcript level is increased 4-6 fold in ANG KO clones (Fig. S3C), without affecting RNase 4 protein (Fig. S3D).

**Figure 3.**
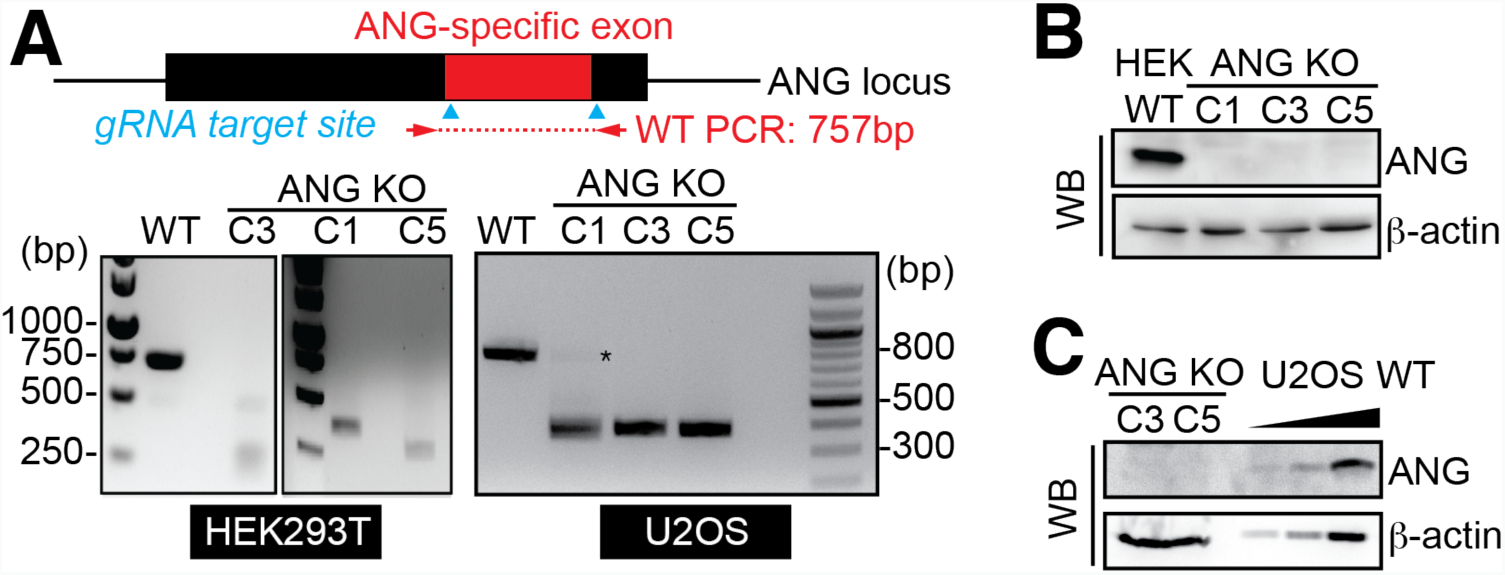
Characterization of Angiogenin knockout cells. (A) PCR of genomic DNA confirms homozygous deletion of majority of ANG coding sequence in HEK293T and U2OS. Guide RNA target sites are shown by cyan arrows, and primer pair (indicated by red arrow) spans the ANG-specific exon (expected amplicon in wild-type is 757bp). U2OS clone 1 was excluded in later experiments due to the residual wild-type band detected by PCR (indicated by asterisk). (B-C) Western blot to confirm loss of ANG protein in the ANG KO clones (B for HEK293T and C for U2OS). (B) 20 μg whole cell lysate for HEK293T WT and KO cells were loaded. (C) 30 μg whole cell lysate for ANG KO clones were compared with a gradient amount (1.5 μg, 6 μg and 30 μg) of wild-type U2OS lysate.

Small RNA profiling of ANG knockout cells suggests there is no global change in any of the six types of tRFs at endogenous levels (without stress) (Fig. 4A). There are also no significant changes in specific tRFs that are consistent between different ANG KO clones (Fig. 4B). In particular, tRF-3 levels are unaltered in ANG KO cells (Fig. 4A and Fig. 4B). We have earlier shown that tRF-3s induced by over-expressing the parental tRNA could specifically repress a target reporter bearing tRF-complementary sequence in the 3’UTR (21). Using this experimental system, we compared the gene silencing activity of three tRF-3s, tRF-3001, 3003 and 3009 (naming based on tRFdb) (49), in both wild-type and ANG knockout cells (Fig. 3C). The over-produced tRF-3s robustly and specifically repressed the luciferase reporter that bears a matching target site (Fig. 3C, left panel), as shown previously (21). Such tRF-3 dependent repression continues in the ANG knockout cells (Fig. 3C, middle and right panels), confirming that functional tRF-3 can be produced in the absence of ANG.

**Figure 4.**
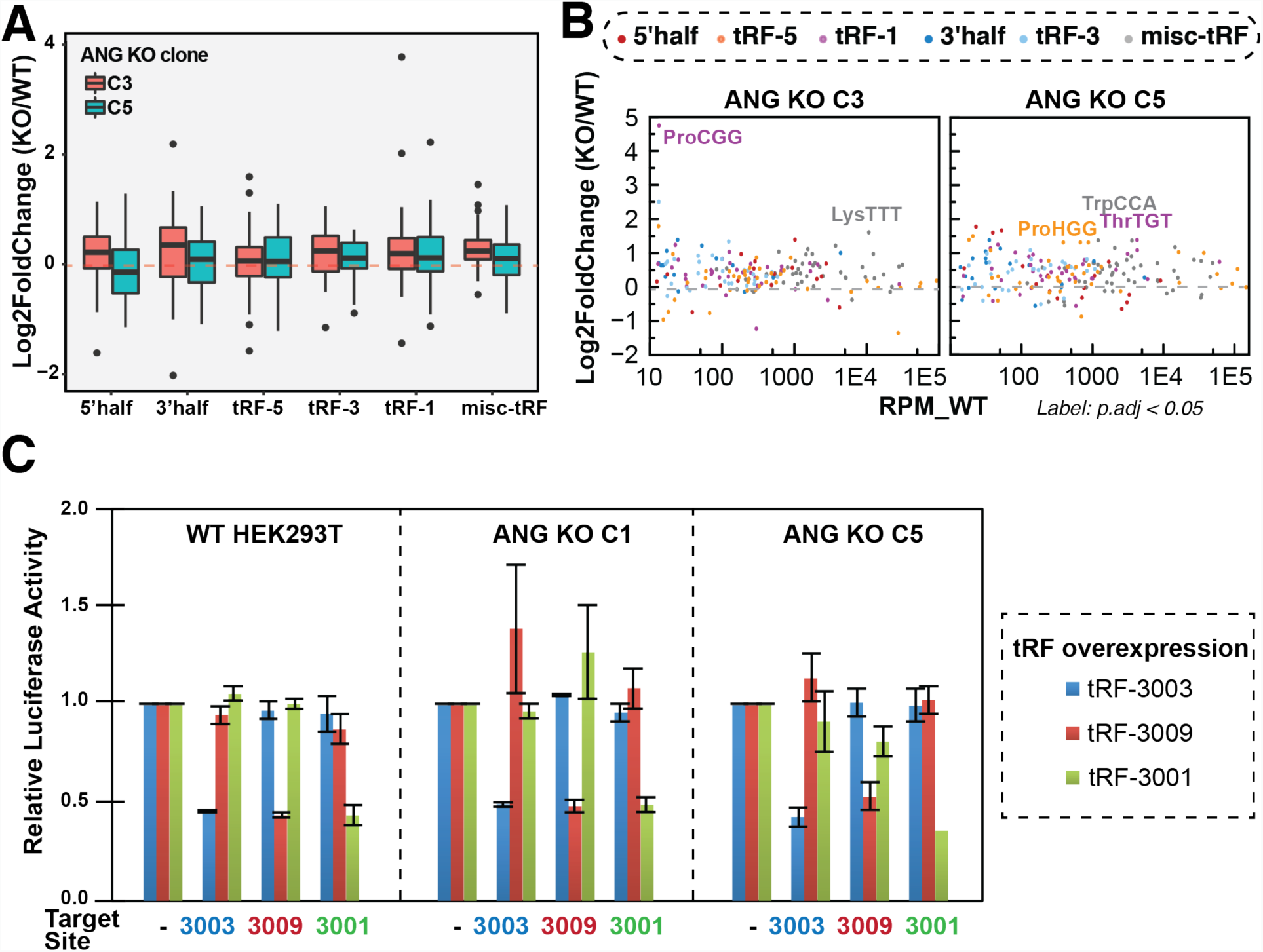
tRF-3s are unchanged in ANG KO cells. (A-B) Changes of tRFs in ANG KO cells without stress by small RNA-seq (A: box-and-whisker plot, B: MA plot). Y-axis shows relative fold change of RPM (reads per million mapped tRF reads) on log2 scale of ANG knockout cells compared to the U2OS wildtype control (averaged from two biological replicates).(B) Significant changes determined by DESeq2 (adjusted p value less than 0.05) are labeled. (C) Gene silencing activity of three tRF-3s (tRF-3003, 3009 and 3001) were measured by dual-luciferase assay in wild-type HEK293T cells and ANG KO cells (two clones C1 and C5). tRFs were over-expressed by corresponding parental tRNAs and relative repression was observed when 3’UTR of renilla luciferase gene has a paired complementary tRF target site (relative to no target site luciferase gene). Error bars represent standard deviation from three biological replicates.

### ANG is dispensable for translation and proliferation arrest induced by high concentration arsenite stress

ANG has been proposed to be the enzyme responsible for the generation of tRNA halves in stressed human cells (16,17). Similar to previous studies, we stressed U2OS cells with high concentration (1 mM) of sodium arsenite (SA) for 1 hour, which has been shown to induce abundant tRNA halves and inhibit translation (17,19,41,44). Indeed the sodium arsenite stress induced tRNA cleavage and produced tRNA halves that can be detected by Sybr gold (Fig. 5A). In addition, high concentration (0.2 mM or 1 mM) of sodium arsenite treatment leads to complete protein translational arrest in both wild-type and ANG KO cells as measured by puromycin pulse labeling (Fig. 5B and Fig. S4B), suggesting ANG is dispensable for the translational arrest induced by high concentration of arsenite. Consistent with previous observations, wild-type cells did not show much translational repression at a lower concentration (50 μM) of arsenite for 1 hour (Fig. S4C) (41). Although, ANG KO cells displayed mild translational repression at 50 μM of arsenite (Fig. S4C), suggesting ANG KO cells might be more sensitive to low concentration of arsenite, the cell growth arrest at 24 hours was similar in wild-type and ANG KO cells (Fig. 5C) regardless of arsenite concentrations (Fig. S4C). Thus, ANG is not required for stress-induced growth arrest.

**Figure 5.**
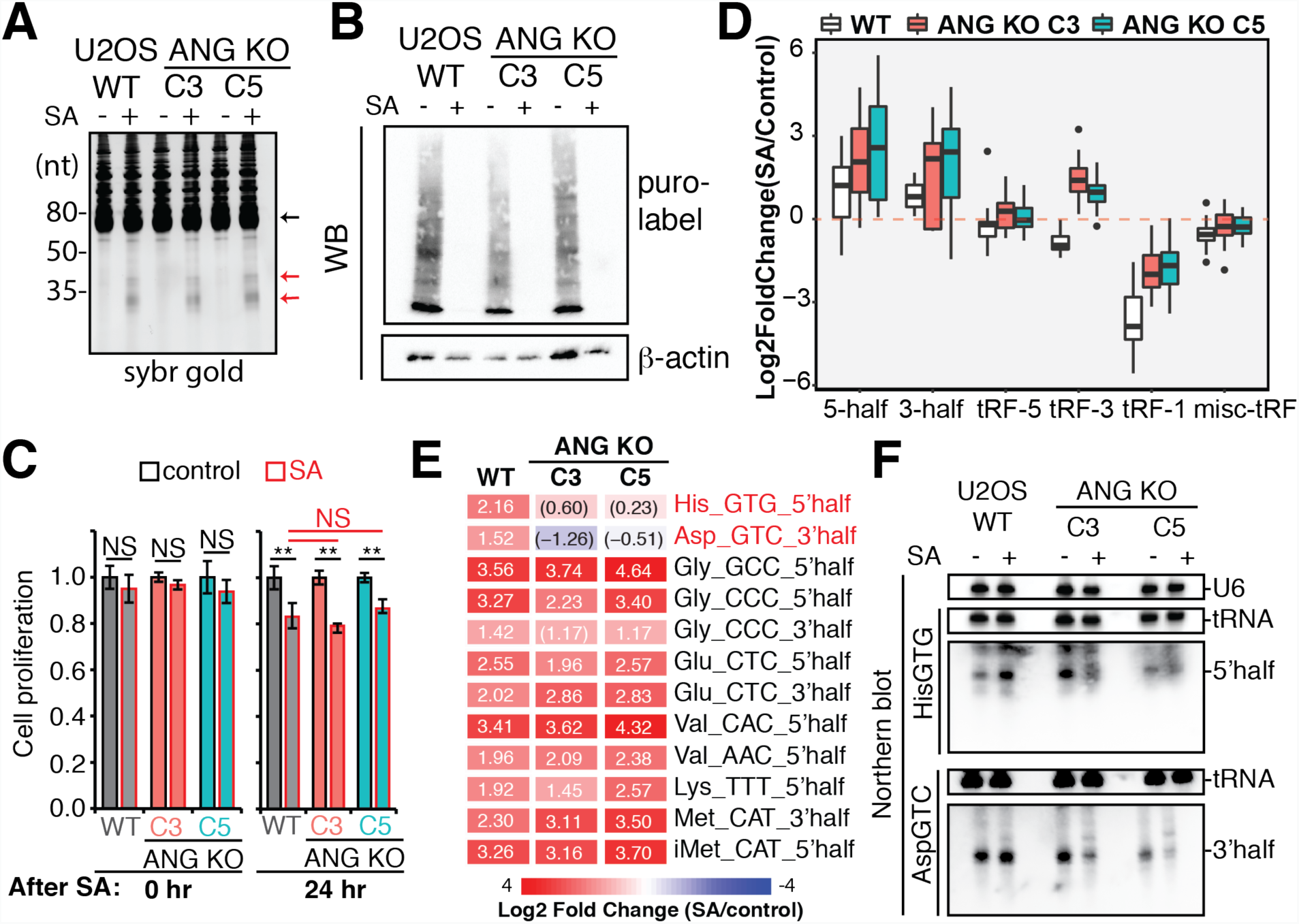
ANG-dependent and ANG-independent stress response. (A-F) Wild-type and ANG KO U2OS cells were treated with 1 mM sodium arsenite for 1 hour to identify ANG-dependent stress response. (A) Sybr gold staining of 5 μg total RNA detected arsenite-induced tRNA half bands (indicated by red arrows) in both wild-type and ANG KO U2OS cells (black arrow points to tRNA band). (B) Puromycin pulse labeling was used to detect translation arrest. (C) Cell proliferation immediately (0 hr) and 24 hour after SA treatment was measured by MTT assay and normalized to wild-type control cells. Error bars represent standard deviation from three biological replicates. Student t-test was performed between control and SA for each condition, or between WT and ANG KO (**p < 0.05). (D-E) Small RNA-seq was used to quantify stress-responsive tRFs (D: box-and-whisker plot, E: heatmap). (D) Y-axis shows relative fold change of stress versus control on log2 scale in wild-type and ANG knockout cells (two clones C3 and C5). (E) Arsenite-induced tRNA half changes in WT and ANG KO cells. Only significant changes in WT are shown here (determined by DESeq2, adjusted p value less than 0.05). Numbers inside each box shows Log2 Fold Change (parenthesis indicates p.adj > 0.05, otherwise p.adj ≤ 0.05). For a more complete list of tRNA halves, refer to Table S3. (F) Northern blot of ANG-dependent stress-induced tRNA halves, 5’ tRNA^HisGTG^ half and 3’ tRNA^AspGTC^ half, in wild-type and ANG KO U2OS cells.

### ANG is required for generating specific tRNA halves induced by high concentration arsenite

Upon ANG over-expression we observed significant and specific increase of both 5’ and 3’ tRNA halves from multiple isoacceptors (Fig. 1 and Fig. 2). However, to our surprise, the stress-induced tRNA half bands did not diminish in ANG KO cells (Fig. 5A). Thus, although ANG can cleave tRNAs in the anticodon loop, most of the tRNA cleavage seen after high concentration arsenite induced stress is ANG-independent.

To identify whether any stress-induced tRNA halves were dependent on ANG, small RNA sequencing was carried out in WT and ANG KO cells with and without sodium arsenite stress. To avoid normalization artifacts from arsenite-induced changes in other types of small RNAs (such as rRNA fragments and miRs), total reads mapped to tRNAs (all tRFs) were used for normalization. Small RNA-seq revealed a global up-regulation of 5’ halves and 3’ halves by SA stress in wild-type cells (Fig. 5D and Fig. S5A), consistent with our Sybr Gold staining (Fig. 5A) and previous publications (17,19,41,44). Such induction remained the same in ANG KO cells (Fig. 5D). Consistent with the sybr gold staining (Fig. 5A) and global trend by small RNA-seq (Fig. 5D), differential analysis reveals that most stress-induced tRNA halves (such as from tRNA^Glu^, tRNA^Gly^, tRNA^Lys^ and tRNA^Val^) are still up-regulated to similar degree in ANG KO cells (Fig. 5E, Table S3). However, there are specific tRNA halves (mainly 5’ half from tRNA^HisGTG^ and 3’ half from tRNA^AspGTC^) that are significantly induced by SA stress in WT but not in ANG KO cells (Fig. 5E, Table S3). The role of ANG in producing the 5’ half of tRNA^HisGTG^ and 3’ half of tRNA^AspGTC^ were confirmed by Northern blot, which also shows that the levels of the parental tRNAs are not changed by arsenite or by ANG KO (Fig. 5F). Interestingly, these fragments were decreased by arsenite treatment in ANG KO cells, suggesting unknown tRNA half destabilizing/degradation pathway(s) might be more active in the ANG KO cells upon stress. Meanwhile, Northern blot for 5’ tRNA halves from tRNA^GluCTC^ and tRNA^GlyGCC^ still shows similar levels of induction by high concentration arsenite in ANG KO cells (Fig. S5C). Altogether our analysis reveals that specific stress-induced tRNA halves are altered in ANG KO cells, and the majority of stress-induced tRNA halves can be generated by ANG-independent pathways.

Interestingly, short tRF-3s were slightly decreased by arsenite stress in wild-type cells, but were increased globally by arsenite stress in ANG KO cells (Fig. 5D and Fig. S5A). This suggests the ANG may have a role in regulating the levels of tRF-3s following stress. As a side-note, tRF-1s are globally down-regulated 8-16-fold by arsenite stress, which is not ANG-dependent (Fig. 5D). The mechanism for this decrease after stress is unclear.

## Discussion

In this study, we profiled tRNA cleavage pattern after over-expression or knock-out of human ANG (angiogenin, RNase 5). The results suggest ANG can induce but is not required for tRNA cleavage to produce tRNA halves. It has been known that ANG is specific towards tRNA while it has rather low ribonuclease activity towards other RNAs *in vitro* (50), and such specificity was explained by ANG’s unique structural features (51,52). However, it was not entirely clear whether ANG cleaves all tRNAs in cells equally well. Here we show by over-expressing human ANG that its cleavage has specificity towards certain tRNAs, including tRNA^Glu^, tRNA^Gly^, tRNA^Lys^, tRNA^Val^, tRNA^His^, tRNA^Asp^ and tRNA^SeC^ (Fig. 1 and Fig. 2). This is consistent with a previous report showing some specificity by recombinant ANG towards certain tRNAs by a customized tRF microarray approach (44) and another report showing that ANG does not produce halves from tRNA^Tyr^ (16). This apparent preference could come from a combination of (a) ANG’s substrate specificity, which could be affected by base sequence (53) and tRNA modifications (54–57) or (b) higher stability of these tRNA halves. 5’ halves from tRNA^Gly^ and tRNA^Glu^ have recently been shown to have higher stabilities due to dimerization (58). It is also interesting that cloning frequency of the 5’ half is higher than the corresponding 3’ half for most tRNAs except tRNA^Asp^, suggesting that either the stability or the clonability of the two halves are different. This might be due to the 3’ attachment of amino acids on 3’ halves that prevent their cloning (42). The effect of all these factors on tRNA half induction by ANG (and other RNases) needs further mechanistic investigation.

tRNA regulation is important in stress biology (59,60) and stress-induced tRNA cleavage seems to be a conserved feature. Oxidative stress (17,61), hyperosmotic stress (44), nutrient deprivation, ER stress (62), virus infection (63,64) and other types of stress increase the levels of tRNA halves. Stress-induced tRNA halves have been suggested to be signaling molecules, rather than by-products to reduce tRNA levels (65). Indeed, tRNA halves have been shown to play various roles in stress pathways. For example, 5’ halves from tRNA^Ala^ and tRNA^Cys^ could interact with YBX-1 to inhibit protein translation and promote stress granule formation independent of eIF2 phosphorylation (17–19). Interestingly, these two 5’ tRNA halves can form stable intermolecular G-quadruplex structure (66) and yet neither of them was induced by sodium arsenite in our experiment. In addition, tRNA halves (especially 3’ halves) from various tRNAs interact with cytochrome C and protect cells from apoptosis during stress (20). Recently stress-induced 3’ half from tRNA^Thr^ in *Trypanosoma Brucei* has been found to stimulate translation (67). It is highly likely that tRNA halves have divergent functions and many tRNA halves (including the ones we identify from selenocystein) have not been investigated in details.

Our results suggest ANG is not the only RNase that mediates stress-induced tRNA cleavage. This is not surprising since RNase A family is vertebrate-specific, yet stress-induced tRNA cleavage is a well-conserved phenomenon in organisms that do not have RNase A. For example, In *Tetrahymena*, *S. cerevisiae* and *Arabidopsis*, stress-induced generation of tRNA halves has been attributed to the RNase T2 family (12,68,69). In mammalian cells, other RNases have also been described to cleave tRNAs, such as Dicer (70–72), RNase 1 (36), RNase L (73) and Schlafen13/SFLN13 (74). RNase L was suggested to act independently from ANG by cleaving anticodon loop of tRNA^Pro^, tRNA^His^ and tRNA^Gln^ (73). Whether different RNases are regulated by different stresses and whether one RNase could have different specificity in different cell types still needs to be determined.

Increasing evidence suggests tRFs and tRNA halves exist in conditions beyond stress. In particular, 3’ tRNA fragments that are 17-25 bases long, tRF-3s, have been detected at high abundance, enter into Argonaute complexes and implicated in important biological functions including gene silencing, retrotransposon silencing and ribosome biogenesis (5,21,23,24). These tRF-3s are clearly not regulated by ANG in unstressed cells from our analysis; further investigation is required to identify factors involved in the biogenesis of these tRF-3s. The results also argue against the tRNA halves serving as precursors for the biogenesis of the shorter tRFs that enter into Argonaute complexes.

ANG over-expression specifically induces tRNA halves from tRNA^Glu^, tRNA^Gly^, tRNA^Lys^, tRNA^Val^, tRNA^His^, tRNA^Asp^ and tRNA^SeC^ (Fig. 2); while ANG knock-out in stressed cells decreases tRNA halves from tRNA^His^ and tRNA^Asp^ (Fig. 5). Recently, tRNA halves including both halves of tRNA^HisGTG^ and tRNA^AspGTC^ (named as SHOT-RNAs, Sex Hormone-dependent tRNA-derived RNAs) were found in estrogen or androgen receptor positive cells and the 5’ halves enhance cell proliferation (42). Notably, levels of two SHOT-RNAs (5’ half of tRNA^HisGTG^ and 5’ half of tRNA^AspGTC^) were decreased by ANG knock-down in BT-474 breast cancer cells (42), suggesting ANG’s role in non-stressed sex-hormone receptor positive cells. More recently, 5’ tRNA halves were found to be up-regulated during stem cell differentiation and appeared to be ANG-independent (75). Abundant tRNA halves have also been detected in body fluids (e.g. blood/serum, sperm, cerebrospinal fluid, urine) and exosomes (76–82). Intriguingly, 5’ halves from tRNA^Gly^, tRNA^Glu^ and tRNA^Val^ in mouse sperm are responsive to paternal diet change and mediate transgenerational change to the offsprings (83,84). Whether ANG or another RNase is involved in the generation of these tRNA halves in body fluids and exosomes is unclear.

Another distinct group of tRFs is tRF-5s of 26-30 nt, also called td-piRNAs, that interact with PIWI proteins (28–30). td-piRNA from tRNA^Glu^ is highly expressed in human monocytes and could be down-regulated by IL-4; this td-piRNA complexed with PIWIL4 plays a role in epigenetic regulation to inhibit *CD1A* transcription (28). This td-piRNA and the corresponding tRNA half are both increased by ANG overexpression (Fig. S2C). Because of the concurrent increase in specific tRNA halves and the tRF-5s of 26-30 nt from the same tRNAs, our results suggest td-piRNAs might be derived from tRNA halves, as recently shown for two td-piRNAs in *Bombyx mori* germ cells (29). Future studies about tRF and tRNA half biogenesis and function would bring new insights into this new category of non-coding RNAs.

### Experimental procedures

#### Cell culture, treatment and proliferation assay

HEK293T and U2OS cells were obtained from ATCC and maintained in HyClone Dulbecco’s High Glucose Modified Eagles Medium (GE #SH30081.01) with 10% FBS. For sodium arsenite treatment, U2OS cells were grown to ~80% confluency and replaced with fresh media, supplemented with 0.05 - 1 mM (final concentration in water) sodium arsenite (Sigma #S7400) in CO_2_ incubator at 37 °C for 1 hour. Cell proliferation was measured with CellTiter 96 Non-Radioactive Cell Proliferation Assay kit (Promega #G4100) at 590-nanometer absorbance on plate reader (BioTek Synergy HT) according to manufacturer’s instructions.

#### Plasmid construction and transfection

Human ANGIOGENIN cDNA sequence (corresponding to 147 amino acids, Uniprot #P03950) was PCR amplified from CCSB human ORFeome collection v5.1 (Forward primer: ATGGTGATGGGCCTGGGCGTT, Reverse primer: TTACGGACGACGGAAAATTGACTG) and cloned into pcDNA3.1 (digested by BamHI and XhoI) by In-Fusion HD cloning plus (Takara #638910). Transient over-expression of pcDNA3.1-ANG and control empty plasmid was performed with Lipofectamine 2000 (Thermo Fisher #11668027) for 72 hours accordingly to manufacturer’s instructions. tRNA overexpression and luciferase reporter assay in HEK293T cells was performed as previously (21).

#### CRISPR knock-out cell generation and confirmation

CRISPR knock-out was generated following previous protocol with slight modifications (85). Wild-type U2OS and HEK293T cells were plated in 6-well plate at ~80% confluency and co-transfected with human codon-optimized Cas9 plasmid (Addgene #41815), two guide RNA (gRNA) containing plasmids, and puromycin selection plasmid pPUR. Two gRNAs were cloned separately into pCR-BluntII-TOPO vector (Addgene #41820). We initially designed 6 gRNA target sequence and screened in HEK293T for the best knockout efficiency on population of cells, based on the screening we used the combination of guide RNA target sequence CCCACCGACCCTGGCTCAGGATA (gRNA2, located early in the coding exon) and CCGTCGTCCGTAACCAGCGGGCC (gRNA4, located immediately after stop codon). Transfection was performed with Lipofectamine 2000 (Thermo Fisher #11668027) for HEK293T and Lipofectamine LTX plus (Thermo Fisher #15338100) for U2OS according to manufacturer’s instructions. 24 hours post transfection, cells were expanded into 10-cm dish and selected with 1 μg/ml puromycin up to when control (non-transfected) cells die. Single cell cloning was subsequently performed on puromycin-selected cells and checked by genomic DNA PCR (QuickExtract DNA Extraction Solution, Epicentre #QE09050; OneTaq polymerase, NEB #M0482) for successfully deleted genomic region (primers for checking ANG genomic deletion: Forward-ATTTGGTGATGCTGTTCTTGGGTCT, Reverse-AGGGAGCAGCCAAGTAGAGAAAATG).

Clones with successfully genomic deletion as determined by a lower band after PCR and Sanger sequencing will then be checked by western blot to confirm loss of protein.

#### RNA extraction and Northern blot

Cells were grown to ~80% confluency and washed briefly twice with ice-cold PBS. Total RNA was extracted by Trizol reagent (Invitrogen #15596018) and purified with column-based Direct-zol RNA Miniprep kit (Zymo Research #R2051). For Sybr gold detection, 5 μg total RNA was separated on 15% Novex Urea-TBE gel (Invitrogen #EC6885BOX) and stained with Sybr Gold (Invitrogen #S11494). For non-radiolabeled Northern blot analysis, 5 - 10 μg total RNA was separated on 15% Novex Urea-TBE gel and transferred to Amersham Hybond-N+ membrane (GE #RNP203B) by Trans-Blot SD semi-dry transfer apparatus (Biorad). After the membrane was cross-linked with Stratalinker (Stratagene) at 254 nanometer wavelength for 12000 μJ and further baked at 70 °C for at least 30 minutes, the membrane was blocked with ExpressHyb hybridization solution (Clontech #636831) at 42 °C for 30 minutes. After incubation with 10-50 pmole/ml biotinylated DNA probe at 42 °C overnight, the membrane was washed three times at room temperature with 1X SSC, 0.1% SDS and then detected with Chemiluminescent Nucleic Acid Detection Module Kit (Thermo Fisher #89880) on G:BOX imaging system (Syngene). Northern probe sequences are U6 (BIO-TGCGTGTCATCCTTGCGCAG), 5’ tRNA^HisGTG^ (BIO-CAGAGTACTAACCACTATACGATCAC GGC), 3’ tRNA^AspGTC^ (BIO-GTCGGGGAATCG AACCCCGGTC), 5’ tRNA^GluCTC^ (BIO-CCTAA CCACTAGACCACCAGGGA) and 5’ tRNA^GlyGCC^ (BIO-TCTACCACTGAACCACCCATGC).

#### Puromycin pulse labeling and western blot

U2OS cells were grown to ~ 80% confluency and replaced with fresh media, supplemented with 10 μg/ml (final concentration in water) puromycin (Sigma #P8833) in CO_2_ incubator at 37 °C for 10 minutes. Cells are washed with ice-cold PBS twice and lysed in RIPA buffer supplemented with Halt protease inhibitor cocktail (Thermo Scientific # 78438). For western blot analysis, 1.5 – 30 μg (determined by Bradford assay) whole cell lysates were separated on 12% SDS-PAGE and transferred to 0.45 μm Amersham Protran nitrocellulose membrane (GE #10600002) by Trans-Blot SD semi-dry transfer apparatus (Biorad). These primary antibodies and corresponding dilutions were used in 3% milk in PBST: anti-angiogenin (Santa Cruz #sc-74528 C1, mouse monoclonal, used at 1:100), anti-β-actin (Santa Cruz #sc-47778 C4, mouse monoclonal, used at 1:2000), anti-RNase4 (Abcam #ab200717, rabbit polyclonal, used at 1:500) and anti-puromycin (Millipore #12D10, mouse monoclonal, used at 1:10,000). Anti-mouse HRP-linked secondary antibody (Cell Signaling #7076) or anti-rabbit secondary antibody (Cell Signaling #7074) was used at 1:5000 dilution. Chemiluminescence detection was done with Immobilon HRP substrate (Millipore #WBKLS0500) on G:BOX imaging system (Syngene).

#### Small RNA-sequencing library preparation

Small RNA-sequencing libraries were prepared following NEBNext Small RNA library kit (NEB #E7300). Briefly, 1 μg total RNA was ligated with 3’ pre-adenylated adaptors and then 5’ adaptors. After reverse transcription and PCR amplification with indexed-adaptors, each library was size selected on 8% TBE-PAGE gel (Invitrogen #EC6215BOX) to enrich for 15-50 nt insert. A brief workflow and a representative gel before size selection are shown in Fig. S1C and S1D. Individual gel purified libraries were pooled and sequenced on Illumina NextSeq 500 sequencer with mid-output or high-output, 75-cycles single-end mode at Genome Analysis and Technology Core (GATC) of University of Virginia, School of Medicine. All small RNA-sequencing data has been deposited into GEO (GSE130764).

#### Small RNA sequencing data analysis

For small RNA-seq analysis, cutadapt v1.15 (Python 2.7.5) was used to trim 3’ adaptor sequence at most 10% errors and remove reads less than 15 nt (cutadapt --nextseq-trim=20 -a AGATCGGAAGAGCACACGTCTGAACTCCA GTCAC -m 15). The reads were then mapped to human genome (gencode GRCh38.p10 Release 27, primary assembly) by STAR aligner v2.5.4 with setting based on previous paper to allow multi-mapping (86), and the total number of mapped reads were used for normalization. In general, mapped percentage is more than 95% (summarized in Table S1). To quantify microRNA and tRNA fragments abundance, unitas v.1.7.3 (87) (with SeqMap v1.0.13) (88) was used with setting – species_miR_only –species homo_sapiens to map the reads to human sequence of miRBase Release 22 (89), genomic tRNA database (90), Ensembl Release 95 and SILVA rRNA database Release 132 (with addition of U13369.1 Human ribosomal DNA complete repeating unit). This setting (equivalent to –tail 2 –intmod 1 –mismatch 1 – insdel 0) will allow 2 non-templated 3’ nucleotides and 1 internal mismatch for miRNA mapping; 1 mismatch and 0 insertion/deletion for tRNA fragments mapping. Overall tRF-3 counts were summed from tRF-3 and tRF-3CCA to count all tRF-3 reads irrespective of 3’CCA addition. tRF reads mapped to specific tRNA are grouped by tRF types (illustrated in Fig. 1A); in the case of multi-mapping, a read is counted as fraction to avoid duplicate counts. For a secondary tRNA fragment mapping method, MINTmap (91) was used to map reads to tRNA without any mismatch. For both unitas and MINTmap, cutoff for tRNA halves (both 5’ and 3’ halves) is 30 nucleotide long. For differential analysis, DESeq2 (48) was used on count matrix of tRFs and miRs.

## Acknowledgements

We would like to thank Dr. Pavel Ivanov and Dr. Paul Anderson (Harvard University) for their technical assistance on stress induction of tRNA halves, Dr. David Rosenkranz (Johannes Gutenberg University) for help with unitas and Dr. Pankaj Kumar for bioinformatics training. We thank all Dutta lab members for helpful discussions, Dr. Yongde Bao at the University of Virginia GATC (genome analysis and technology core) and the Egelman lab (University of Virginia) for sharing equipment.

## Author contributions

ZS and AD designed the experiments. ZS, MK and ES generated CRISPR knockout cells. CK performed luciferase reporter assay. ZS performed all the other experiments and analysis.

## FOOTNOTES

This work is supported by NIH R01 AR067712, University of Virginia Brain Institute Seed Funding and University of Virginia Supporting Transformative Autism Research (STAR) Pilot Award.

The content is solely the responsibility of the authors and does not represent the official views of the National Institutes of Health.

## Abbreviations

ANG: Angiogenin
RPM: reads per million
tRF: tRNA-derived RNA fragment
td-piRNA: tRNA-derived PIWI-interacting RNA
miR: microRNA
SeC: selenocystein
SA: sodium arsenite
KO: knock-out

