## Supplemental Figures and Tables for "The role of ANGIOGENIN, or lack thereof, in the generation of stress-induced tRNA halves and of smaller tRNA fragments that enter Argonaute complexes"

**Figure S1. Set-up of ANG over-expression and small RNA-sequencing.**

**Figure S2. tRF changes upon ANG over-expression.**

**Figure S3. Generation of ANG knock-out cells.**

**Figure S4. U2OS wild-type and ANG KO cells under different concentrations of arsenite stress.**

**Figure S5. Stress-induced tRNA halves in ANG knock-out cells.**

**Table S1. Statistics of small RNA-sequencing samples.**

**Table S2. DESeq2 differential analysis of tRFs and miRs by HEK293T ANG over-expression (3 biological replicates).**

**Table S3. DESeq2 differential analysis of tRNA halves by SA stress in U2OS wild-type and ANG KO cells (2 biological replicates).**

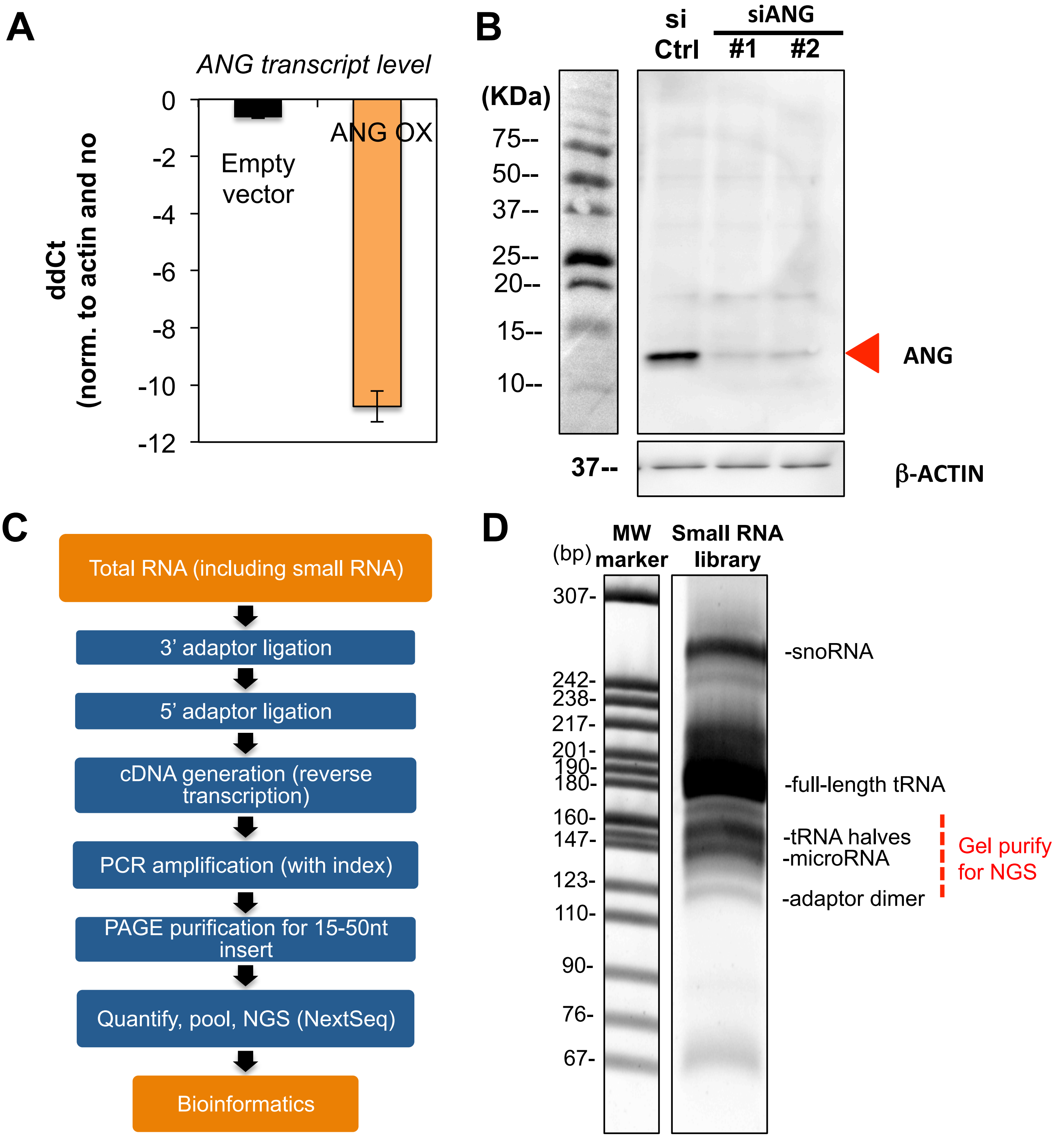


**Figure S1. Set-up of ANG over-expression and small RNA-sequencing.**
(A) qPCR to quantify ANG transcript after ANG over-expression compared to no transfection or empty vector controls. Delta delta Ct values are normalized to β-actin and no transfection control.
(B) Validation of ANG antibody specificity after siANG (#1: UUUCACAAGCAACAACAACGU; #2: AAAGAAGACUUGCUUAUUCUU) in western blot.
(C) Workflow of small RNA-sequencing library preparation.
(D) An example of small RNA-seq library before PAGE purification and the fragment size between 15-50 nt (corresponding to the red dashed area) were selected on PAGE gel.

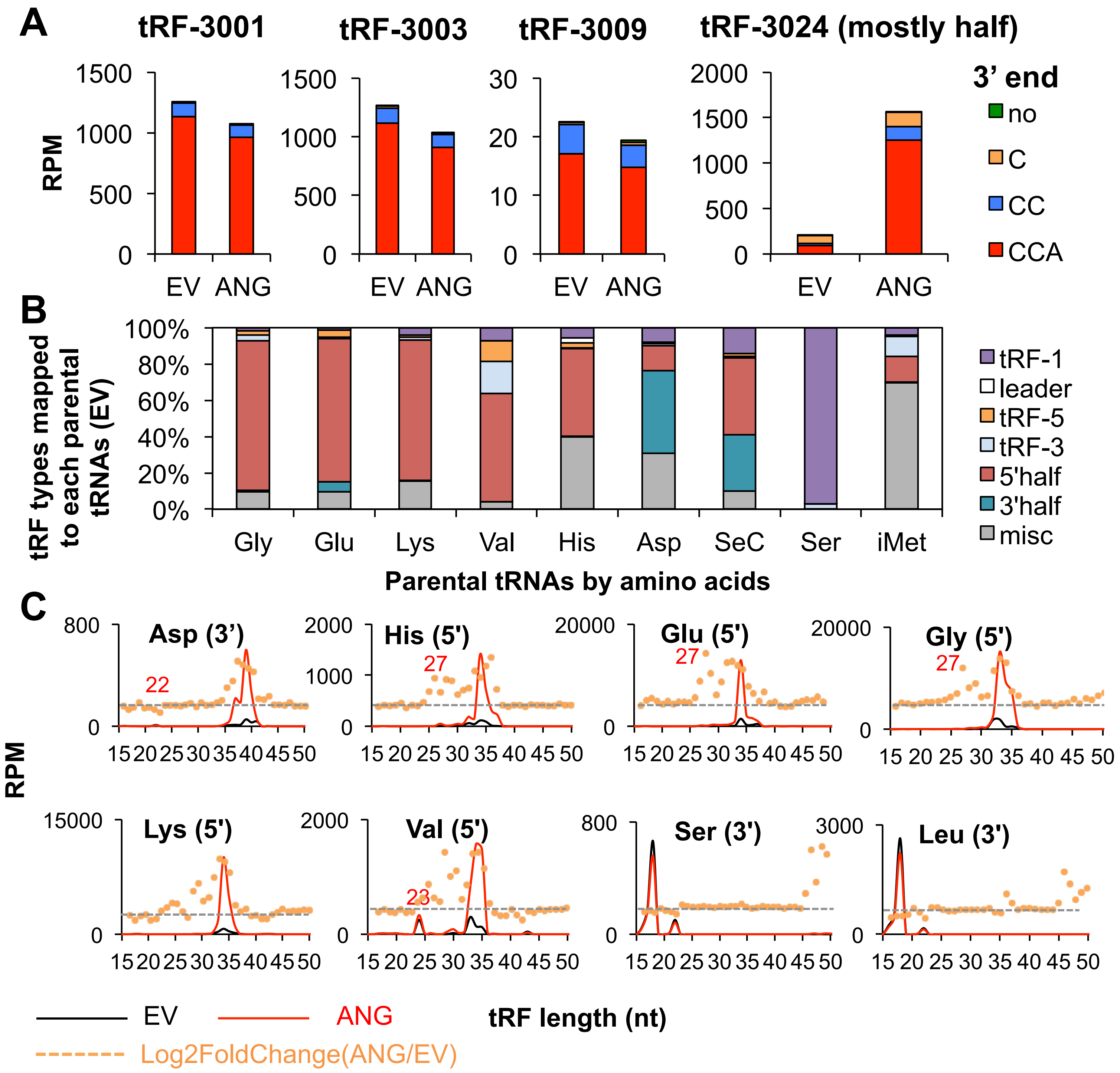


**Figure S2. tRF changes upon ANG over-expression.**

(A) Most tRF-3s contain an intact CCA 3' end. tRF-3001, tRF-3003, tRF-3009 and tRF-3024 (naming based on tRFdb) were checked for their 3'end sequence.

(B) Relative percentage of tRF types for each specific parental tRNA group was shown in the control (EV: empty vector) cells averaged from three biological replicates.

(C) tRF length (X-axis: nucleotides/nt) for each indicated group of tRFs were compared between control (empty vector, in black line) and ANG over-expression (in red line) averaged from three biological replicates. Y-axis shows RPM (reads per million mapped reads). Orange dot shows Log2FoldChange of ANG overexpression over control (gray dashed line indicates Log2FoldChange=0).

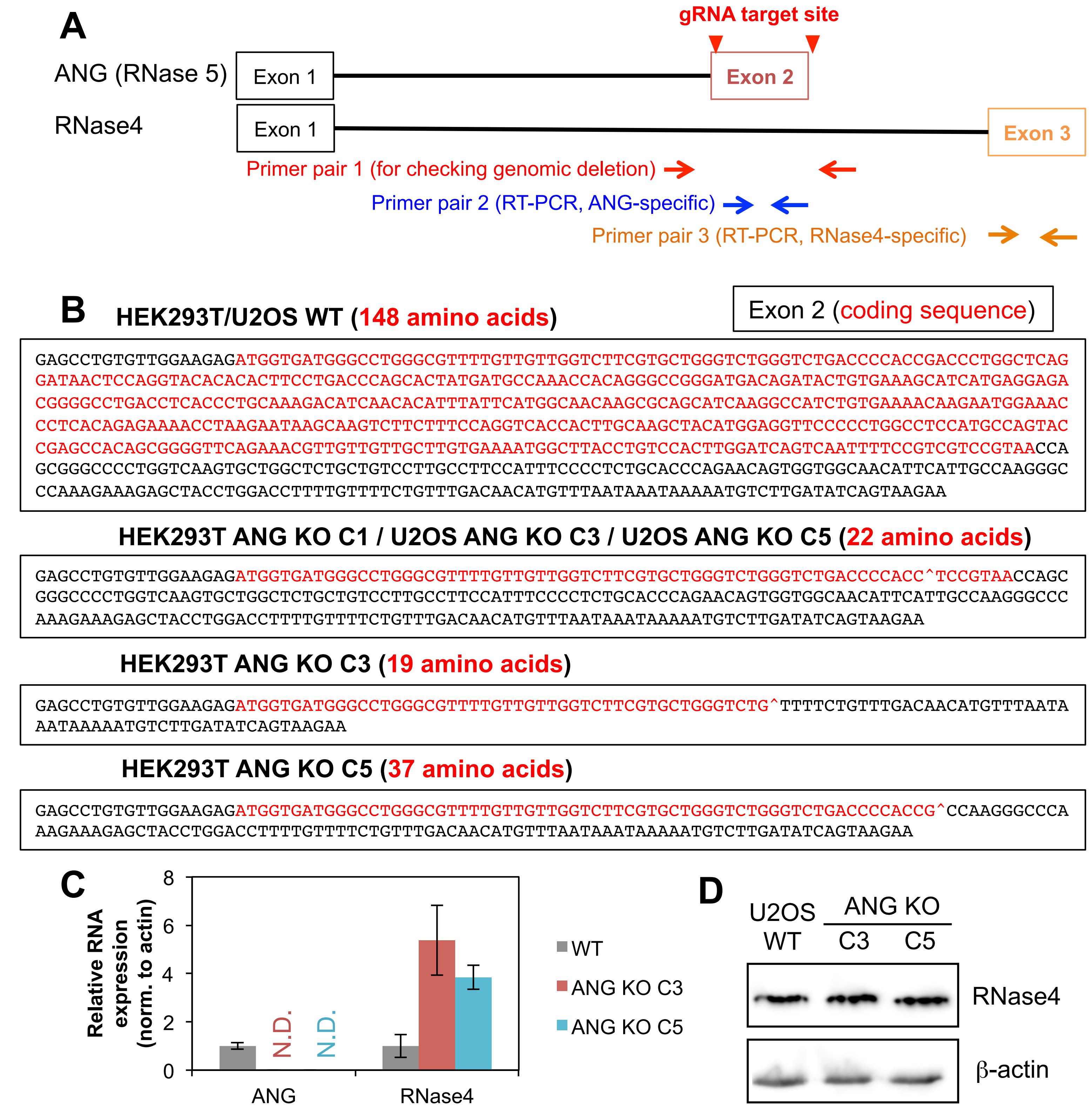


**Figure S3. Generation of ANG knock-out cells.**

(A) Scheme of design of *ANG*-targeting gRNAs and genomic locations of primer pairs. Primer pair 1 was used to check genomic deletion of *ANG*, primer pair 2 was used in qRT-PCR for *ANG*, and primer pair 3 was used in qRT-PCR for *RNase 4*.

(B) Sanger sequencing of genomic sequence in wild-type and ANG knock-out cells (^ indicates site of deletion).

(C) qRT-PCR to detect *ANG* and *RNase4* RNA level in U2OS WT and ANG KO cells (N.D. = not detected, error bars represent standard errors from three technical replicates). Relative expression was calculated by $2^{-ddCt}$ (normalized to β-actin and WT).

(D) Western blot to detect RNase4 protein level in U2OS WT and ANG KO cells.

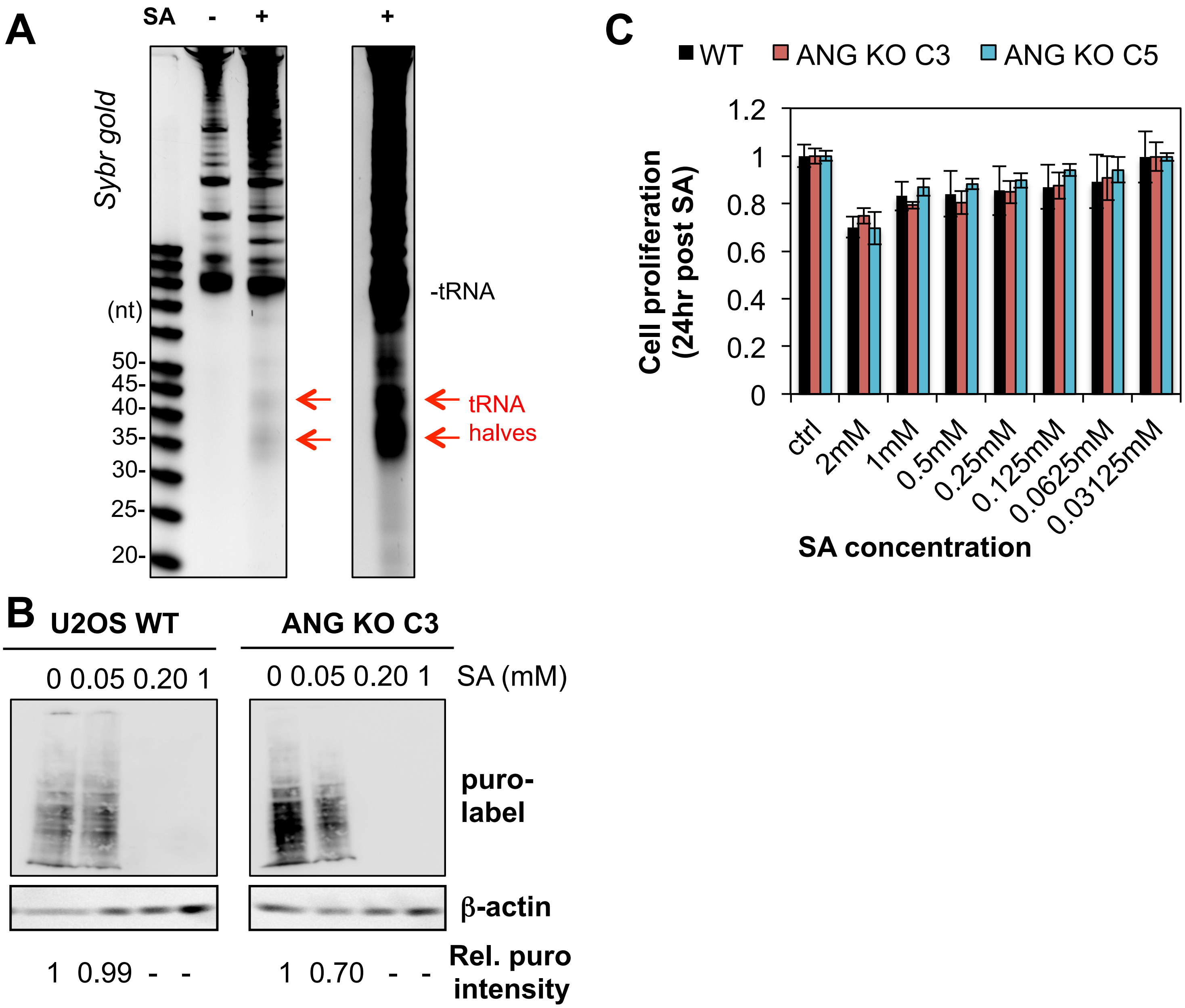


**Figure S4. U2OS wild-type and ANG KO cells under different concentrations of arsenite stress.**

(A) tRNA halves were detected by sybr gold of total RNAs after 1 mM sodium arsenite treatment for 1 hour in U2OS cells.

(B) U2OS wild-type and ANG KO cells were treated with different concentrations (0, 0.05 mM, 0.2 mM or 1 mM) of sodium arsenite for 1 hour. Protein translation was monitored by puromycin labeling for 10 minutes before cell lysis. β-actin was used as a loading control. Signal intensity was measured in each lane by ImageJ and subtracted by background. Relative ratio between anti-puromycin and anti-actin signal was calculated and normalized to the no arsenite treatment control.

(C) U2OS wild-type and ANG KO cells were treated with different concentrations (from 31.25 μM to 2 mM) of sodium arsenite for 1 hour. 24 hours post treatment, cell proliferation/metabolism was measured by MTT assay. Absorbance values were normalized to control WT cells, error bars represent standard deviation from triplicates.

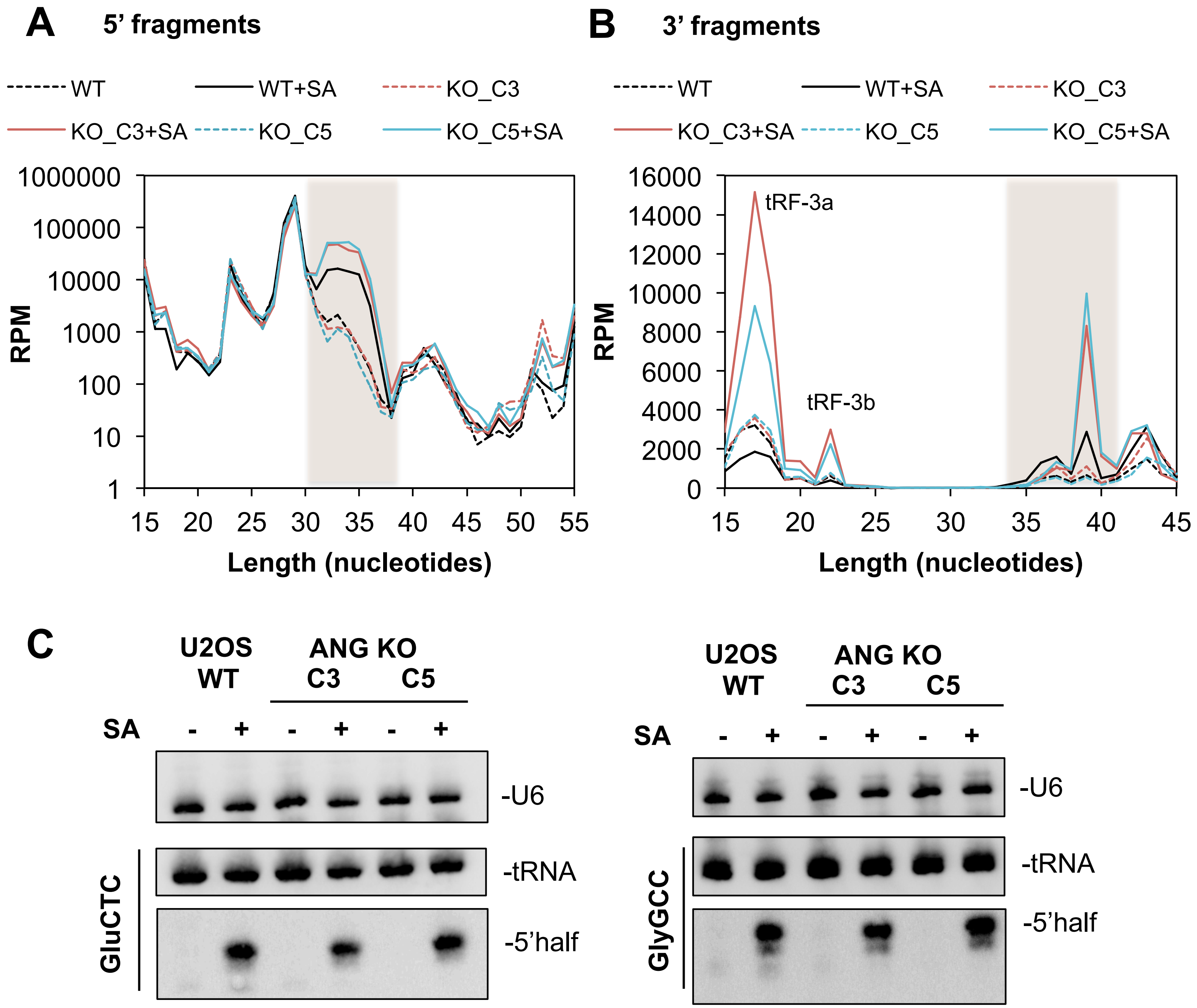


**Figure S5. Stress-induced tRNA halves in ANG knock-out cells.**
(A-B) Length distribution of all 5’ tRNA fragments (start from base 1) or 3’ tRNA fragments in U2OS wild-type and ANG KO cells without (dashed line) and with (solid line) sodium arsenite treatment. Y-axis: reads per million mapped tRF reads (on log scale for 5’ fragments) from small RNA-seq; X-axis: length of fragment. Shaded region corresponding to tRNA halves. Values were averaged from two biological replicates.
(C) Northern blot confirms arsenite-induced tRNA 5’ halves from $tRNA^{GluCTC}$ and $tRNA^{GlyGCC}$ in both wild-type and ANG knockout U2OS cells (Northern probe sequence see Methods).

Table S1. Statistics of small RNA-sequencing samples

| Sample | condition | total reads | reads with adaptors | after cutadapt (>=15nt) | average insert length | mapping% (STAR aligner) |
|---|---|---|---|---|---|---|
| 1 | HEK293T empty vector1 | 9,879,092 | 9,771,025 | 8,193,333 | 27 | 99 |
| 2 | HEK293T empty vector2 | 15,568,241 | 15,417,120 | 12,778,162 | 27 | 99 |
| 3 | HEK293T empty vector3 | 10,857,861 | 10,808,397 | 8,816,623 | 24 | 99 |
| 4 | HEK293T ANG OX1 | 13,021,013 | 12,874,111 | 11,369,891 | 31 | 99 |
| 5 | HEK293T ANG OX2 | 12,094,391 | 11,938,703 | 9,962,738 | 30 | 99 |
| 6 | HEK293T ANG OX3 | 20,471,981 | 20,351,276 | 16,553,229 | 28 | 99 |
| 7 | U2OS wild-type control1 | 19,459,539 | 19,270,126 | 10,107,456 | 24 | 98 |
| 8 | U2OS wild-type control2 | 24,448,091 | 24,313,985 | 16,256,108 | 24 | 96 |
| 9 | U2OS wild-type SA1 | 15,200,446 | 15,104,287 | 9,977,622 | 24 | 98 |
| 10 | U2OS wild-type SA2 | 28,178,878 | 28,055,204 | 18,777,064 | 24 | 98 |
| 11 | U2OS ANG KO clone3 control1 | 7,311,867 | 7,281,460 | 4,187,108 | 26 | 98 |
| 12 | U2OS ANG KO clone3 control2 | 16,026,564 | 15,955,173 | 9,189,606 | 23 | 96 |
| 13 | U2OS ANG KO clone3 SA1 | 14,840,747 | 14,742,476 | 8,350,771 | 23 | 98 |
| 14 | U2OS ANG KO clone3 SA2 | 21,869,702 | 21,690,547 | 14,313,118 | 24 | 96 |
| 15 | U2OS ANG KO clone5 contorl1 | 14,510,130 | 14,404,804 | 10,176,785 | 24 | 98 |
| 16 | U2OS ANG KO clone5 control2 | 27,049,958 | 26,816,633 | 19,648,853 | 25 | 98 |
| 17 | U2OS ANG KO clone5 SA1 | 16,290,627 | 16,179,439 | 10,410,913 | 25 | 98 |
| 18 | U2OS ANG KO clone5 SA2 | 48,767,939 | 48,399,658 | 30,805,686 | 25 | 97 |

Table S2. DESeq2 differential analysis of tRFs and miRs by HEK293T ANG over-expression (3 biological replicates)

*Table sorted first by adjusted p value and then by mean expression (EV). To simplify the table, only MeanExpression(EV) >10 are shown. Red font: p adjusted for multiple hypothesis testing is below 0.05.*

| name | MeanExpression (EV) | log2FoldChange (ANG/EV) | pvalue | p.adj |
|---|---|---|---|---|
| Glu_CTC_3half | 7988.24 | 4.96 | 1.52E-40 | 2.68E-37 |
| Glu_TTC_3half | 5086.55 | 5.05 | 7.94E-30 | 6.99E-27 |
| SeC_TCA_5half | 492.46 | 5.12 | 6.68E-16 | 3.92E-13 |
| Ser_GCT_3half | 104.57 | 5.59 | 2.26E-12 | 9.93E-10 |
| Glu_TTC_5half | 52315.87 | 4.25 | 3.22E-12 | 1.13E-09 |
| Lys_TTT_5half | 34618.67 | 3.87 | 4.97E-11 | 1.46E-08 |
| SeC_TCA_3half | 363.20 | 7.17 | 6.37E-11 | 1.60E-08 |
| Leu_CAG_5half | 83.77 | 3.47 | 1.08E-09 | 2.39E-07 |
| Ser_TGA_3half | 37.48 | 4.35 | 2.71E-09 | 5.29E-07 |
| Glu_CTC_5half | 133299.60 | 2.72 | 6.73E-09 | 1.08E-06 |
| Ala_CGC_5half | 51.35 | 3.12 | 6.19E-09 | 1.08E-06 |
| Asp_GTC_5half | 3877.90 | 3.57 | 3.82E-08 | 5.60E-06 |
| Ser_CGA_5half | 45.33 | 6.69 | 3.57E-07 | 4.83E-05 |
| Val_CAC_5half | 35126.93 | 2.69 | 1.15E-06 | 0.000144343 |
| Gly_CCC_5half | 64742.34 | 2.36 | 1.95E-06 | 0.000190921 |
| Lys_TTT_3half | 310.44 | 3.50 | 1.93E-06 | 0.000190921 |
| Asn_GTT_3half | 270.45 | 3.17 | 1.92E-06 | 0.000190921 |
| Thr_TGT_3half | 38.25 | 6.22 | 1.83E-06 | 0.000190921 |
| iMet_CAT_5half | 651.23 | 3.26 | 2.63E-06 | 0.000243204 |
| Gln_CTG_3half | 332.71 | 3.49 | 6.03E-06 | 0.000530921 |
| Ser_GCT_5half | 49.26 | 4.27 | 8.66E-06 | 0.000726173 |
| Val_CAC_3half | 33.96 | 4.47 | 2.94E-05 | 0.002354654 |
| Val_AAC_3half | 21.42 | 5.32 | 3.10E-05 | 0.002375096 |
| Leu_TAG_5half | 99.15 | 3.25 | 5.93E-05 | 0.00434716 |
| iMet_CAT_3half | 13.41 | 4.61 | 8.75E-05 | 0.006156805 |
| Val_TAC_5half | 3862.55 | 2.72 | 9.93E-05 | 0.006723382 |
| Val_AAC_5half | 12653.81 | 2.04 | 0.000117068 | 0.00735855 |
| miR-8069-2-3p | 439.57 | 3.26 | 0.000116263 | 0.00735855 |
| Trp_CCA_3half | 23.72 | 4.44 | 0.000139039 | 0.008438218 |
| Met_CAT_3half | 194.24 | 3.45 | 0.000143883 | 0.00844114 |
| Ala_AGC_5half | 19.64 | 2.39 | 0.000233028 | 0.013229976 |
| Ala_TGC_5half | 58.40 | 2.01 | 0.000306622 | 0.01686422 |
| Ser_AGA_3half | 47.43 | 3.39 | 0.000316435 | 0.016876554 |
| Asp_GTC_3half | 12865.93 | 2.97 | 0.000388977 | 0.020135295 |
| Gly_GCC_3half | 828.65 | 4.71 | 0.000463088 | 0.023209718 |
| Arg_TCG_3half | 43.09 | 3.02 | 0.000474744 | 0.023209718 |
| His_GTG_5half | 14163.10 | 3.29 | 0.000557996 | 0.026542528 |
| Gln_TTG_5half | 291.59 | 1.86 | 0.000602415 | 0.027901333 |
| SeC_TCA_misc | 117.41 | 2.99 | 0.000657443 | 0.029669221 |
| Thr_AGT_3half | 64.83 | 3.31 | 0.000854753 | 0.03760914 |
| Gly_GCC_5half | 343287.02 | 2.64 | 0.00089866 | 0.038576617 |
| Arg_CCG_5half | 142.93 | 3.64 | 0.002088701 | 0.087526516 |
| Thr_TGT_5half | 25.36 | 2.37 | 0.002429149 | 0.097461206 |
| Thr_CGT_3half | 46.76 | 3.44 | 0.002709166 | 0.105958486 |
| Leu_CAG_3half | 11.68 | 2.14 | 0.00332039 | 0.127041013 |
| Val_TAC_3half | 26.64 | 1.82 | 0.003731578 | 0.136824539 |
| Ser_CGA_3half | 20.76 | 2.96 | 0.003720719 | 0.136824539 |
| Leu_CAA_3half | 22.34 | 1.87 | 0.005245651 | 0.184646907 |
| Glu_CTC_tRF5 | 7125.08 | 2.15 | 0.006027749 | 0.208016451 |
| Gly_CCC_3half | 3333.92 | 3.42 | 0.006301555 | 0.213283412 |
| Gln_CTG_5half | 4257.28 | 2.03 | 0.007186018 | 0.238630037 |
| Pro_TGG_5half | 801.90 | 1.39 | 0.007414276 | 0.24110675 |
| Arg_CCG_3half | 51.76 | 2.13 | 0.007534586 | 0.24110675 |
| Pro_CGG_5half | 535.55 | 1.35 | 0.008104223 | 0.254704155 |
| His_GTG_3half | 15.06 | 3.23 | 0.01128103 | 0.348326543 |

| | | | | |
|---|---|---|---|---|
| Cys_GCA_5half | 19.45 | 2.03 | 0.013697291 | 0.415641942 |
| Ser_AGA_5half | 19.53 | 1.51 | 0.01956781 | 0.573989091 |
| miR-136-5p | 45.23 | 1.18 | 0.027560225 | 0.795180276 |
| miR-148a-3p | 910579.01 | -0.90 | 0.281033132 | 0.999893527 |
| Ser_TGA_tRF1 | 473506.81 | 0.14 | 0.84388536 | 0.999893527 |
| miR-10a-5p | 395272.61 | -0.47 | 0.407750495 | 0.999893527 |
| miR-20a-5p | 203754.39 | 0.41 | 0.460210205 | 0.999893527 |
| miR-92a-3p | 155740.07 | -0.23 | 0.588991542 | 0.999893527 |
| miR-17-5p | 139377.40 | 0.28 | 0.613461137 | 0.999893527 |
| Tyr_GTA_tRF3 | 134574.89 | -0.24 | 0.610929061 | 0.999893527 |
| miR-30d-5p | 101751.74 | -0.18 | 0.706560934 | 0.999893527 |
| miR-10b-5p | 99313.55 | -0.43 | 0.56892663 | 0.999893527 |
| miR-378a-3p | 80128.13 | -0.31 | 0.517066465 | 0.999893527 |
| miR-25-3p | 74074.22 | -0.23 | 0.540819589 | 0.999893527 |
| miR-7-5p | 71905.56 | -0.85 | 0.263609463 | 0.999893527 |
| miR-26a-5p | 69381.39 | 0.00 | 0.993195059 | 0.999893527 |
| miR-218-5p | 64141.12 | -0.37 | 0.581151847 | 0.999893527 |
| miR-103a-3p | 59448.05 | 0.26 | 0.480215866 | 0.999893527 |
| miR-92a-1-3p | 58888.74 | -0.24 | 0.644921646 | 0.999893527 |
| miR-218-2-5p | 57199.64 | -0.35 | 0.539025652 | 0.999893527 |
| miR-21-5p | 55814.84 | -0.24 | 0.604896171 | 0.999893527 |
| miR-27b-3p | 54772.86 | -0.25 | 0.592359807 | 0.999893527 |
| miR-148b-3p | 54671.48 | -0.38 | 0.361497844 | 0.999893527 |
| miR-423-3p | 54537.21 | -0.36 | 0.520564341 | 0.999893527 |
| miR-151a-3p | 50892.26 | -0.91 | 0.245902453 | 0.999893527 |
| miR-93-5p | 50391.28 | 0.03 | 0.966743596 | 0.999893527 |
| miR-221-3p | 49019.94 | -0.04 | 0.91747011 | 0.999893527 |
| miR-99a-5p | 47126.66 | -0.19 | 0.698340567 | 0.999893527 |
| let-7g-5p | 41421.65 | -0.16 | 0.749094435 | 0.999893527 |
| miR-222-3p | 38906.32 | 0.02 | 0.976973684 | 0.999893527 |
| miR-191-5p | 37899.69 | 0.09 | 0.86458195 | 0.999893527 |
| Gly_GCC_misc | 37814.13 | -0.30 | 0.395917809 | 0.999893527 |
| miR-30a-5p | 37539.45 | -0.31 | 0.568895124 | 0.999893527 |
| miR-7-3-5p | 33112.86 | -0.63 | 0.367211148 | 0.999893527 |
| let-7f-5p | 31973.38 | 0.12 | 0.730296558 | 0.999893527 |
| miR-128-3p | 31646.57 | -0.03 | 0.944295295 | 0.999893527 |
| miR-182-5p | 31555.32 | 0.05 | 0.939594343 | 0.999893527 |
| miR-26a-2-5p | 26596.29 | 0.02 | 0.973859212 | 0.999893527 |
| Lys_CTT_misc | 24204.17 | 0.07 | 0.887400845 | 0.999893527 |
| miR-30e-5p | 24077.18 | -0.02 | 0.968252043 | 0.999893527 |
| miR-183-5p | 23128.78 | -0.23 | 0.693094159 | 0.999893527 |
| miR-340-5p | 22843.76 | 0.16 | 0.663805135 | 0.999893527 |
| miR-103a-1-3p | 21980.72 | 0.08 | 0.846376558 | 0.999893527 |
| miR-186-5p | 19647.98 | 0.00 | 0.999391181 | 0.999893527 |
| miR-26b-5p | 18755.85 | 0.12 | 0.79375597 | 0.999893527 |
| miR-192-5p | 18080.44 | -0.66 | 0.247987591 | 0.999893527 |
| Glu_CTC_misc | 18055.27 | -0.20 | 0.668956953 | 0.999893527 |
| Arg_CCT_tRF1 | 17803.65 | 0.10 | 0.864905904 | 0.999893527 |
| miR-196b-5p | 17776.86 | -0.09 | 0.850779424 | 0.999893527 |
| let-7a-5p | 16915.57 | 0.06 | 0.895502744 | 0.999893527 |
| miR-101-2-3p | 16864.49 | -0.17 | 0.733906194 | 0.999893527 |
| miR-1260a-5p | 16457.87 | 0.04 | 0.936046001 | 0.999893527 |
| miR-99b-5p | 16356.09 | -0.33 | 0.398946064 | 0.999893527 |
| Cys_GCA_tRF3 | 16238.76 | -0.04 | 0.955383038 | 0.999893527 |
| miR-30c-1-5p | 15857.82 | 0.12 | 0.79753977 | 0.999893527 |
| Pro_TGG_tRF1 | 15838.69 | -0.21 | 0.621844898 | 0.999893527 |
| miR-1307-3p | 14187.01 | -0.40 | 0.592403766 | 0.999893527 |
| Ile_AAT_tRF1 | 14111.32 | 0.15 | 0.725111334 | 0.999893527 |
| let-7f-1-5p | 13354.52 | 0.06 | 0.858095313 | 0.999893527 |
| Gln_CTG_tRF3 | 12995.07 | -0.45 | 0.36643688 | 0.999893527 |
| miR-106b-3p | 12909.19 | -0.09 | 0.877787917 | 0.999893527 |
| Leu_CAG_tRF3 | 12718.90 | 0.02 | 0.970319029 | 0.999893527 |

| | | | | |
|---|---|---|---|---|
| Leu_CAA_tRF3 | 12691.88 | -0.17 | 0.765172661 | 0.999893527 |
| miR-532-5p | 12464.01 | -0.38 | 0.369637338 | 0.999893527 |
| miR-126-3p | 12414.30 | -0.08 | 0.907685545 | 0.999893527 |
| Gly_GCC_tRF3 | 11899.50 | 0.11 | 0.818923429 | 0.999893527 |
| His_GTG_misc | 11683.00 | -0.45 | 0.210131193 | 0.999893527 |
| miR-374b-5p | 11546.38 | 0.09 | 0.826091141 | 0.999893527 |
| let-7i-5p | 11349.37 | -0.09 | 0.873241796 | 0.999893527 |
| Met_CAT_misc | 10929.67 | -0.05 | 0.889508185 | 0.999893527 |
| miR-106b-5p | 10873.92 | 0.31 | 0.451460893 | 0.999893527 |
| miR-19a-3p | 10709.69 | 0.01 | 0.97230061 | 0.999893527 |
| miR-4521-5p | 10551.57 | 0.08 | 0.870778771 | 0.999893527 |
| miR-9-1-5p | 10409.15 | -0.24 | 0.66852306 | 0.999893527 |
| miR-425-5p | 10327.50 | -0.11 | 0.798534566 | 0.999893527 |
| miR-128-1-3p | 10231.10 | -0.04 | 0.920228744 | 0.999893527 |
| Gly_GCC_tRF5 | 10131.93 | 1.63 | 0.040602276 | 0.999893527 |
| Thr_TGT_tRF1 | 9801.20 | -0.21 | 0.593712599 | 0.999893527 |
| miR-30c-5p | 9488.11 | 0.05 | 0.902136432 | 0.999893527 |
| Asn_GTT_misc | 9328.78 | -0.59 | 0.331949906 | 0.999893527 |
| Cys_GCA_tRF1 | 9008.26 | 0.07 | 0.874410531 | 0.999893527 |
| miR-18a-5p | 8740.64 | 0.43 | 0.309966143 | 0.999893527 |
| Gly_CCC_misc | 8654.62 | -0.18 | 0.623999517 | 0.999893527 |
| let-7a-2-5p | 8642.57 | 0.09 | 0.855805983 | 0.999893527 |
| Asp_GTC_misc | 8598.80 | 0.18 | 0.650302539 | 0.999893527 |
| miR-125a-5p | 8548.06 | 0.05 | 0.926771398 | 0.999893527 |
| Lys_TTT_misc | 8439.02 | -0.02 | 0.949500408 | 0.999893527 |
| Val_CAC_tRF3 | 8406.54 | -0.05 | 0.924132634 | 0.999893527 |
| Lys_CTT_tRF1 | 8375.76 | -0.19 | 0.83027193 | 0.999893527 |
| miR-374a-5p | 8355.66 | 0.05 | 0.920116634 | 0.999893527 |
| miR-16-1-5p | 7936.42 | 0.30 | 0.527124826 | 0.999893527 |
| miR-615-3p | 7673.97 | -0.16 | 0.690746291 | 0.999893527 |
| miR-140-3p | 7631.98 | -0.02 | 0.953213659 | 0.999893527 |
| Ala_CGC_tRF1 | 7631.02 | 0.01 | 0.98013111 | 0.999893527 |
| Gln_CTG_misc | 7057.77 | -0.35 | 0.437435697 | 0.999893527 |
| miR-1180-3p | 7044.96 | -0.04 | 0.943590789 | 0.999893527 |
| Val_AAC_tRF3 | 6927.41 | -0.03 | 0.959946757 | 0.999893527 |
| miR-9-5p | 6687.29 | -0.22 | 0.670449174 | 0.999893527 |
| miR-744-5p | 6650.62 | -0.14 | 0.849342433 | 0.999893527 |
| Arg_CCT_misc | 6582.03 | 0.30 | 0.465224897 | 0.999893527 |
| miR-320a-3p | 6481.86 | 0.19 | 0.713744406 | 0.999893527 |
| miR-10a-3p | 6406.72 | -0.09 | 0.835562602 | 0.999893527 |
| Leu_TAG_tRF1 | 6313.90 | -0.38 | 0.438342105 | 0.999893527 |
| Phe_GAA_tRF3 | 6243.41 | 0.11 | 0.867404529 | 0.999893527 |
| miR-30e-3p | 6155.48 | -0.15 | 0.712887258 | 0.999893527 |
| miR-455-5p | 5968.95 | -0.43 | 0.287530448 | 0.999893527 |
| miR-378d-1-3p | 5801.13 | 1.13 | 0.105560299 | 0.999893527 |
| Gly_TCC_tRF1 | 5662.95 | 0.44 | 0.639968451 | 0.999893527 |
| miR-185-5p | 5641.16 | 0.15 | 0.685899563 | 0.999893527 |
| miR-210-3p | 5615.89 | -0.08 | 0.870320255 | 0.999893527 |
| Thr_AGT_tRF1 | 5596.77 | 0.12 | 0.776762961 | 0.999893527 |
| miR-130b-5p | 5579.46 | -0.11 | 0.785838202 | 0.999893527 |
| miR-107-3p | 5578.89 | -0.14 | 0.83889176 | 0.999893527 |
| miR-423-5p | 5554.53 | -0.04 | 0.94620268 | 0.999893527 |
| miR-32-5p | 5515.09 | -0.06 | 0.871320597 | 0.999893527 |
| miR-96-5p | 5413.42 | 0.00 | 0.994153024 | 0.999893527 |
| Ser_AGA_tRF3 | 5339.58 | 0.15 | 0.73106668 | 0.999893527 |
| miR-92b-3p | 5030.69 | -0.11 | 0.821052704 | 0.999893527 |
| miR-148a-5p | 5004.06 | -0.69 | 0.193165686 | 0.999893527 |
| miR-19b-2-3p | 4995.28 | 0.31 | 0.462889811 | 0.999893527 |
| miR-30b-5p | 4969.21 | 0.12 | 0.743916881 | 0.999893527 |
| Gln_TTG_tRF3 | 4787.38 | -0.18 | 0.633317115 | 0.999893527 |
| miR-339-5p | 4768.99 | 0.13 | 0.843912889 | 0.999893527 |
| Val_CAC_tRF5 | 4764.10 | 0.65 | 0.249119684 | 0.999893527 |

| | | | | |
|---|---|---|---|---|
| miR-106a-5p | 4704.56 | 0.35 | 0.557490915 | 0.999893527 |
| miR-221-5p | 4701.05 | -0.51 | 0.550270821 | 0.999893527 |
| Glu_TTC_misc | 4501.05 | -0.11 | 0.887648092 | 0.999893527 |
| miR-19b-3p | 4492.60 | 0.56 | 0.166206504 | 0.999893527 |
| Val_AAC_tRF5 | 4341.24 | 0.29 | 0.525685563 | 0.999893527 |
| miR-361-3p | 4295.12 | -0.65 | 0.252115935 | 0.999893527 |
| miR-181a-2-5p | 4281.96 | -0.03 | 0.944841226 | 0.999893527 |
| Thr_TGT_tRF3 | 4216.38 | 0.06 | 0.86052471 | 0.999893527 |
| miR-29a-3p | 4122.11 | -0.01 | 0.980800974 | 0.999893527 |
| Ser_TGA_tRF3 | 4092.84 | -0.16 | 0.713453751 | 0.999893527 |
| miR-16-5p | 3999.65 | 0.58 | 0.162574741 | 0.999893527 |
| Cys_GCA_misc | 3991.48 | -0.15 | 0.747966593 | 0.999893527 |
| miR-629-5p | 3961.51 | -0.19 | 0.714834907 | 0.999893527 |
| miR-30a-3p | 3806.22 | -0.39 | 0.56672238 | 0.999893527 |
| miR-320b-1-3p | 3787.50 | 0.28 | 0.630575967 | 0.999893527 |
| miR-24-1-3p | 3765.85 | 0.02 | 0.971834139 | 0.999893527 |
| miR-152-3p | 3719.34 | -0.11 | 0.759042595 | 0.999893527 |
| miR-941 | 3703.16 | -0.47 | 0.394248212 | 0.999893527 |
| miR-196a-5p | 3594.65 | 0.24 | 0.569832873 | 0.999893527 |
| miR-769-5p | 3593.31 | -0.41 | 0.37178659 | 0.999893527 |
| miR-22-3p | 3547.35 | -0.02 | 0.967843061 | 0.999893527 |
| miR-4483-3p | 3496.50 | -0.25 | 0.813617609 | 0.999893527 |
| miR-23b-3p | 3459.68 | 0.03 | 0.945553546 | 0.999893527 |
| Thr_CGT_tRF1 | 3422.63 | 0.42 | 0.543398174 | 0.999893527 |
| Pro_TGG_misc | 3372.90 | -0.17 | 0.656647217 | 0.999893527 |
| Pro_AGG_misc | 3344.56 | -0.21 | 0.660796641 | 0.999893527 |
| Glu_CTC_tRF1 | 3273.50 | 0.35 | 0.620557547 | 0.999893527 |
| Gly_TCC_5half | 3213.24 | 1.38 | 0.05693124 | 0.999893527 |
| iMet_CAT_misc | 3213.02 | -0.17 | 0.691722765 | 0.999893527 |
| Arg_TCG_tRF1 | 3207.73 | 0.04 | 0.935607372 | 0.999893527 |
| miR-15b-5p | 3151.78 | 0.17 | 0.64323432 | 0.999893527 |
| miR-16-2-3p | 3105.36 | 0.15 | 0.813041471 | 0.999893527 |
| miR-27a-3p | 3058.47 | 0.11 | 0.868831908 | 0.999893527 |
| miR-101-3p | 3022.23 | -0.27 | 0.699500628 | 0.999893527 |
| miR-196a-1-5p | 3018.75 | 0.10 | 0.796811663 | 0.999893527 |
| miR-941-5-3p | 2979.06 | -0.61 | 0.286071277 | 0.999893527 |
| miR-19b-1-3p | 2955.84 | -0.54 | 0.420627973 | 0.999893527 |
| Val_CAC_tRF1 | 2931.40 | 0.19 | 0.853143099 | 0.999893527 |
| miR-24-3p | 2864.37 | 0.10 | 0.821748646 | 0.999893527 |
| miR-563-3p | 2802.42 | 0.13 | 0.871182624 | 0.999893527 |
| miR-34a-5p | 2796.91 | -0.12 | 0.748918577 | 0.999893527 |
| miR-181d-5p | 2690.29 | -0.09 | 0.848433983 | 0.999893527 |
| miR-146b-5p | 2684.76 | -0.14 | 0.768143982 | 0.999893527 |
| Gly_GCC_tRF1 | 2682.78 | -0.23 | 0.685480152 | 0.999893527 |
| let-7c-5p | 2656.93 | 0.08 | 0.879134476 | 0.999893527 |
| Ala_AGC_misc | 2645.77 | -0.47 | 0.264128083 | 0.999893527 |
| miR-660-5p | 2631.13 | 0.06 | 0.913691027 | 0.999893527 |
| miR-181b-1-5p | 2608.68 | -0.01 | 0.976517555 | 0.999893527 |
| miR-625-3p | 2608.46 | -0.27 | 0.601303102 | 0.999893527 |
| Leu_AAG_tRF1 | 2596.21 | 0.02 | 0.979699852 | 0.999893527 |
| Lys_TTT_tRF3 | 2567.56 | 0.05 | 0.930460999 | 0.999893527 |
| Arg_ACG_tRF1 | 2542.44 | 0.15 | 0.746796173 | 0.999893527 |
| miR-7706-3p | 2533.44 | -0.30 | 0.556906737 | 0.999893527 |
| Tyr_GTA_tRF1 | 2470.13 | -0.07 | 0.93877794 | 0.999893527 |
| miR-374a-3p | 2419.53 | -0.61 | 0.247508387 | 0.999893527 |
| Val_TAC_tRF1 | 2407.44 | -0.21 | 0.729910397 | 0.999893527 |
| miR-652-3p | 2397.39 | 0.02 | 0.975589416 | 0.999893527 |
| miR-4653-3p | 2355.40 | -0.15 | 0.760219787 | 0.999893527 |
| miR-589-5p | 2340.40 | -0.43 | 0.474701311 | 0.999893527 |
| miR-107 | 2299.31 | -0.30 | 0.670898992 | 0.999893527 |
| miR-181a-5p | 2264.43 | -0.27 | 0.535066856 | 0.999893527 |
| Gly_CCC_tRF3 | 2264.16 | 0.07 | 0.904438899 | 0.999893527 |

| | | | | |
|---|---|---|---|---|
| miR-4454-5p | 2245.96 | 0.13 | 0.839808887 | 0.999893527 |
| Asp_GTC_tRF1 | 2240.70 | 0.24 | 0.73233562 | 0.999893527 |
| Tyr_GTA_tRF5 | 2217.53 | -0.28 | 0.700745067 | 0.999893527 |
| Glu_TTC_tRF3 | 2198.24 | 0.14 | 0.828069038 | 0.999893527 |
| miR-12135-3p | 2182.52 | 0.14 | 0.828420281 | 0.999893527 |
| miR-582-3p | 2181.89 | -0.78 | 0.168019472 | 0.999893527 |
| miR-92a-2-3p | 2133.43 | -0.09 | 0.870803198 | 0.999893527 |
| miR-708-5p | 2112.90 | 0.11 | 0.84928804 | 0.999893527 |
| miR-330-3p | 2112.10 | -0.11 | 0.828073961 | 0.999893527 |
| miR-450b-5p | 2101.83 | -0.24 | 0.557309311 | 0.999893527 |
| Gln_TTG_misc | 2071.34 | -0.29 | 0.436604394 | 0.999893527 |
| miR-361-5p | 2062.72 | 0.22 | 0.690362239 | 0.999893527 |
| Gly_TCC_misc | 2048.80 | 0.36 | 0.44276912 | 0.999893527 |
| miR-340-3p | 2047.96 | -0.04 | 0.947346133 | 0.999893527 |
| miR-28-3p | 2001.48 | -0.19 | 0.763051979 | 0.999893527 |
| miR-7-1-5p | 1971.52 | -0.30 | 0.561015989 | 0.999893527 |
| miR-454-3p | 1966.21 | 0.25 | 0.584580849 | 0.999893527 |
| Ser_GCT_tRF3 | 1938.85 | -0.01 | 0.984219277 | 0.999893527 |
| miR-1307-5p | 1918.78 | -0.39 | 0.326906278 | 0.999893527 |
| miR-345-5p | 1904.52 | -0.17 | 0.741736207 | 0.999893527 |
| miR-484 | 1882.30 | 0.12 | 0.762166199 | 0.999893527 |
| Gln_TTG_tRF1 | 1881.93 | 0.08 | 0.846388484 | 0.999893527 |
| miR-140-5p | 1848.51 | 0.06 | 0.929294293 | 0.999893527 |
| miR-622-3p | 1833.10 | -0.06 | 0.914540539 | 0.999893527 |
| miR-31-5p | 1789.90 | 0.35 | 0.402674414 | 0.999893527 |
| miR-7975-3p | 1780.12 | 0.17 | 0.815994904 | 0.999893527 |
| miR-1261-5p | 1748.55 | 0.26 | 0.561264418 | 0.999893527 |
| miR-1271-5p | 1705.83 | 0.00 | 0.993370177 | 0.999893527 |
| miR-711-5p | 1693.49 | 0.23 | 0.755091393 | 0.999893527 |
| miR-450a-1-5p | 1681.81 | -0.20 | 0.665251575 | 0.999893527 |
| miR-99b-3p | 1661.81 | -0.47 | 0.415014533 | 0.999893527 |
| miR-503-5p | 1645.40 | 0.14 | 0.719968465 | 0.999893527 |
| Tyr_GTA_misc | 1627.67 | -0.41 | 0.676676267 | 0.999893527 |
| miR-362-5p | 1618.26 | -0.01 | 0.977263835 | 0.999893527 |
| miR-590-3p | 1595.82 | 0.22 | 0.694953363 | 0.999893527 |
| His_GTG_tRF1 | 1591.83 | -0.02 | 0.968825702 | 0.999893527 |
| miR-15b-3p | 1575.55 | 0.08 | 0.907384947 | 0.999893527 |
| miR-5100-3p | 1573.98 | 0.10 | 0.865235473 | 0.999893527 |
| miR-143-3p | 1570.33 | 0.31 | 0.648284145 | 0.999893527 |
| Arg_ACG_tRF3 | 1548.49 | -0.04 | 0.93563746 | 0.999893527 |
| Glu_TTC_tRF5 | 1525.58 | 1.05 | 0.055291197 | 0.999893527 |
| miR-542-3p | 1514.00 | 0.23 | 0.77541069 | 0.999893527 |
| miR-500a-3p | 1485.66 | -0.25 | 0.462325759 | 0.999893527 |
| Val_CAC_misc | 1475.56 | -0.12 | 0.768795306 | 0.999893527 |
| miR-576-3p | 1456.51 | -0.77 | 0.218989519 | 0.999893527 |
| miR-98-5p | 1453.19 | -0.11 | 0.838485015 | 0.999893527 |
| miR-125b-2-5p | 1451.79 | 0.04 | 0.952028071 | 0.999893527 |
| miR-484-5p | 1432.71 | 0.01 | 0.974434503 | 0.999893527 |
| Gly_TCC_tRF3 | 1431.86 | 0.11 | 0.836118166 | 0.999893527 |
| let-7e-5p | 1421.67 | 0.12 | 0.782850272 | 0.999893527 |
| miR-548o-3p | 1421.45 | -0.78 | 0.306781185 | 0.999893527 |
| miR-369-3p | 1419.09 | -0.17 | 0.866748846 | 0.999893527 |
| Pro_CGG_misc | 1413.33 | -0.19 | 0.697639618 | 0.999893527 |
| miR-196a-2-5p | 1410.32 | 0.20 | 0.620729393 | 0.999893527 |
| Lys_CTT_tRF5 | 1402.46 | 0.78 | 0.260397644 | 0.999893527 |
| Val_AAC_misc | 1390.38 | -0.20 | 0.631413224 | 0.999893527 |
| miR-218-1-5p | 1382.47 | -0.19 | 0.818671465 | 0.999893527 |
| miR-18a-3p | 1381.85 | 0.14 | 0.821742054 | 0.999893527 |
| miR-342-3p | 1371.61 | -0.12 | 0.737236088 | 0.999893527 |
| miR-127-3p | 1341.78 | 0.83 | 0.591204213 | 0.999893527 |
| miR-455-3p | 1326.64 | -0.06 | 0.941436185 | 0.999893527 |
| miR-194-5p | 1314.61 | 0.18 | 0.725423672 | 0.999893527 |

| | | | | |
|---|---|---|---|---|
| miR-25-5p | 1298.15 | 0.06 | 0.942156716 | 0.999893527 |
| miR-197-3p | 1291.93 | -0.09 | 0.793237318 | 0.999893527 |
| miR-1260b-5p | 1291.61 | 0.25 | 0.500543827 | 0.999893527 |
| miR-130b-3p | 1286.53 | 0.46 | 0.453413006 | 0.999893527 |
| miR-95-3p | 1283.85 | -0.17 | 0.701087738 | 0.999893527 |
| miR-3919-3p | 1271.59 | 0.00 | 0.999590316 | 0.999893527 |
| miR-486-5p | 1265.07 | -0.18 | 0.696094666 | 0.999893527 |
| miR-339-3p | 1263.55 | -0.27 | 0.604622002 | 0.999893527 |
| miR-505-3p | 1263.44 | -0.07 | 0.905307624 | 0.999893527 |
| Ser_CGA_tRF3 | 1261.96 | -0.03 | 0.955712539 | 0.999893527 |
| Arg_CCG_tRF1 | 1258.08 | 0.32 | 0.741413359 | 0.999893527 |
| Thr_AGT_tRF3 | 1251.27 | -0.02 | 0.964898767 | 0.999893527 |
| miR-328-3p | 1231.78 | -0.18 | 0.650101455 | 0.999893527 |
| Ile_AAT_misc | 1229.00 | 0.13 | 0.77439112 | 0.999893527 |
| miR-181b-5p | 1198.62 | -0.06 | 0.877990121 | 0.999893527 |
| let-7f-2-5p | 1186.96 | 0.01 | 0.985155532 | 0.999893527 |
| miR-3613-5p | 1178.46 | 0.06 | 0.905924082 | 0.999893527 |
| miR-3615-3p | 1173.03 | -0.38 | 0.355452604 | 0.999893527 |
| Gly_CCC_tRF5 | 1162.75 | 1.37 | 0.055839095 | 0.999893527 |
| miR-421-3p | 1133.18 | 0.31 | 0.638489016 | 0.999893527 |
| miR-20b-5p | 1132.07 | 0.33 | 0.580455233 | 0.999893527 |
| miR-424-3p | 1109.53 | 0.17 | 0.700637609 | 0.999893527 |
| miR-378h-5p | 1104.51 | -0.34 | 0.341121047 | 0.999893527 |
| miR-4633-5p | 1101.40 | -0.22 | 0.670775165 | 0.999893527 |
| miR-9-1-3p | 1089.59 | 0.15 | 0.836036759 | 0.999893527 |
| miR-215-5p | 1055.09 | -0.09 | 0.870106898 | 0.999893527 |
| miR-23a-3p | 1018.86 | 0.24 | 0.615822993 | 0.999893527 |
| miR-132-3p | 1006.08 | -0.12 | 0.763562721 | 0.999893527 |
| miR-92a-1-5p | 1003.02 | -0.02 | 0.974119625 | 0.999893527 |
| miR-3158-2-3p | 1001.85 | -0.88 | 0.282363842 | 0.999893527 |
| Leu_CAA_misc | 999.59 | 0.05 | 0.907938037 | 0.999893527 |
| miR-641-5p | 994.03 | -0.15 | 0.752115406 | 0.999893527 |
| miR-375-3p | 980.68 | -0.13 | 0.743725739 | 0.999893527 |
| miR-9-3p | 975.18 | 0.25 | 0.723638126 | 0.999893527 |
| miR-7974-3p | 973.10 | -0.41 | 0.529079335 | 0.999893527 |
| miR-125b-5p | 966.84 | -0.01 | 0.980490016 | 0.999893527 |
| miR-424-5p | 960.52 | 0.48 | 0.354761684 | 0.999893527 |
| miR-331-3p | 945.16 | -0.20 | 0.703794126 | 0.999893527 |
| Trp_CCA_tRF3 | 940.92 | -0.16 | 0.696116454 | 0.999893527 |
| Arg_TCT_misc | 933.45 | -0.07 | 0.922837271 | 0.999893527 |
| miR-671-5p | 916.94 | 0.42 | 0.509410614 | 0.999893527 |
| miR-1301-3p | 897.04 | 0.05 | 0.929337132 | 0.999893527 |
| Ser_GCT_tRF1 | 888.45 | 0.09 | 0.854032826 | 0.999893527 |
| let-7d-5p | 884.47 | 0.22 | 0.800108336 | 0.999893527 |
| miR-598-3p | 884.14 | -0.07 | 0.894062763 | 0.999893527 |
| Pro_AGG_5half | 868.65 | 0.86 | 0.089734986 | 0.999893527 |
| miR-1296-5p | 863.02 | -0.28 | 0.650934835 | 0.999893527 |
| miR-1304-3p | 859.21 | -0.43 | 0.453883024 | 0.999893527 |
| Ala_TGC_tRF1 | 859.20 | 0.23 | 0.553079145 | 0.999893527 |
| Asn_GTT_tRF1 | 842.01 | 0.17 | 0.871027416 | 0.999893527 |
| miR-132-5p | 840.77 | -0.03 | 0.94969891 | 0.999893527 |
| miR-6820-3p | 833.73 | -0.10 | 0.789801276 | 0.999893527 |
| Val_AAC_tRF1 | 832.04 | -0.04 | 0.940300508 | 0.999893527 |
| Arg_CCG_misc | 831.46 | -0.03 | 0.950628625 | 0.999893527 |
| miR-149-5p | 830.79 | -0.25 | 0.496855497 | 0.999893527 |
| miR-421 | 828.68 | 0.36 | 0.564814862 | 0.999893527 |
| miR-1269b-5p | 818.12 | -0.50 | 0.400894072 | 0.999893527 |
| miR-194-2-5p | 817.61 | 0.26 | 0.590370464 | 0.999893527 |
| His_GTG_leader | 813.76 | 0.14 | 0.725897432 | 0.999893527 |
| Val_TAC_tRF5 | 810.52 | -0.03 | 0.95737299 | 0.999893527 |
| miR-760-5p | 806.39 | -0.35 | 0.526861825 | 0.999893527 |
| Thr_TGT_misc | 805.99 | 0.27 | 0.510102839 | 0.999893527 |

| | | | | |
|---|---|---|---|---|
| miR-378a-5p | 801.37 | 0.03 | 0.962907053 | 0.999893527 |
| miR-181a-2-3p | 793.43 | -0.18 | 0.644599532 | 0.999893527 |
| Lys_CTT_tRF3 | 792.86 | -0.08 | 0.824549758 | 0.999893527 |
| Val_TAC_misc | 782.24 | 0.00 | 0.991976327 | 0.999893527 |
| Gln_CTG_tRF1 | 770.53 | 0.03 | 0.947883823 | 0.999893527 |
| miR-296-3p | 764.26 | -0.17 | 0.838057781 | 0.999893527 |
| let-7a-3p | 757.71 | 0.03 | 0.964463273 | 0.999893527 |
| miR-15a-5p | 757.30 | 0.60 | 0.259821728 | 0.999893527 |
| let-7b-5p | 752.22 | 0.24 | 0.757997843 | 0.999893527 |
| miR-4455-5p | 738.99 | 0.17 | 0.701030982 | 0.999893527 |
| Asn_GTT_tRF3 | 735.87 | 0.34 | 0.598839059 | 0.999893527 |
| miR-7977-5p | 728.39 | 0.14 | 0.731916205 | 0.999893527 |
| miR-671-3p | 715.46 | -0.48 | 0.441596807 | 0.999893527 |
| miR-128-2-3p | 709.73 | -0.16 | 0.739861569 | 0.999893527 |
| miR-17-3p | 708.94 | -0.12 | 0.750450139 | 0.999893527 |
| miR-28-5p | 698.78 | -0.02 | 0.954771012 | 0.999893527 |
| Ala_TGC_misc | 695.70 | 0.05 | 0.920957871 | 0.999893527 |
| Phe_GAA_misc | 687.91 | -0.07 | 0.891822467 | 0.999893527 |
| miR-425-3p | 687.84 | -0.21 | 0.622467194 | 0.999893527 |
| miR-187-3p | 679.40 | -0.17 | 0.775899555 | 0.999893527 |
| miR-618 | 676.78 | -0.01 | 0.983142354 | 0.999893527 |
| miR-502-3p | 667.79 | -0.07 | 0.876382709 | 0.999893527 |
| miR-194-1-5p | 660.03 | 0.04 | 0.943967878 | 0.999893527 |
| miR-1269b | 655.33 | -0.27 | 0.607578464 | 0.999893527 |
| miR-4473-3p | 651.59 | -0.26 | 0.597573817 | 0.999893527 |
| miR-942-5p | 650.43 | 0.09 | 0.828894865 | 0.999893527 |
| miR-548k | 636.66 | -0.17 | 0.678297548 | 0.999893527 |
| miR-301a-5p | 629.89 | -0.05 | 0.941899473 | 0.999893527 |
| miR-126-5p | 626.33 | -0.01 | 0.990168699 | 0.999893527 |
| Arg_TCG_misc | 621.00 | 0.11 | 0.803007307 | 0.999893527 |
| miR-181b-2-5p | 616.18 | 0.07 | 0.911380157 | 0.999893527 |
| miR-374b-3p | 609.15 | -0.25 | 0.599806279 | 0.999893527 |
| miR-548av-3p | 594.75 | -1.13 | 0.141468283 | 0.999893527 |
| miR-577-5p | 592.16 | 0.09 | 0.840403554 | 0.999893527 |
| miR-324-3p | 589.37 | -0.06 | 0.907735072 | 0.999893527 |
| Trp_CCA_tRF1 | 582.67 | 0.01 | 0.99167289 | 0.999893527 |
| miR-501-3p | 579.56 | -0.03 | 0.940881133 | 0.999893527 |
| Lys_TTT_tRF5 | 573.94 | 0.72 | 0.317406145 | 0.999893527 |
| miR-10b-3p | 572.24 | -0.02 | 0.952509796 | 0.999893527 |
| miR-379-5p | 564.40 | 0.59 | 0.684385974 | 0.999893527 |
| miR-582-5p | 563.59 | 0.20 | 0.791751872 | 0.999893527 |
| miR-93-3p | 562.00 | -0.02 | 0.969877028 | 0.999893527 |
| miR-181c-3p | 557.45 | -0.15 | 0.718537822 | 0.999893527 |
| miR-181a-1-3p | 556.35 | -0.39 | 0.415095759 | 0.999893527 |
| miR-7974 | 551.88 | -0.47 | 0.398095578 | 0.999893527 |
| miR-330-5p | 549.06 | -0.34 | 0.439538139 | 0.999893527 |
| Leu_CAG_misc | 531.92 | -0.14 | 0.823242353 | 0.999893527 |
| miR-1275-5p | 530.38 | 0.43 | 0.555890138 | 0.999893527 |
| miR-32-3p | 530.23 | 0.00 | 0.993956453 | 0.999893527 |
| miR-365b-3p | 529.86 | 0.34 | 0.664326814 | 0.999893527 |
| Thr_CGT_misc | 527.08 | -0.17 | 0.65614408 | 0.999893527 |
| miR-450a-5p | 522.03 | -0.01 | 0.987331264 | 0.999893527 |
| miR-365a-3p | 521.95 | 0.18 | 0.797121939 | 0.999893527 |
| iMet_CAT_tRF3 | 508.46 | 0.10 | 0.908855408 | 0.999893527 |
| miR-576-5p | 505.11 | -0.43 | 0.334417848 | 0.999893527 |
| Ser_AGA_tRF1 | 498.91 | 0.36 | 0.620743657 | 0.999893527 |
| miR-103a-2-3p | 497.05 | 0.31 | 0.448683486 | 0.999893527 |
| miR-181a-3p | 496.24 | -0.27 | 0.464792451 | 0.999893527 |
| miR-4791-5p | 495.37 | -0.05 | 0.951733248 | 0.999893527 |
| miR-148b-5p | 494.34 | -0.21 | 0.652429607 | 0.999893527 |
| Thr_CGT_tRF3 | 492.29 | 0.26 | 0.619395947 | 0.999893527 |
| miR-200c-3p | 485.26 | 0.41 | 0.612032224 | 0.999893527 |

| | | | | |
|---|---|---|---|---|
| Ser_AGA_tRF5 | 476.10 | 0.67 | 0.274515407 | 0.999893527 |
| miR-30d-3p | 461.68 | -0.41 | 0.454913761 | 0.999893527 |
| miR-4443-5p | 460.36 | 0.25 | 0.656640441 | 0.999893527 |
| miR-548e-3p | 458.64 | -0.16 | 0.759258238 | 0.999893527 |
| miR-874-3p | 450.59 | -0.04 | 0.957467552 | 0.999893527 |
| miR-543 | 448.24 | 0.82 | 0.730074816 | 0.999893527 |
| miR-125b-2-3p | 439.45 | -0.22 | 0.665834726 | 0.999893527 |
| miR-591-5p | 431.23 | 0.24 | 0.661193073 | 0.999893527 |
| miR-196a-2-3p | 430.78 | -0.61 | 0.377877514 | 0.999893527 |
| miR-2277-5p | 426.58 | -0.36 | 0.482397247 | 0.999893527 |
| Ile_AAT_tRF5 | 425.29 | -0.20 | 0.782657839 | 0.999893527 |
| Thr_AGT_misc | 422.81 | 0.31 | 0.607866501 | 0.999893527 |
| Phe_GAA_tRF1 | 418.25 | 0.54 | 0.305576331 | 0.999893527 |
| miR-193b-3p | 415.56 | -0.23 | 0.602675772 | 0.999893527 |
| miR-548g-3p | 414.93 | -0.28 | 0.727336993 | 0.999893527 |
| miR-708-3p | 414.51 | -0.43 | 0.435658138 | 0.999893527 |
| Arg_TCT_tRF3 | 413.87 | 0.02 | 0.976344029 | 0.999893527 |
| miR-301b-3p | 412.99 | -0.22 | 0.615317096 | 0.999893527 |
| miR-199a-2-3p | 404.53 | -0.07 | 0.906216657 | 0.999893527 |
| miR-1303-3p | 404.16 | -0.09 | 0.908713625 | 0.999893527 |
| miR-195-5p | 392.63 | 0.32 | 0.592151245 | 0.999893527 |
| Leu_TAA_misc | 390.02 | 0.17 | 0.800497326 | 0.999893527 |
| miR-3182-5p | 388.17 | 0.11 | 0.80679622 | 0.999893527 |
| let-7a-3-3p | 384.64 | 0.19 | 0.837353913 | 0.999893527 |
| miR-6748-3p | 381.87 | 0.27 | 0.609093287 | 0.999893527 |
| Ala_CGC_misc | 381.75 | -0.25 | 0.620475524 | 0.999893527 |
| miR-4286-5p | 379.88 | -0.03 | 0.937656362 | 0.999893527 |
| miR-193a-5p | 376.04 | 0.06 | 0.93951836 | 0.999893527 |
| miR-1226-3p | 374.63 | -0.07 | 0.925818984 | 0.999893527 |
| miR-548bc-3p | 373.50 | -0.72 | 0.248567952 | 0.999893527 |
| Ser_GCT_misc | 360.83 | 0.12 | 0.761534621 | 0.999893527 |
| miR-382-5p | 357.73 | 0.32 | 0.631860458 | 0.999893527 |
| Arg_ACG_misc | 354.03 | -0.08 | 0.838908876 | 0.999893527 |
| miR-12130-3p | 347.81 | 0.08 | 0.925312287 | 0.999893527 |
| Lys_TTT_tRF1 | 347.39 | 0.18 | 0.785417718 | 0.999893527 |
| miR-331-5p | 347.11 | 0.04 | 0.906776137 | 0.999893527 |
| miR-3198-1-5p | 345.09 | 0.20 | 0.690498786 | 0.999893527 |
| miR-449c-5p | 339.82 | -0.28 | 0.470370984 | 0.999893527 |
| miR-7-1-3p | 339.47 | 0.11 | 0.797669887 | 0.999893527 |
| miR-199b-5p | 330.52 | -0.14 | 0.813044939 | 0.999893527 |
| miR-29b-1-3p | 330.22 | -0.36 | 0.618004499 | 0.999893527 |
| miR-3909 | 330.20 | -0.37 | 0.380982113 | 0.999893527 |
| miR-548aq-3p | 326.83 | -0.43 | 0.423453325 | 0.999893527 |
| Leu_CAG_tRF5 | 326.04 | 0.14 | 0.777285972 | 0.999893527 |
| Ser_CGA_tRF1 | 324.15 | 0.03 | 0.961708916 | 0.999893527 |
| miR-935-3p | 324.10 | -0.12 | 0.870257794 | 0.999893527 |
| miR-6073-3p | 322.62 | 0.24 | 0.62280707 | 0.999893527 |
| miR-1-3p | 321.63 | 0.28 | 0.47990134 | 0.999893527 |
| miR-30c-2-3p | 320.46 | -0.28 | 0.642031563 | 0.999893527 |
| miR-34c-5p | 318.86 | -0.34 | 0.402251594 | 0.999893527 |
| Ala_TGC_tRF5 | 310.70 | -0.01 | 0.985428635 | 0.999893527 |
| miR-676-3p | 308.85 | -0.18 | 0.762810327 | 0.999893527 |
| miR-139-5p | 308.83 | 0.24 | 0.75858876 | 0.999893527 |
| miR-548d-1-5p | 306.32 | -0.06 | 0.879484241 | 0.999893527 |
| miR-500b-5p | 305.06 | 0.07 | 0.87373224 | 0.999893527 |
| miR-497-5p | 304.16 | 0.16 | 0.843857758 | 0.999893527 |
| miR-3158-3p | 303.38 | -0.33 | 0.441321898 | 0.999893527 |
| miR-27b-5p | 298.81 | 0.01 | 0.984723151 | 0.999893527 |
| miR-301a-3p | 297.72 | -0.05 | 0.941342662 | 0.999893527 |
| Leu_AAG_tRF5 | 290.67 | 0.84 | 0.270412577 | 0.999893527 |
| miR-20a-3p | 288.43 | 0.31 | 0.501559533 | 0.999893527 |
| miR-6765-3p | 284.85 | 0.01 | 0.980411249 | 0.999893527 |

| | | | | |
|---|---|---|---|---|
| miR-501-5p | 281.08 | 0.16 | 0.772212762 | 0.999893527 |
| miR-543-3p | 281.04 | 0.86 | 0.724576295 | 0.999893527 |
| miR-4677-3p | 278.96 | -0.42 | 0.477963317 | 0.999893527 |
| Glu_CTC_tRF3 | 277.27 | -0.27 | 0.561114785 | 0.999893527 |
| Leu_AAG_misc | 276.73 | 0.18 | 0.732542619 | 0.999893527 |
| miR-296-5p | 269.94 | 0.28 | 0.767483713 | 0.999893527 |
| miR-1303 | 267.64 | -0.16 | 0.846144145 | 0.999893527 |
| miR-409-3p | 266.16 | 0.84 | 0.557256144 | 0.999893527 |
| miR-200b-3p | 265.24 | 0.58 | 0.503189664 | 0.999893527 |
| miR-370-3p | 265.05 | 0.84 | 0.57889845 | 0.999893527 |
| Trp_CCA_misc | 264.78 | 0.03 | 0.945806701 | 0.999893527 |
| Arg_TCG_tRF3 | 262.95 | 0.08 | 0.886824034 | 0.999893527 |
| miR-760-3p | 262.25 | -0.28 | 0.609783836 | 0.999893527 |
| miR-1278 | 262.16 | -0.51 | 0.287298177 | 0.999893527 |
| miR-1246 | 262.06 | 0.21 | 0.825431947 | 0.999893527 |
| miR-320c-1-3p | 261.18 | 0.25 | 0.544483189 | 0.999893527 |
| miR-212-5p | 258.89 | 0.03 | 0.960783579 | 0.999893527 |
| miR-5701-3-5p | 255.66 | -0.22 | 0.779651246 | 0.999893527 |
| miR-532-3p | 255.16 | -0.13 | 0.794923557 | 0.999893527 |
| Ala_AGC_tRF5 | 254.33 | -0.27 | 0.553575476 | 0.999893527 |
| miR-146a-5p | 253.85 | 0.22 | 0.721414287 | 0.999893527 |
| miR-23c-3p | 253.34 | 0.06 | 0.905856772 | 0.999893527 |
| miR-27a-5p | 250.97 | -0.50 | 0.439103328 | 0.999893527 |
| miR-548am-3p | 250.18 | -1.03 | 0.161219241 | 0.999893527 |
| miR-411-5p | 247.64 | 0.70 | 0.609705229 | 0.999893527 |
| miR-769-3p | 247.33 | -0.31 | 0.457560698 | 0.999893527 |
| miR-641 | 243.09 | -0.04 | 0.93499055 | 0.999893527 |
| miR-1306-5p | 241.55 | 0.06 | 0.895096779 | 0.999893527 |
| Ala_CGC_tRF5 | 241.12 | 0.04 | 0.948579044 | 0.999893527 |
| miR-454-5p | 240.47 | -0.15 | 0.798877911 | 0.999893527 |
| miR-3159-3p | 238.09 | 0.00 | 0.996703329 | 0.999893527 |
| miR-548i-4-5p | 236.19 | -0.17 | 0.6917795 | 0.999893527 |
| miR-3200-3p | 234.31 | -0.30 | 0.621393381 | 0.999893527 |
| miR-381-3p | 234.05 | 0.78 | 0.582545917 | 0.999893527 |
| Asp_GTC_tRF5 | 233.05 | 0.59 | 0.284214912 | 0.999893527 |
| miR-628-5p | 231.74 | 0.17 | 0.834213248 | 0.999893527 |
| miR-618-5p | 231.46 | -0.15 | 0.800773402 | 0.999893527 |
| miR-1306-3p | 230.51 | 0.18 | 0.622847286 | 0.999893527 |
| miR-7706 | 229.12 | -0.23 | 0.632337816 | 0.999893527 |
| miR-8085-3p | 227.93 | 0.24 | 0.620263635 | 0.999893527 |
| Asp_GTC_tRF3 | 227.85 | -0.24 | 0.652044654 | 0.999893527 |
| miR-1268a | 227.56 | -0.05 | 0.960042159 | 0.999893527 |
| miR-4705-5p | 226.80 | -0.06 | 0.948193383 | 0.999893527 |
| miR-6828-5p | 226.05 | 0.05 | 0.940398731 | 0.999893527 |
| miR-324-5p | 224.88 | 0.22 | 0.593758933 | 0.999893527 |
| Leu_TAG_misc | 223.88 | 0.09 | 0.853430914 | 0.999893527 |
| miR-129-5p | 223.35 | -0.33 | 0.530063059 | 0.999893527 |
| miR-548f-1-3p | 220.06 | -0.49 | 0.350194098 | 0.999893527 |
| miR-6529-3p | 218.79 | -0.04 | 0.945140887 | 0.999893527 |
| miR-4664-3p | 218.56 | -0.33 | 0.538979466 | 0.999893527 |
| miR-4746-5p | 215.51 | -0.46 | 0.322448889 | 0.999893527 |
| miR-33a-3p | 213.87 | -0.12 | 0.824030863 | 0.999893527 |
| miR-4787-3p | 211.29 | 0.71 | 0.44882501 | 0.999893527 |
| miR-224-5p | 208.64 | 0.15 | 0.899580864 | 0.999893527 |
| miR-4326-5p | 208.07 | -0.40 | 0.42408346 | 0.999893527 |
| miR-29c-3p | 205.37 | -0.06 | 0.885232397 | 0.999893527 |
| miR-4699-5p | 201.24 | 0.08 | 0.909208736 | 0.999893527 |
| miR-92b-5p | 197.58 | -0.44 | 0.294696023 | 0.999893527 |
| miR-3074-5p | 194.99 | -0.37 | 0.351148483 | 0.999893527 |
| miR-1843-5p | 194.84 | -0.18 | 0.71330064 | 0.999893527 |
| iMet_CAT_tRF1 | 194.66 | 0.56 | 0.25998267 | 0.999893527 |
| Leu_TAA_tRF1 | 190.98 | 0.06 | 0.947254267 | 0.999893527 |

| | | | | |
|---|---|---|---|---|
| miR-574-3p | 188.66 | 0.42 | 0.373032594 | 0.999893527 |
| miR-433-3p | 188.30 | 0.35 | 0.795422869 | 0.999893527 |
| miR-29c-5p | 188.01 | -0.35 | 0.53672809 | 0.999893527 |
| miR-6128-3p | 187.93 | -0.44 | 0.277661585 | 0.999893527 |
| miR-550a-1-5p | 186.15 | -0.38 | 0.427885616 | 0.999893527 |
| Gln_TTG_leader | 186.05 | 0.63 | 0.532781325 | 0.999893527 |
| miR-203a-3p | 185.89 | 0.22 | 0.741578754 | 0.999893527 |
| miR-100-5p | 185.80 | -0.06 | 0.912454795 | 0.999893527 |
| miR-338-5p | 185.08 | 0.12 | 0.772768958 | 0.999893527 |
| miR-6720-3p | 184.33 | -0.06 | 0.890642608 | 0.999893527 |
| miR-5001-3p | 184.05 | -0.06 | 0.935170822 | 0.999893527 |
| miR-19b-1-5p | 183.76 | 0.06 | 0.895270265 | 0.999893527 |
| miR-181c-5p | 182.29 | -0.25 | 0.682899659 | 0.999893527 |
| miR-1304-5p | 182.14 | -0.21 | 0.683113997 | 0.999893527 |
| miR-877-5p | 181.77 | -0.10 | 0.846931334 | 0.999893527 |
| miR-10395-3p | 180.41 | 0.50 | 0.312546643 | 0.999893527 |
| miR-200a-3p | 178.47 | 0.07 | 0.881563983 | 0.999893527 |
| miR-652-5p | 178.06 | 0.12 | 0.842487917 | 0.999893527 |
| let-7f-2-3p | 177.76 | -0.67 | 0.301178953 | 0.999893527 |
| miR-3168-5p | 173.69 | -0.13 | 0.923630471 | 0.999893527 |
| miR-7976-5p | 173.00 | -0.14 | 0.723250555 | 0.999893527 |
| miR-553-3p | 172.27 | 1.02 | 0.090473118 | 0.999893527 |
| miR-1268b-5p | 172.14 | -0.11 | 0.884431653 | 0.999893527 |
| miR-129-1-5p | 171.84 | -0.43 | 0.472133938 | 0.999893527 |
| miR-22-5p | 170.81 | 0.26 | 0.654088578 | 0.999893527 |
| miR-548bc | 170.59 | -0.60 | 0.343701924 | 0.999893527 |
| Ile_TAT_misc | 169.91 | -0.30 | 0.711779645 | 0.999893527 |
| miR-6075-3p | 169.73 | 0.71 | 0.517576268 | 0.999893527 |
| miR-2392-3p | 168.95 | 0.13 | 0.866833533 | 0.999893527 |
| Ser_TGA_tRF5 | 168.68 | 0.68 | 0.278038962 | 0.999893527 |
| SeC_TCA_tRF1 | 166.11 | 0.48 | 0.514000063 | 0.999893527 |
| miR-2110-5p | 162.84 | -0.11 | 0.815189238 | 0.999893527 |
| miR-378d | 162.80 | -0.01 | 0.98599962 | 0.999893527 |
| miR-3065-5p | 162.72 | 0.24 | 0.667526658 | 0.999893527 |
| miR-6500-3p | 162.37 | -0.10 | 0.820362926 | 0.999893527 |
| miR-3184-5p | 162.06 | 0.27 | 0.49587937 | 0.999893527 |
| miR-3909-3p | 161.55 | -0.47 | 0.309301191 | 0.999893527 |
| miR-195-3p | 161.32 | -0.18 | 0.714942286 | 0.999893527 |
| miR-495-3p | 161.30 | 0.95 | 0.741324823 | 0.999893527 |
| miR-1273c-3p | 160.32 | -0.21 | 0.679535987 | 0.999893527 |
| miR-744-3p | 159.87 | 0.14 | 0.756522713 | 0.999893527 |
| miR-622-5p | 159.59 | 0.10 | 0.879726144 | 0.999893527 |
| miR-1278-3p | 159.32 | -0.40 | 0.443800552 | 0.999893527 |
| miR-877-3p | 158.38 | -0.06 | 0.874734541 | 0.999893527 |
| miR-18b-5p | 157.02 | 0.21 | 0.683626727 | 0.999893527 |
| Leu_AAG_5half | 155.90 | 1.67 | 0.044108645 | 0.999893527 |
| miR-556-3p | 155.04 | -0.44 | 0.326550712 | 0.999893527 |
| miR-2110-3p | 154.87 | -0.23 | 0.719154569 | 0.999893527 |
| miR-346 | 152.62 | 0.07 | 0.927799523 | 0.999893527 |
| Thr_CGT_leader | 152.30 | -0.56 | 0.467471419 | 0.999893527 |
| miR-5010-3p | 151.22 | -0.36 | 0.352193903 | 0.999893527 |
| miR-24-2-5p | 150.80 | -0.06 | 0.908593634 | 0.999893527 |
| Leu_CAA_tRF5 | 148.98 | 0.25 | 0.606703295 | 0.999893527 |
| miR-629-3p | 147.20 | -0.21 | 0.642679861 | 0.999893527 |
| miR-181b-1-3p | 145.29 | 0.24 | 0.672258317 | 0.999893527 |
| miR-6731-5p | 143.87 | 0.12 | 0.8842798 | 0.999893527 |
| miR-3945-3p | 141.61 | 0.27 | 0.773550428 | 0.999893527 |
| miR-378c-5p | 141.19 | -0.27 | 0.595796958 | 0.999893527 |
| miR-30b-3p | 141.16 | 0.47 | 0.471823093 | 0.999893527 |
| Gln_CTG_tRF5 | 139.55 | 0.45 | 0.321344415 | 0.999893527 |
| let-7d-3p | 139.49 | 0.22 | 0.789660426 | 0.999893527 |
| miR-134-5p | 138.76 | 0.92 | 0.690378675 | 0.999893527 |

| | | | | |
|---|---|---|---|---|
| miR-1285-1-5p | 137.17 | -0.38 | 0.624435564 | 0.999893527 |
| miR-335-3p | 136.90 | 0.68 | 0.7592773 | 0.999893527 |
| miR-491-5p | 135.52 | 0.11 | 0.821216663 | 0.999893527 |
| miR-1911-3p | 135.18 | -0.44 | 0.351914579 | 0.999893527 |
| Gly_TCC_tRF5 | 134.61 | 1.10 | 0.152243023 | 0.999893527 |
| miR-1292-5p | 134.23 | -0.19 | 0.680113376 | 0.999893527 |
| miR-551b-3p | 133.97 | -0.10 | 0.856931672 | 0.999893527 |
| miR-3176-3p | 133.26 | -0.30 | 0.466615054 | 0.999893527 |
| miR-185-3p | 133.15 | -0.31 | 0.473845947 | 0.999893527 |
| miR-628-3p | 132.45 | 0.67 | 0.266143371 | 0.999893527 |
| miR-548b-5p | 130.49 | 0.22 | 0.591413532 | 0.999893527 |
| miR-4773-2-5p | 130.39 | -0.12 | 0.871595066 | 0.999893527 |
| miR-3150a-5p | 129.04 | -0.13 | 0.901027686 | 0.999893527 |
| miR-1290-3p | 128.96 | 0.90 | 0.399292057 | 0.999893527 |
| miR-378f-3p | 127.22 | -0.91 | 0.080685023 | 0.999893527 |
| miR-1843-3p | 126.29 | 0.12 | 0.839310267 | 0.999893527 |
| miR-1255a | 125.14 | -1.06 | 0.168501321 | 0.999893527 |
| miR-9903-3p | 123.05 | -0.05 | 0.924239778 | 0.999893527 |
| miR-1-2-3p | 122.89 | 0.18 | 0.684662401 | 0.999893527 |
| miR-6817-3p | 121.89 | 0.47 | 0.622304473 | 0.999893527 |
| miR-191-3p | 121.69 | -0.23 | 0.642654291 | 0.999893527 |
| miR-548u-3p | 120.12 | -0.73 | 0.309467951 | 0.999893527 |
| miR-1285-1-3p | 119.91 | -0.24 | 0.650967888 | 0.999893527 |
| miR-6127-3p | 119.87 | -0.15 | 0.82227449 | 0.999893527 |
| miR-188-5p | 118.88 | 0.55 | 0.275084409 | 0.999893527 |
| miR-10400-3p | 118.20 | -0.13 | 0.885557029 | 0.999893527 |
| miR-3144-3p | 118.20 | -0.19 | 0.755827096 | 0.999893527 |
| miR-1250-5p | 117.36 | -0.06 | 0.893167223 | 0.999893527 |
| miR-16-1-3p | 116.46 | 0.11 | 0.860750872 | 0.999893527 |
| miR-548ah-3p | 116.25 | -0.81 | 0.206387892 | 0.999893527 |
| miR-199b-3p | 115.81 | -0.09 | 0.892045296 | 0.999893527 |
| miR-651-5p | 115.59 | -0.29 | 0.634732811 | 0.999893527 |
| miR-7-2-5p | 115.09 | -0.21 | 0.709477192 | 0.999893527 |
| miR-4473 | 113.28 | -0.27 | 0.698129188 | 0.999893527 |
| miR-3170-5p | 113.25 | -0.44 | 0.384517524 | 0.999893527 |
| miR-152-5p | 112.36 | -0.17 | 0.853394313 | 0.999893527 |
| Ser_GCT_tRF5 | 109.31 | 1.03 | 0.200187288 | 0.999893527 |
| miR-4661-5p | 109.04 | -0.72 | 0.220158395 | 0.999893527 |
| miR-451a-5p | 107.67 | 0.74 | 0.624978253 | 0.999893527 |
| miR-1299 | 106.56 | -0.75 | 0.393641357 | 0.999893527 |
| miR-6827-3p | 106.19 | -0.73 | 0.239466819 | 0.999893527 |
| miR-4484-3p | 105.85 | 0.93 | 0.132110975 | 0.999893527 |
| miR-766-3p | 105.43 | 0.05 | 0.901551978 | 0.999893527 |
| miR-619-5p | 104.55 | -0.07 | 0.935998939 | 0.999893527 |
| miR-1299-3p | 104.36 | -0.50 | 0.424292011 | 0.999893527 |
| Ile_GAT_tRF5 | 102.87 | -0.20 | 0.782575998 | 0.999893527 |
| miR-3177-3p | 102.54 | -0.18 | 0.68843055 | 0.999893527 |
| miR-3615 | 101.39 | -0.39 | 0.414340318 | 0.999893527 |
| miR-485-3p | 100.55 | 0.50 | 0.857820988 | 0.999893527 |
| miR-12136-3p | 100.49 | 0.34 | 0.442441852 | 0.999893527 |
| miR-3173-5p | 100.19 | -0.01 | 0.989313146 | 0.999893527 |
| miR-410-5p | 100.08 | -0.28 | 0.714505583 | 0.999893527 |
| miR-548h-5p | 99.83 | 0.00 | 0.998507427 | 0.999893527 |
| miR-30c-1-3p | 99.34 | 0.07 | 0.895764042 | 0.999893527 |
| miR-365a-5p | 98.27 | -0.31 | 0.542152873 | 0.999893527 |
| Leu_TAG_tRF5 | 97.69 | 0.66 | 0.37538133 | 0.999893527 |
| miR-653-5p | 96.00 | -0.51 | 0.349357379 | 0.999893527 |
| miR-128-1-5p | 95.97 | 0.07 | 0.891383285 | 0.999893527 |
| miR-4421-3p | 95.96 | -0.54 | 0.24850739 | 0.999893527 |
| miR-4321-3p | 94.99 | 0.42 | 0.410566843 | 0.999893527 |
| miR-1277-5p | 94.92 | 0.24 | 0.576151275 | 0.999893527 |
| miR-125a-3p | 93.87 | 0.06 | 0.915421602 | 0.999893527 |

| | | | | |
|---|---|---|---|---|
| miR-499a-5p | 92.59 | -0.09 | 0.854110248 | 0.999893527 |
| miR-486-2-5p | 89.30 | -0.45 | 0.352867607 | 0.999893527 |
| miR-3157-3p | 88.68 | -0.16 | 0.76602268 | 0.999893527 |
| miR-452-5p | 88.46 | -0.13 | 0.906173704 | 0.999893527 |
| miR-10525-3p | 88.25 | -0.55 | 0.437441464 | 0.999893527 |
| miR-411-3p | 87.67 | 0.63 | 0.816813384 | 0.999893527 |
| Lys_CTT_leader | 86.94 | -0.44 | 0.439674743 | 0.999893527 |
| miR-33a-5p | 86.73 | 0.05 | 0.92033738 | 0.999893527 |
| miR-556-5p | 86.33 | 0.12 | 0.786754584 | 0.999893527 |
| His_GTG_tRF3 | 85.47 | 0.36 | 0.624813089 | 0.999893527 |
| Asn_GTT_leader | 85.13 | -0.53 | 0.582349014 | 0.999893527 |
| miR-548ay-5p | 85.07 | 0.08 | 0.878697335 | 0.999893527 |
| miR-548j-5p | 84.74 | 0.07 | 0.891477256 | 0.999893527 |
| miR-342-5p | 84.23 | -0.10 | 0.836599235 | 0.999893527 |
| Leu_CAG_tRF1 | 84.08 | 0.32 | 0.76799105 | 0.999893527 |
| miR-6715b-3p | 84.04 | -0.10 | 0.839713841 | 0.999893527 |
| miR-301b-5p | 83.99 | -0.22 | 0.748323812 | 0.999893527 |
| miR-21-3p | 83.12 | -0.54 | 0.244170144 | 0.999893527 |
| miR-346-5p | 82.20 | -0.26 | 0.731817868 | 0.999893527 |
| miR-182-3p | 81.87 | -0.11 | 0.833562709 | 0.999893527 |
| miR-26a-2-3p | 81.12 | -0.46 | 0.377011315 | 0.999893527 |
| miR-4521-3p | 80.71 | 0.24 | 0.705743902 | 0.999893527 |
| miR-548au-5p | 80.64 | 0.20 | 0.662309707 | 0.999893527 |
| miR-142-5p | 78.61 | 0.62 | 0.580058956 | 0.999893527 |
| miR-3155a-5p | 78.10 | -0.60 | 0.204501289 | 0.999893527 |
| miR-625-5p | 77.95 | 0.29 | 0.506203837 | 0.999893527 |
| miR-26b-3p | 77.56 | -0.11 | 0.883288527 | 0.999893527 |
| miR-4473-5p | 77.26 | -0.01 | 0.991713727 | 0.999893527 |
| miR-199a-5p | 76.50 | 0.14 | 0.858837282 | 0.999893527 |
| miR-196b-3p | 76.23 | 0.07 | 0.876921367 | 0.999893527 |
| miR-624-5p | 76.07 | -0.01 | 0.985563361 | 0.999893527 |
| miR-1291-5p | 75.74 | -0.17 | 0.694689558 | 0.999893527 |
| miR-581-5p | 75.73 | -1.21 | 0.204057174 | 0.999893527 |
| miR-3941-5p | 75.58 | 0.36 | 0.477449306 | 0.999893527 |
| miR-219a-1-3p | 75.51 | -0.33 | 0.654700577 | 0.999893527 |
| miR-1908-5p | 75.17 | 0.13 | 0.873232821 | 0.999893527 |
| Ile_TAT_tRF3 | 74.60 | 0.45 | 0.37248414 | 0.999893527 |
| miR-641-3p | 74.57 | -0.12 | 0.851791758 | 0.999893527 |
| miR-4467-5p | 74.22 | -0.23 | 0.783216473 | 0.999893527 |
| Val_TAC_tRF3 | 74.07 | 0.17 | 0.705404295 | 0.999893527 |
| miR-548t-3p | 74.04 | 0.22 | 0.762047676 | 0.999893527 |
| miR-6747-3p | 73.79 | 0.37 | 0.409910744 | 0.999893527 |
| miR-1295a-3p | 72.38 | -0.11 | 0.831362006 | 0.999893527 |
| miR-1910-5p | 72.03 | 0.08 | 0.936090722 | 0.999893527 |
| miR-502-5p | 71.42 | -0.03 | 0.953552799 | 0.999893527 |
| miR-548k-5p | 70.75 | -0.30 | 0.631583274 | 0.999893527 |
| miR-7704-5p | 70.19 | 0.54 | 0.607294058 | 0.999893527 |
| miR-31-3p | 69.76 | 0.12 | 0.875031531 | 0.999893527 |
| miR-3682-3p | 69.58 | 0.24 | 0.677077487 | 0.999893527 |
| miR-486-3p | 69.58 | -0.49 | 0.667965489 | 0.999893527 |
| miR-3910-1-5p | 69.28 | -0.11 | 0.83088904 | 0.999893527 |
| miR-1291-3p | 69.21 | -1.55 | 0.089985824 | 0.999893527 |
| miR-585-3p | 68.42 | -0.23 | 0.599675572 | 0.999893527 |
| miR-619-3p | 68.18 | -0.61 | 0.602704648 | 0.999893527 |
| Pro_CGG_tRF1 | 68.14 | -0.69 | 0.536881691 | 0.999893527 |
| miR-1295a | 67.92 | -0.18 | 0.729507975 | 0.999893527 |
| miR-1285-3p | 67.49 | -0.33 | 0.642555946 | 0.999893527 |
| miR-548h-5-5p | 67.28 | 0.17 | 0.69350957 | 0.999893527 |
| miR-3928-3p | 66.85 | -0.45 | 0.363208882 | 0.999893527 |
| miR-548ae-2-5p | 65.99 | -0.02 | 0.977838433 | 0.999893527 |
| miR-101-1-3p | 64.92 | -0.09 | 0.879083995 | 0.999893527 |
| miR-6797-3p | 64.85 | 0.19 | 0.721646894 | 0.999893527 |

| | | | | |
|---|---|---|---|---|
| miR-4286 | 64.61 | -0.16 | 0.749420524 | 0.999893527 |
| miR-2355-3p | 64.36 | -0.33 | 0.478387538 | 0.999893527 |
| miR-589-3p | 63.54 | 0.36 | 0.390714415 | 0.999893527 |
| miR-183-3p | 62.65 | 0.01 | 0.991123134 | 0.999893527 |
| Ser_AGA_misc | 62.32 | 0.15 | 0.846855547 | 0.999893527 |
| miR-1249-3p | 61.30 | -0.11 | 0.864757395 | 0.999893527 |
| miR-3199-1-5p | 61.07 | 0.90 | 0.306009596 | 0.999893527 |
| miR-1287-5p | 60.92 | -0.47 | 0.296478852 | 0.999893527 |
| miR-190a-5p | 60.90 | 0.17 | 0.735710818 | 0.999893527 |
| miR-627-3p | 60.87 | 0.23 | 0.704961549 | 0.999893527 |
| miR-8086-3p | 60.81 | 0.17 | 0.825508951 | 0.999893527 |
| miR-345-3p | 60.77 | 0.44 | 0.612886626 | 0.999893527 |
| Met_CAT_tRF3 | 60.39 | 1.01 | 0.073540096 | 0.999893527 |
| miR-580-3p | 60.18 | 0.38 | 0.541200589 | 0.999893527 |
| Ser_CGA_misc | 60.01 | 0.27 | 0.57798595 | 0.999893527 |
| miR-449a | 59.70 | -0.05 | 0.910708316 | 0.999893527 |
| miR-99a-3p | 59.58 | -0.03 | 0.963904335 | 0.999893527 |
| miR-16-2-5p | 59.51 | 0.54 | 0.3547507 | 0.999893527 |
| miR-181b-3p | 59.51 | -0.03 | 0.965286191 | 0.999893527 |
| miR-378g | 59.25 | -0.05 | 0.928662535 | 0.999893527 |
| miR-548h-3-3p | 58.12 | 0.01 | 0.98169832 | 0.999893527 |
| Ile_GAT_misc | 57.60 | 0.15 | 0.792353982 | 0.999893527 |
| miR-551b-5p | 57.30 | -0.04 | 0.956762034 | 0.999893527 |
| miR-3193-5p | 57.27 | -0.01 | 0.987590451 | 0.999893527 |
| miR-151b | 57.22 | -0.36 | 0.743159211 | 0.999893527 |
| miR-450a-1-3p | 56.88 | 0.57 | 0.487801222 | 0.999893527 |
| miR-1255a-5p | 56.61 | -0.70 | 0.328953356 | 0.999893527 |
| miR-1298-5p | 56.61 | -1.09 | 0.126764984 | 0.999893527 |
| miR-4786-3p | 56.38 | 0.02 | 0.962378941 | 0.999893527 |
| miR-876-5p | 55.93 | 0.12 | 0.818652441 | 0.999893527 |
| miR-212-3p | 55.81 | -0.35 | 0.461935984 | 0.999893527 |
| miR-548ar-3p | 55.80 | 0.03 | 0.951079684 | 0.999893527 |
| miR-4273-5p | 55.62 | 0.10 | 0.901313867 | 0.999893527 |
| Arg_ACG_3half | 55.58 | 1.51 | 0.174858637 | 0.999893527 |
| miR-5585-3p | 55.12 | 0.01 | 0.993031226 | 0.999893527 |
| miR-579-5p | 54.84 | -0.19 | 0.684455002 | 0.999893527 |
| miR-6868-3p | 54.78 | -0.33 | 0.612067343 | 0.999893527 |
| miR-3929-3p | 54.61 | 0.23 | 0.728796121 | 0.999893527 |
| Leu_TAA_tRF5 | 53.96 | 0.85 | 0.24791824 | 0.999893527 |
| let-7e-3p | 53.96 | -0.42 | 0.393417338 | 0.999893527 |
| miR-942-3p | 52.79 | 0.33 | 0.626557178 | 0.999893527 |
| miR-550a-3p | 52.71 | 0.10 | 0.839326794 | 0.999893527 |
| miR-3143-5p | 52.68 | -0.42 | 0.647535418 | 0.999893527 |
| miR-138-5p | 52.39 | -0.23 | 0.728151726 | 0.999893527 |
| miR-5699-5p | 52.37 | 0.55 | 0.367799184 | 0.999893527 |
| Thr_AGT_leader | 51.95 | -0.57 | 0.433602507 | 0.999893527 |
| miR-561-5p | 51.75 | -0.65 | 0.203297062 | 0.999893527 |
| miR-326 | 50.92 | -0.60 | 0.245066117 | 0.999893527 |
| miR-3907-3p | 50.89 | -0.59 | 0.595797377 | 0.999893527 |
| miR-5684-3p | 50.59 | -0.54 | 0.568657825 | 0.999893527 |
| miR-548am-5p | 50.47 | -0.30 | 0.537201419 | 0.999893527 |
| miR-3157-5p | 50.41 | 0.05 | 0.93853436 | 0.999893527 |
| Arg_CCG_tRF3 | 49.87 | 0.45 | 0.373975586 | 0.999893527 |
| miR-3190-3p | 49.64 | -0.23 | 0.711302003 | 0.999893527 |
| miR-6843-5p | 49.62 | 0.47 | 0.700056962 | 0.999893527 |
| miR-659-5p | 49.60 | -0.72 | 0.309205956 | 0.999893527 |
| let-7a-1-3p | 49.28 | 0.07 | 0.907276229 | 0.999893527 |
| miR-548ay-3p | 49.21 | 0.23 | 0.679222031 | 0.999893527 |
| miR-548b-3p | 49.19 | 0.06 | 0.907090085 | 0.999893527 |
| miR-431-5p | 49.12 | 0.47 | 0.731722114 | 0.999893527 |
| miR-2681-3p | 48.98 | -0.06 | 0.946527401 | 0.999893527 |
| miR-7851-3p | 48.84 | 0.05 | 0.92969965 | 0.999893527 |

| | | | | |
|---|---|---|---|---|
| miR-4284-5p | 48.59 | 0.67 | 0.503965353 | 0.999893527 |
| miR-6720-5p | 48.51 | -0.16 | 0.815800245 | 0.999893527 |
| miR-9903 | 48.14 | -0.74 | 0.219038447 | 0.999893527 |
| miR-6724-4-3p | 47.60 | -0.61 | 0.473037484 | 0.999893527 |
| miR-29b-2-5p | 47.07 | 0.14 | 0.855364376 | 0.999893527 |
| Ala_AGC_tRF3 | 46.87 | 0.13 | 0.805675528 | 0.999893527 |
| miR-590-5p | 46.78 | 0.02 | 0.96945727 | 0.999893527 |
| miR-135b-5p | 46.70 | 0.10 | 0.880122339 | 0.999893527 |
| miR-96-3p | 46.58 | -0.58 | 0.461709318 | 0.999893527 |
| miR-1257-5p | 46.53 | -0.88 | 0.142408498 | 0.999893527 |
| miR-4485-3p | 46.52 | 0.15 | 0.786342392 | 0.999893527 |
| miR-500b-3p | 46.42 | -0.86 | 0.063321062 | 0.999893527 |
| miR-5582-3p | 46.14 | 0.01 | 0.99183744 | 0.999893527 |
| Pro_AGG_tRF3 | 45.93 | -0.04 | 0.972737061 | 0.999893527 |
| miR-4652-5p | 45.81 | -0.37 | 0.607937614 | 0.999893527 |
| miR-378i | 45.71 | -0.27 | 0.590066182 | 0.999893527 |
| miR-1343-3p | 45.12 | -0.66 | 0.320160909 | 0.999893527 |
| miR-125b-1-5p | 45.03 | -0.43 | 0.552521866 | 0.999893527 |
| miR-505-5p | 44.65 | 0.19 | 0.695663876 | 0.999893527 |
| miR-1266-5p | 44.60 | 0.01 | 0.992940219 | 0.999893527 |
| miR-3662 | 44.58 | -0.86 | 0.204240756 | 0.999893527 |
| miR-24-1-5p | 44.57 | -0.21 | 0.640345563 | 0.999893527 |
| miR-4517-5p | 44.38 | 0.19 | 0.776207484 | 0.999893527 |
| miR-1268a-5p | 44.19 | 0.03 | 0.972848937 | 0.999893527 |
| miR-147b-3p | 44.19 | -0.17 | 0.765788225 | 0.999893527 |
| let-7c-3p | 44.15 | -0.42 | 0.565115526 | 0.999893527 |
| miR-550a-3-5p | 43.95 | -0.29 | 0.572327889 | 0.999893527 |
| miR-1262-5p | 43.31 | 0.16 | 0.863087345 | 0.999893527 |
| miR-199a-2-5p | 42.82 | 0.02 | 0.976168553 | 0.999893527 |
| miR-23b-5p | 42.82 | 0.03 | 0.953194332 | 0.999893527 |
| miR-155-5p | 42.79 | 0.29 | 0.747469051 | 0.999893527 |
| miR-548ag-2-5p | 41.96 | -0.56 | 0.226647248 | 0.999893527 |
| miR-5699-3p | 41.48 | 0.15 | 0.770344038 | 0.999893527 |
| miR-548f-5-3p | 41.20 | -0.82 | 0.291022595 | 0.999893527 |
| miR-3127-5p | 40.66 | 0.35 | 0.568849608 | 0.999893527 |
| miR-653-3p | 40.61 | 0.07 | 0.942343796 | 0.999893527 |
| miR-193a-3p | 40.45 | 0.17 | 0.719459045 | 0.999893527 |
| miR-4313-3p | 40.40 | -0.03 | 0.958382334 | 0.999893527 |
| miR-1275 | 40.07 | 0.49 | 0.30311719 | 0.999893527 |
| miR-4659a-3p | 39.99 | -0.39 | 0.430283963 | 0.999893527 |
| miR-4531-3p | 39.70 | 0.58 | 0.464217975 | 0.999893527 |
| Ala_TGC_tRF3 | 39.67 | 0.12 | 0.814005546 | 0.999893527 |
| miR-192-3p | 39.44 | -0.32 | 0.541112213 | 0.999893527 |
| miR-450a-2-3p | 39.28 | -0.13 | 0.800778024 | 0.999893527 |
| miR-7851-5p | 39.26 | 0.30 | 0.676008012 | 0.999893527 |
| miR-1468-5p | 39.07 | -0.56 | 0.330041847 | 0.999893527 |
| miR-4448-3p | 39.05 | -0.18 | 0.712474047 | 0.999893527 |
| miR-557-3p | 38.99 | -0.20 | 0.689335262 | 0.999893527 |
| miR-4306-3p | 38.95 | -0.54 | 0.613404135 | 0.999893527 |
| let-7a-1-5p | 38.94 | 0.35 | 0.639838347 | 0.999893527 |
| miR-378f | 38.93 | -0.22 | 0.708623149 | 0.999893527 |
| miR-4784-3p | 38.77 | -1.07 | 0.315924961 | 0.999893527 |
| miR-4677-5p | 38.73 | -0.56 | 0.425099581 | 0.999893527 |
| miR-346-3p | 38.71 | -0.31 | 0.599785731 | 0.999893527 |
| miR-449a-5p | 38.60 | 0.18 | 0.74181427 | 0.999893527 |
| Phe_GAA_leader | 38.44 | -0.19 | 0.865495649 | 0.999893527 |
| miR-12114-5p | 37.62 | -0.49 | 0.428337426 | 0.999893527 |
| miR-323a-3p | 37.57 | 0.45 | 0.857929375 | 0.999893527 |
| miR-548d-5p | 37.55 | 0.13 | 0.816040621 | 0.999893527 |
| miR-190b-5p | 37.50 | 0.27 | 0.632545088 | 0.999893527 |
| miR-326-3p | 37.04 | -0.55 | 0.483363318 | 0.999893527 |
| miR-597-3p | 36.98 | -0.31 | 0.55316109 | 0.999893527 |

| | | | | |
|---|---|---|---|---|
| miR-6832-5p | 36.96 | -0.05 | 0.93503155 | 0.999893527 |
| miR-504-5p | 36.67 | -0.10 | 0.906921959 | 0.999893527 |
| miR-548d-2-5p | 36.62 | -0.41 | 0.470676648 | 0.999893527 |
| miR-1252-5p | 36.62 | -0.04 | 0.947156924 | 0.999893527 |
| miR-365b-5p | 36.48 | 0.07 | 0.912753664 | 0.999893527 |
| miR-1255a-3p | 36.06 | -0.32 | 0.531912807 | 0.999893527 |
| miR-4430-3p | 35.83 | 0.24 | 0.790414805 | 0.999893527 |
| miR-3912-3p | 35.75 | 0.20 | 0.709675567 | 0.999893527 |
| miR-10399-3p | 35.69 | -0.58 | 0.341816053 | 0.999893527 |
| miR-193b-5p | 35.68 | 0.11 | 0.846141324 | 0.999893527 |
| Pro_TGG_tRF3 | 35.67 | -0.02 | 0.982152298 | 0.999893527 |
| miR-218-1-3p | 35.62 | 0.34 | 0.632214446 | 0.999893527 |
| miR-4742-3p | 35.31 | 0.15 | 0.764159633 | 0.999893527 |
| miR-1271-3p | 35.21 | -0.18 | 0.734007901 | 0.999893527 |
| miR-873-3p | 34.97 | -0.26 | 0.690361396 | 0.999893527 |
| miR-138-1-5p | 34.96 | 0.21 | 0.816888102 | 0.999893527 |
| miR-921-5p | 34.82 | -0.10 | 0.926414065 | 0.999893527 |
| miR-3648-2-3p | 34.80 | -0.27 | 0.585475294 | 0.999893527 |
| miR-7150-5p | 34.80 | 0.09 | 0.92070872 | 0.999893527 |
| miR-4695-3p | 34.65 | 0.20 | 0.744770607 | 0.999893527 |
| miR-6505-5p | 34.52 | 0.06 | 0.898889541 | 0.999893527 |
| miR-133a-1-3p | 34.22 | -0.33 | 0.542296916 | 0.999893527 |
| miR-216b-5p | 33.91 | -0.06 | 0.909632332 | 0.999893527 |
| miR-4741-3p | 33.83 | -0.36 | 0.52608758 | 0.999893527 |
| miR-3691-5p | 33.73 | 0.14 | 0.854158014 | 0.999893527 |
| miR-325-5p | 33.34 | -0.18 | 0.761674537 | 0.999893527 |
| miR-550a-3-3p | 33.25 | -0.36 | 0.467371355 | 0.999893527 |
| miR-3127-3p | 33.19 | 0.07 | 0.924152997 | 0.999893527 |
| miR-320d-1-3p | 33.08 | 0.28 | 0.596830275 | 0.999893527 |
| miR-5001-5p | 33.01 | 0.16 | 0.802185359 | 0.999893527 |
| miR-146b-3p | 33.01 | -0.14 | 0.799833513 | 0.999893527 |
| miR-1276-5p | 32.99 | -0.28 | 0.62426044 | 0.999893527 |
| miR-548ag-1-5p | 32.98 | -0.10 | 0.861978487 | 0.999893527 |
| miR-548l | 32.93 | 0.22 | 0.749722897 | 0.999893527 |
| miR-548j-3p | 32.92 | -0.50 | 0.472901884 | 0.999893527 |
| miR-3929-5p | 32.71 | -0.37 | 0.513072503 | 0.999893527 |
| miR-503-3p | 32.52 | -0.03 | 0.96237633 | 0.999893527 |
| Ser_TGA_misc | 32.45 | 0.59 | 0.364376519 | 0.999893527 |
| Arg_CCT_tRF3 | 32.30 | 0.62 | 0.414084153 | 0.999893527 |
| miR-5708-3p | 32.24 | 0.33 | 0.698988338 | 0.999893527 |
| miR-4451-3p | 32.24 | -0.39 | 0.612249919 | 0.999893527 |
| miR-1199-5p | 32.19 | 0.11 | 0.858094616 | 0.999893527 |
| Pro_AGG_tRF5 | 31.99 | 0.83 | 0.155299628 | 0.999893527 |
| miR-142-3p | 31.86 | 0.80 | 0.451504582 | 0.999893527 |
| miR-3934-5p | 31.82 | 0.09 | 0.852587878 | 0.999893527 |
| miR-1288-3p | 31.63 | -1.15 | 0.148625963 | 0.999893527 |
| miR-3674-5p | 31.55 | -1.41 | 0.223214977 | 0.999893527 |
| miR-320c | 31.53 | 0.31 | 0.604822737 | 0.999893527 |
| miR-3944-3p | 31.47 | -0.44 | 0.469164284 | 0.999893527 |
| miR-378g-5p | 31.44 | -1.05 | 0.167733572 | 0.999893527 |
| miR-362-3p | 31.38 | 0.04 | 0.938330245 | 0.999893527 |
| miR-3913-2-5p | 31.22 | -1.33 | 0.066921981 | 0.999893527 |
| miR-410-3p | 31.07 | 0.23 | 0.925474547 | 0.999893527 |
| miR-1234-5p | 30.81 | 0.81 | 0.528437838 | 0.999893527 |
| miR-181b-2-3p | 30.79 | -0.25 | 0.650677165 | 0.999893527 |
| miR-1262 | 30.67 | 0.52 | 0.568648323 | 0.999893527 |
| miR-579-3p | 30.66 | 0.03 | 0.967438892 | 0.999893527 |
| miR-3677-3p | 30.65 | -0.43 | 0.410170065 | 0.999893527 |
| Asp_GTC_leader | 30.60 | 0.15 | 0.804677236 | 0.999893527 |
| miR-548ba-3p | 30.55 | -0.20 | 0.806999963 | 0.999893527 |
| miR-548l-5p | 30.55 | -0.22 | 0.683441779 | 0.999893527 |
| miR-5009-5p | 30.41 | -1.26 | 0.046868751 | 0.999893527 |

| | | | | |
|---|---|---|---|---|
| miR-6872-3p | 30.26 | 0.26 | 0.649783565 | 0.999893527 |
| miR-6739-3p | 30.25 | -0.13 | 0.833667554 | 0.999893527 |
| miR-26a-1-5p | 30.24 | -0.04 | 0.941471766 | 0.999893527 |
| miR-642a-5p | 30.06 | 0.40 | 0.638082837 | 0.999893527 |
| miR-3648-2-5p | 29.93 | 0.12 | 0.842679713 | 0.999893527 |
| miR-3065-3p | 29.81 | 0.26 | 0.67694196 | 0.999893527 |
| Leu_CAA_tRF1 | 29.72 | -0.28 | 0.779828118 | 0.999893527 |
| miR-588-3p | 29.72 | -0.87 | 0.11285921 | 0.999893527 |
| miR-6516-5p | 29.62 | 0.12 | 0.848150003 | 0.999893527 |
| miR-335-5p | 29.42 | 0.75 | 0.598688109 | 0.999893527 |
| miR-1273h-3p | 29.41 | -0.78 | 0.395598031 | 0.999893527 |
| miR-548n-5p | 29.36 | -0.32 | 0.663462655 | 0.999893527 |
| Leu_TAA_tRF3 | 29.30 | 0.17 | 0.816676459 | 0.999893527 |
| miR-6866-5p | 29.24 | -0.67 | 0.382289361 | 0.999893527 |
| miR-548az-5p | 29.22 | -0.38 | 0.526670564 | 0.999893527 |
| miR-3136-5p | 29.09 | -0.09 | 0.8828417 | 0.999893527 |
| miR-548ag-2-3p | 29.02 | -0.26 | 0.761528042 | 0.999893527 |
| miR-6724-4-5p | 28.81 | -0.41 | 0.506040706 | 0.999893527 |
| miR-122-5p | 28.38 | 1.61 | 0.142033419 | 0.999893527 |
| miR-3665-3p | 28.22 | -0.25 | 0.776647212 | 0.999893527 |
| miR-483-3p | 28.04 | -0.14 | 0.790926304 | 0.999893527 |
| miR-5696-3p | 27.95 | 0.31 | 0.580475111 | 0.999893527 |
| Arg_CCG_leader | 27.93 | -0.51 | 0.450914602 | 0.999893527 |
| miR-3187-3p | 27.88 | -0.10 | 0.854075486 | 0.999893527 |
| miR-6505-3p | 27.67 | 0.24 | 0.695969566 | 0.999893527 |
| miR-1305-3p | 27.65 | 0.06 | 0.93198292 | 0.999893527 |
| miR-1269a-5p | 27.48 | 0.06 | 0.938275101 | 0.999893527 |
| miR-551a | 27.42 | 0.22 | 0.750456957 | 0.999893527 |
| miR-6783-3p | 27.21 | -0.47 | 0.41923067 | 0.999893527 |
| miR-764-3p | 27.07 | -0.15 | 0.799405881 | 0.999893527 |
| miR-592-5p | 27.02 | 0.13 | 0.816223247 | 0.999893527 |
| miR-10401-3p | 26.99 | -0.30 | 0.748877861 | 0.999893527 |
| Pro_AGG_leader | 26.87 | 0.12 | 0.884682244 | 0.999893527 |
| miR-940-3p | 26.84 | -0.03 | 0.959508925 | 0.999893527 |
| miR-494-3p | 26.81 | 0.25 | 0.861709893 | 0.999893527 |
| miR-1273h-5p | 26.73 | -0.26 | 0.73167061 | 0.999893527 |
| miR-8485-3p | 26.50 | 0.60 | 0.610213341 | 0.999893527 |
| miR-452-3p | 26.42 | -0.37 | 0.619293746 | 0.999893527 |
| miR-767-5p | 26.32 | 0.75 | 0.301199464 | 0.999893527 |
| miR-6842-3p | 26.31 | -0.58 | 0.380473091 | 0.999893527 |
| miR-10400-5p | 26.12 | 0.15 | 0.846331545 | 0.999893527 |
| miR-26a-1-3p | 26.04 | -0.18 | 0.733603885 | 0.999893527 |
| miR-4470-3p | 25.93 | 0.14 | 0.798448922 | 0.999893527 |
| miR-541-3p | 25.71 | 0.06 | 0.948283601 | 0.999893527 |
| miR-548u | 25.70 | -0.32 | 0.553501357 | 0.999893527 |
| miR-5685-3p | 25.68 | -0.97 | 0.168826978 | 0.999893527 |
| miR-382-3p | 25.67 | 0.76 | 0.629215882 | 0.999893527 |
| miR-4647-3p | 25.63 | -0.23 | 0.693673281 | 0.999893527 |
| miR-885-3p | 25.61 | 0.38 | 0.543876011 | 0.999893527 |
| miR-514a-3p | 25.56 | -1.13 | 0.221388999 | 0.999893527 |
| miR-766-5p | 25.49 | 0.41 | 0.439295305 | 0.999893527 |
| miR-5002-3p | 25.40 | 0.50 | 0.727239441 | 0.999893527 |
| miR-6515-3p | 25.39 | -0.14 | 0.798043055 | 0.999893527 |
| miR-4458-5p | 25.39 | 0.22 | 0.769167109 | 0.999893527 |
| miR-1303-5p | 25.36 | -0.33 | 0.580956935 | 0.999893527 |
| miR-548ac-5p | 25.06 | -1.15 | 0.197518911 | 0.999893527 |
| miR-6873-3p | 25.05 | 0.06 | 0.922653295 | 0.999893527 |
| miR-33b-5p | 25.01 | 0.14 | 0.846366142 | 0.999893527 |
| miR-219a-1-5p | 24.97 | -0.38 | 0.644130932 | 0.999893527 |
| miR-664-5p | 24.88 | 0.11 | 0.91192767 | 0.999893527 |
| miR-6869-5p | 24.85 | -0.17 | 0.794241585 | 0.999893527 |
| miR-6768-3p | 24.73 | -0.52 | 0.544031732 | 0.999893527 |

| | | | | |
|---|---|---|---|---|
| miR-1285-2-5p | 24.69 | 0.33 | 0.753655321 | 0.999893527 |
| miR-33b-3p | 24.65 | 0.35 | 0.606156283 | 0.999893527 |
| miR-1269a-3p | 24.60 | -0.40 | 0.452290761 | 0.999893527 |
| miR-548f-3-3p | 24.56 | 0.46 | 0.568473307 | 0.999893527 |
| Pro_TGG_tRF5 | 24.54 | 0.93 | 0.102694622 | 0.999893527 |
| miR-7846-3p | 24.46 | 0.55 | 0.655671941 | 0.999893527 |
| miR-548ax-3p | 24.33 | -0.64 | 0.404015204 | 0.999893527 |
| miR-2116-3p | 24.32 | -0.02 | 0.976537988 | 0.999893527 |
| miR-219b-5p | 24.27 | 0.17 | 0.760957919 | 0.999893527 |
| miR-1273c-5p | 24.13 | 0.12 | 0.85340392 | 0.999893527 |
| miR-369-5p | 24.06 | 0.63 | 0.613095532 | 0.999893527 |
| miR-2277-3p | 23.87 | -0.26 | 0.733750907 | 0.999893527 |
| miR-616-5p | 23.66 | 0.28 | 0.676475053 | 0.999893527 |
| miR-4496-3p | 23.63 | 0.69 | 0.283349734 | 0.999893527 |
| miR-664a-3p | 23.41 | 0.51 | 0.345923499 | 0.999893527 |
| miR-3609-3p | 23.38 | -0.05 | 0.93608279 | 0.999893527 |
| miR-548l-3p | 23.19 | -0.76 | 0.30071731 | 0.999893527 |
| Ser_AGA_leader | 23.16 | -0.67 | 0.645592506 | 0.999893527 |
| miR-6818-3p | 23.05 | 0.01 | 0.982955426 | 0.999893527 |
| miR-3684-3p | 22.88 | -0.16 | 0.790052103 | 0.999893527 |
| miR-500a-5p | 22.83 | 0.19 | 0.768856565 | 0.999893527 |
| miR-1268a-3p | 22.76 | 0.10 | 0.889374008 | 0.999893527 |
| Tyr_ATA_tRF3 | 22.63 | -0.59 | 0.473410731 | 0.999893527 |
| miR-204-5p | 22.56 | -0.21 | 0.742398231 | 0.999893527 |
| miR-449b-5p | 22.52 | -0.69 | 0.264085543 | 0.999893527 |
| miR-9903-5p | 22.52 | -0.25 | 0.637536337 | 0.999893527 |
| miR-3908-5p | 22.49 | -0.55 | 0.323809167 | 0.999893527 |
| miR-550a-5p | 22.43 | -0.22 | 0.784176113 | 0.999893527 |
| miR-383-5p | 22.39 | -0.14 | 0.83360245 | 0.999893527 |
| Gln_CTG_leader | 22.39 | 0.66 | 0.505355261 | 0.999893527 |
| miR-4757-5p | 22.12 | -0.12 | 0.92667464 | 0.999893527 |
| Ala_TGC_leader | 22.07 | -1.01 | 0.349320172 | 0.999893527 |
| miR-374c-5p | 22.07 | -0.36 | 0.512627815 | 0.999893527 |
| miR-3605-3p | 22.06 | -0.14 | 0.828533922 | 0.999893527 |
| miR-34a-3p | 22.00 | 0.11 | 0.869405318 | 0.999893527 |
| miR-3074-3p | 21.87 | -0.12 | 0.83263363 | 0.999893527 |
| iMet_CAT_leader | 21.86 | 0.10 | 0.929477261 | 0.999893527 |
| miR-1277-3p | 21.84 | 0.23 | 0.679770632 | 0.999893527 |
| miR-217-5p | 21.79 | -0.09 | 0.886886447 | 0.999893527 |
| miR-1224-5p | 21.74 | 0.18 | 0.852702049 | 0.999893527 |
| miR-548an-5p | 21.72 | 0.11 | 0.855755513 | 0.999893527 |
| miR-3133-5p | 21.71 | -1.01 | 0.101381926 | 0.999893527 |
| miR-4474-3p | 21.65 | -0.54 | 0.404180817 | 0.999893527 |
| miR-1248-5p | 21.64 | 0.26 | 0.717658308 | 0.999893527 |
| miR-1305 | 21.52 | 0.60 | 0.373095438 | 0.999893527 |
| miR-3143-3p | 21.36 | -0.94 | 0.100906048 | 0.999893527 |
| miR-5684-5p | 21.32 | 0.66 | 0.561087778 | 0.999893527 |
| miR-6857-3p | 21.28 | -0.34 | 0.610768856 | 0.999893527 |
| miR-5682-5p | 21.25 | 0.35 | 0.773893577 | 0.999893527 |
| miR-3651-3p | 21.24 | 1.61 | 0.062097477 | 0.999893527 |
| miR-6758-3p | 21.24 | -0.22 | 0.81629375 | 0.999893527 |
| miR-3616-5p | 21.22 | -0.18 | 0.749779924 | 0.999893527 |
| miR-2278-5p | 21.19 | 0.14 | 0.809778216 | 0.999893527 |
| miR-642a-3p | 21.16 | -0.53 | 0.442761056 | 0.999893527 |
| miR-605-3p | 21.04 | 0.88 | 0.228253324 | 0.999893527 |
| miR-6084-5p | 21.04 | 0.64 | 0.471788533 | 0.999893527 |
| miR-643-3p | 21.03 | 0.04 | 0.938328043 | 0.999893527 |
| miR-3909-5p | 20.99 | 0.60 | 0.573928243 | 0.999893527 |
| miR-574-5p | 20.95 | 0.53 | 0.387611197 | 0.999893527 |
| miR-215-3p | 20.95 | -0.66 | 0.36497306 | 0.999893527 |
| miR-3115 | 20.90 | 0.11 | 0.856830763 | 0.999893527 |
| miR-663a-3p | 20.88 | 0.46 | 0.480513269 | 0.999893527 |

| | | | | |
|---|---|---|---|---|
| miR-937-3p | 20.88 | -0.51 | 0.397188045 | 0.999893527 |
| miR-210-5p | 20.77 | 0.33 | 0.66123024 | 0.999893527 |
| miR-548aj-2-3p | 20.66 | 0.33 | 0.568259295 | 0.999893527 |
| miR-6882-5p | 20.57 | 0.08 | 0.900137982 | 0.999893527 |
| miR-548ag | 20.57 | -0.20 | 0.735701222 | 0.999893527 |
| miR-30c-2-5p | 20.54 | 0.83 | 0.347659253 | 0.999893527 |
| miR-149-3p | 20.50 | -0.06 | 0.922218497 | 0.999893527 |
| miR-4713-5p | 20.45 | 0.30 | 0.668890643 | 0.999893527 |
| miR-3135b-3p | 20.38 | 0.18 | 0.871340352 | 0.999893527 |
| miR-6507-5p | 20.38 | 0.48 | 0.439925791 | 0.999893527 |
| Glu_CTC_leader | 20.29 | 0.26 | 0.660697108 | 0.999893527 |
| miR-9899-5p | 20.16 | 0.52 | 0.731991765 | 0.999893527 |
| miR-1293-5p | 20.06 | -1.63 | 0.110514999 | 0.999893527 |
| miR-3662-3p | 19.96 | -0.27 | 0.645720032 | 0.999893527 |
| Pro_CGG_tRF3 | 19.95 | -0.02 | 0.986231313 | 0.999893527 |
| miR-660-3p | 19.90 | 0.06 | 0.921739444 | 0.999893527 |
| SeC_TCA_tRF5 | 19.75 | 0.40 | 0.584280233 | 0.999893527 |
| miR-4676-3p | 19.74 | -0.08 | 0.894882891 | 0.999893527 |
| miR-4762-3p | 19.61 | -0.35 | 0.547851621 | 0.999893527 |
| miR-4449-3p | 19.58 | 0.25 | 0.850259626 | 0.999893527 |
| miR-665-3p | 19.47 | 0.01 | 0.99600984 | 0.999893527 |
| miR-3679-5p | 19.43 | 0.08 | 0.897044196 | 0.999893527 |
| miR-627-5p | 19.41 | 0.47 | 0.482683824 | 0.999893527 |
| miR-139-3p | 19.39 | -0.61 | 0.30518273 | 0.999893527 |
| miR-573-5p | 19.24 | -0.15 | 0.821800693 | 0.999893527 |
| miR-181d-3p | 19.21 | -0.34 | 0.602609932 | 0.999893527 |
| miR-621-5p | 19.19 | -0.20 | 0.792231751 | 0.999893527 |
| miR-376a-2-3p | 19.16 | 0.65 | 0.675594676 | 0.999893527 |
| miR-592 | 19.12 | -0.21 | 0.761540209 | 0.999893527 |
| miR-548y-5p | 19.06 | 0.00 | 0.993535173 | 0.999893527 |
| Leu_AAG_3half | 18.95 | -0.45 | 0.877196473 | 0.999893527 |
| miR-3126-3p | 18.90 | -0.41 | 0.612085645 | 0.999893527 |
| Gln_TTG_3half | 18.88 | 2.36 | 0.054446716 | 0.999893527 |
| miR-1243-5p | 18.82 | 0.23 | 0.753326262 | 0.999893527 |
| miR-3922-3p | 18.78 | -0.74 | 0.247766381 | 0.999893527 |
| miR-6516-3p | 18.68 | 0.39 | 0.532984193 | 0.999893527 |
| Tyr_GTA_3half | 18.65 | -0.16 | 0.908777647 | 0.999893527 |
| miR-3166-5p | 18.58 | -0.39 | 0.624521184 | 0.999893527 |
| miR-548a-3-3p | 18.57 | 0.23 | 0.717412765 | 0.999893527 |
| miR-3611-5p | 18.57 | 0.54 | 0.348526314 | 0.999893527 |
| miR-3939-5p | 18.52 | 0.17 | 0.824884048 | 0.999893527 |
| miR-5091-5p | 18.52 | -0.71 | 0.297702587 | 0.999893527 |
| miR-3195-3p | 18.43 | 0.80 | 0.560360715 | 0.999893527 |
| miR-935 | 18.29 | 0.07 | 0.947486659 | 0.999893527 |
| miR-491-3p | 18.27 | -0.24 | 0.692792054 | 0.999893527 |
| miR-1322-3p | 18.25 | -0.30 | 0.595642354 | 0.999893527 |
| miR-129-2-3p | 18.06 | 0.24 | 0.710960008 | 0.999893527 |
| miR-7108-3p | 18.03 | 0.22 | 0.824122614 | 0.999893527 |
| miR-591-3p | 18.03 | -0.32 | 0.685631265 | 0.999893527 |
| miR-8485-5p | 18.02 | -0.15 | 0.793147424 | 0.999893527 |
| miR-136-3p | 17.96 | 0.84 | 0.588720546 | 0.999893527 |
| Asn_GTT_tRF5 | 17.94 | 0.46 | 0.526547131 | 0.999893527 |
| miR-147b-5p | 17.91 | 0.28 | 0.704129534 | 0.999893527 |
| miR-602-5p | 17.91 | -0.32 | 0.686699735 | 0.999893527 |
| miR-409-5p | 17.87 | 0.52 | 0.734078868 | 0.999893527 |
| miR-1538-3p | 17.83 | 0.15 | 0.815564184 | 0.999893527 |
| miR-6511b-1-3p | 17.80 | -0.29 | 0.734618961 | 0.999893527 |
| miR-3908-3p | 17.73 | -0.14 | 0.867917324 | 0.999893527 |
| miR-3940-3p | 17.63 | 0.17 | 0.792833802 | 0.999893527 |
| miR-3201-3p | 17.62 | 0.88 | 0.206830141 | 0.999893527 |
| miR-2110 | 17.62 | -0.25 | 0.68681926 | 0.999893527 |
| Arg_TCT_tRF5 | 17.51 | 0.98 | 0.231602862 | 0.999893527 |

| | | | | |
|---|---|---|---|---|
| miR-3179-4-3p | 17.49 | 0.48 | 0.495793767 | 0.999893527 |
| miR-4423-3p | 17.30 | -0.26 | 0.714551595 | 0.999893527 |
| miR-4663-3p | 17.27 | 0.69 | 0.295064225 | 0.999893527 |
| miR-6793-5p | 17.14 | 0.55 | 0.552583969 | 0.999893527 |
| miR-10523-5p | 17.04 | 0.16 | 0.818596172 | 0.999893527 |
| miR-4786-5p | 16.97 | -1.17 | 0.126528339 | 0.999893527 |
| miR-6126-3p | 16.97 | 0.04 | 0.952416946 | 0.999893527 |
| miR-4780-3p | 16.92 | 0.49 | 0.44574113 | 0.999893527 |
| miR-127-5p | 16.85 | 0.04 | 0.9650149 | 0.999893527 |
| miR-10394-5p | 16.76 | -0.09 | 0.878973255 | 0.999893527 |
| miR-664-3p | 16.67 | 0.57 | 0.390444922 | 0.999893527 |
| miR-4449-5p | 16.66 | -0.22 | 0.862599267 | 0.999893527 |
| miR-3620-3p | 16.63 | -0.51 | 0.480421546 | 0.999893527 |
| miR-338-3p | 16.62 | -0.05 | 0.958500528 | 0.999893527 |
| miR-380-3p | 16.58 | 0.67 | 0.620709855 | 0.999893527 |
| miR-5689-5p | 16.58 | -0.47 | 0.668897032 | 0.999893527 |
| miR-9901-5p | 16.47 | -0.15 | 0.811777384 | 0.999893527 |
| miR-5000-3p | 16.45 | 0.21 | 0.755458687 | 0.999893527 |
| Cys_GCA_leader | 16.43 | -0.98 | 0.416208717 | 0.999893527 |
| miR-578-3p | 16.39 | 0.06 | 0.924449735 | 0.999893527 |
| miR-1203-5p | 16.31 | 1.08 | 0.121860173 | 0.999893527 |
| miR-1268b-3p | 16.30 | 0.14 | 0.844648448 | 0.999893527 |
| miR-5697-5p | 16.27 | -0.10 | 0.878239077 | 0.999893527 |
| miR-1247-3p | 16.26 | 0.93 | 0.364297421 | 0.999893527 |
| Phe_GAA_tRF5 | 16.25 | 0.83 | 0.267715044 | 0.999893527 |
| miR-4784-5p | 16.22 | 0.12 | 0.870248287 | 0.999893527 |
| miR-548ar-5p | 16.22 | 0.44 | 0.516342351 | 0.999893527 |
| miR-4421 | 16.22 | -0.38 | 0.522690675 | 0.999893527 |
| miR-4309-3p | 16.21 | 0.44 | 0.546109683 | 0.999893527 |
| Ser_GCT_leader | 16.17 | -0.99 | 0.370469013 | 0.999893527 |
| Gln_TTG_tRF5 | 16.01 | 0.32 | 0.67448693 | 0.999893527 |
| miR-449a-3p | 16.01 | -0.78 | 0.457918961 | 0.999893527 |
| miR-4508-5p | 15.96 | 0.40 | 0.573298742 | 0.999893527 |
| miR-548an-3p | 15.86 | -0.75 | 0.301281302 | 0.999893527 |
| miR-764-5p | 15.79 | 0.36 | 0.617425653 | 0.999893527 |
| miR-6789-3p | 15.75 | -0.34 | 0.606385409 | 0.999893527 |
| miR-4277-3p | 15.67 | 0.79 | 0.400585036 | 0.999893527 |
| miR-4742-5p | 15.60 | 0.27 | 0.682955543 | 0.999893527 |
| miR-3180-3-5p | 15.57 | 0.37 | 0.681920609 | 0.999893527 |
| Cys_GCA_3half | 15.55 | 0.54 | 0.706796993 | 0.999893527 |
| miR-943-3p | 15.49 | -0.09 | 0.883866265 | 0.999893527 |
| miR-548d-1-3p | 15.43 | -0.04 | 0.965913344 | 0.999893527 |
| miR-3140-3p | 15.43 | 0.24 | 0.769390542 | 0.999893527 |
| miR-4665-5p | 15.37 | -0.72 | 0.284173483 | 0.999893527 |
| miR-320b | 15.18 | 0.13 | 0.893971508 | 0.999893527 |
| miR-5094-5p | 15.11 | -0.66 | 0.346264032 | 0.999893527 |
| miR-378d-2-5p | 15.11 | 0.34 | 0.627756 | 0.999893527 |
| miR-181a-1-5p | 15.09 | 0.06 | 0.923195542 | 0.999893527 |
| miR-3181-3p | 15.06 | 1.31 | 0.351572757 | 0.999893527 |
| miR-4635-3p | 14.99 | -0.88 | 0.496487269 | 0.999893527 |
| miR-548y-3p | 14.93 | 0.29 | 0.673412144 | 0.999893527 |
| miR-3665-5p | 14.87 | 1.01 | 0.162572839 | 0.999893527 |
| miR-511-5p | 14.85 | -0.10 | 0.87878917 | 0.999893527 |
| miR-145-3p | 14.84 | 0.54 | 0.72054609 | 0.999893527 |
| miR-153-3p | 14.77 | -0.28 | 0.775380783 | 0.999893527 |
| miR-138-2-3p | 14.77 | 1.47 | 0.069092401 | 0.999893527 |
| miR-1293-3p | 14.58 | -0.13 | 0.865456707 | 0.999893527 |
| miR-548q-5p | 14.54 | 0.59 | 0.387487548 | 0.999893527 |
| miR-760 | 14.51 | -0.75 | 0.243803808 | 0.999893527 |
| miR-6771-5p | 14.51 | -0.55 | 0.487947425 | 0.999893527 |
| miR-718-3p | 14.48 | -0.24 | 0.717175792 | 0.999893527 |
| miR-4778-3p | 14.41 | 0.74 | 0.365588629 | 0.999893527 |

| | | | | |
|---|---|---|---|---|
| miR-3938 | 14.38 | 0.31 | 0.686358932 | 0.999893527 |
| miR-548y | 14.34 | -0.77 | 0.234039568 | 0.999893527 |
| miR-676-5p | 14.34 | 0.18 | 0.777075367 | 0.999893527 |
| miR-3174-5p | 14.33 | -0.01 | 0.993302451 | 0.999893527 |
| miR-144-3p | 14.32 | 0.99 | 0.511674835 | 0.999893527 |
| miR-3139 | 14.30 | -0.25 | 0.719281376 | 0.999893527 |
| miR-3942-3p | 14.28 | -0.29 | 0.739245268 | 0.999893527 |
| miR-615-5p | 14.26 | 0.04 | 0.949111582 | 0.999893527 |
| miR-188-3p | 14.22 | -0.43 | 0.531778013 | 0.999893527 |
| miR-3128-5p | 14.17 | -0.35 | 0.607847126 | 0.999893527 |
| miR-1260b | 14.17 | 0.59 | 0.374132995 | 0.999893527 |
| miR-4430-5p | 14.11 | 0.02 | 0.981451827 | 0.999893527 |
| miR-577 | 14.10 | 0.34 | 0.696834907 | 0.999893527 |
| miR-15a-3p | 14.10 | -0.33 | 0.605978714 | 0.999893527 |
| miR-4301-5p | 14.09 | -0.70 | 0.295805542 | 0.999893527 |
| miR-3200-5p | 13.99 | 0.66 | 0.338312739 | 0.999893527 |
| miR-4296-5p | 13.98 | -0.10 | 0.888489354 | 0.999893527 |
| miR-101-2-5p | 13.93 | 0.20 | 0.837919127 | 0.999893527 |
| miR-4433b-5p | 13.92 | -0.31 | 0.763257646 | 0.999893527 |
| Arg_TCT_tRF1 | 13.92 | 0.21 | 0.821229713 | 0.999893527 |
| miR-5581-3p | 13.92 | -0.28 | 0.722130169 | 0.999893527 |
| miR-1247-5p | 13.89 | 0.08 | 0.92821856 | 0.999893527 |
| miR-6733-5p | 13.84 | 0.12 | 0.866413846 | 0.999893527 |
| miR-1976-3p | 13.84 | 0.56 | 0.515023398 | 0.999893527 |
| Pro_CGG_tRF5 | 13.81 | 0.80 | 0.24330318 | 0.999893527 |
| miR-1226-5p | 13.73 | -0.07 | 0.913241921 | 0.999893527 |
| miR-6806-3p | 13.69 | -0.43 | 0.572119806 | 0.999893527 |
| miR-4797-3p | 13.68 | -0.12 | 0.852263098 | 0.999893527 |
| miR-4684-3p | 13.62 | -0.91 | 0.208831114 | 0.999893527 |
| miR-1227-3p | 13.54 | 0.93 | 0.360675895 | 0.999893527 |
| miR-6833-3p | 13.52 | -0.16 | 0.796911843 | 0.999893527 |
| miR-548ab | 13.45 | 0.00 | 0.999893527 | 0.999893527 |
| miR-3135a-3p | 13.45 | 0.11 | 0.898214728 | 0.999893527 |
| miR-4778-5p | 13.44 | 0.58 | 0.593060478 | 0.999893527 |
| miR-4724-5p | 13.32 | -0.66 | 0.372642549 | 0.999893527 |
| Gly_CCC_leader | 13.30 | -0.81 | 0.227018716 | 0.999893527 |
| Asn_GTT_5half | 13.27 | 1.16 | 0.237595427 | 0.999893527 |
| Ala_CGC_leader | 13.27 | -0.50 | 0.695738136 | 0.999893527 |
| miR-548z-3p | 13.23 | 0.11 | 0.87780082 | 0.999893527 |
| miR-4488-5p | 13.20 | 0.21 | 0.867252893 | 0.999893527 |
| miR-548s-3p | 13.20 | 0.22 | 0.816584256 | 0.999893527 |
| miR-663b-3p | 13.15 | -0.12 | 0.862748289 | 0.999893527 |
| miR-98-3p | 13.13 | -0.11 | 0.893698094 | 0.999893527 |
| miR-548n | 12.97 | 0.16 | 0.828712073 | 0.999893527 |
| miR-4671-5p | 12.96 | 0.43 | 0.535943776 | 0.999893527 |
| miR-551a-3p | 12.84 | 0.19 | 0.779495083 | 0.999893527 |
| miR-6082-5p | 12.84 | 0.31 | 0.751808188 | 0.999893527 |
| miR-6511b-2-3p | 12.83 | 0.11 | 0.889547003 | 0.999893527 |
| Ser_CGA_tRF5 | 12.75 | 0.92 | 0.284875451 | 0.999893527 |
| miR-4268-5p | 12.75 | -0.59 | 0.40643776 | 0.999893527 |
| miR-194-3p | 12.70 | 0.07 | 0.924135373 | 0.999893527 |
| miR-3614-3p | 12.69 | -0.30 | 0.655763892 | 0.999893527 |
| miR-1305-5p | 12.68 | -0.75 | 0.301263141 | 0.999893527 |
| miR-10527-5p | 12.62 | -0.04 | 0.953211538 | 0.999893527 |
| miR-10396a-3p | 12.56 | -0.71 | 0.598634932 | 0.999893527 |
| let-7a-3-5p | 12.44 | 0.38 | 0.704737993 | 0.999893527 |
| miR-9901-3p | 12.39 | -0.61 | 0.63199999 | 0.999893527 |
| Ala_CGC_tRF3 | 12.37 | 0.09 | 0.910633209 | 0.999893527 |
| miR-3145-3p | 12.37 | 0.80 | 0.326226786 | 0.999893527 |
| miR-4781-3p | 12.31 | -0.50 | 0.460826219 | 0.999893527 |
| miR-4436a-3p | 12.31 | 0.02 | 0.977886968 | 0.999893527 |
| miR-378e-3p | 12.29 | -1.09 | 0.236463448 | 0.999893527 |

| | | | | |
|---|---|---|---|---|
| miR-542-5p | 12.24 | -0.61 | 0.363877965 | 0.999893527 |
| miR-3059-3p | 12.23 | 0.10 | 0.904738411 | 0.999893527 |
| miR-3196-5p | 12.23 | 0.76 | 0.539576856 | 0.999893527 |
| miR-1273c | 12.23 | -0.06 | 0.925202867 | 0.999893527 |
| miR-3133-3p | 12.22 | 0.47 | 0.562610862 | 0.999893527 |
| miR-378c-3p | 12.21 | 0.59 | 0.390736503 | 0.999893527 |
| miR-4766-3p | 12.19 | 0.45 | 0.607174548 | 0.999893527 |
| miR-6878-5p | 12.19 | -0.19 | 0.777757891 | 0.999893527 |
| miR-1255b-1-5p | 12.19 | -0.30 | 0.686655657 | 0.999893527 |
| miR-3146-3p | 12.15 | 0.30 | 0.727460004 | 0.999893527 |
| miR-3942-5p | 12.14 | -1.27 | 0.090528618 | 0.999893527 |
| miR-3661-5p | 12.11 | 0.00 | 0.997236288 | 0.999893527 |
| miR-3179-4-5p | 12.07 | 0.21 | 0.871126669 | 0.999893527 |
| miR-939-5p | 12.05 | 0.57 | 0.500508011 | 0.999893527 |
| miR-577-3p | 12.00 | -0.39 | 0.668400024 | 0.999893527 |
| miR-4525-3p | 11.95 | -0.72 | 0.280132807 | 0.999893527 |
| miR-4504-3p | 11.92 | -0.26 | 0.779115179 | 0.999893527 |
| miR-6514-3p | 11.91 | -0.42 | 0.599841046 | 0.999893527 |
| miR-326-5p | 11.90 | -0.25 | 0.713228197 | 0.999893527 |
| miR-3613-3p | 11.89 | 0.98 | 0.339045087 | 0.999893527 |
| miR-7854-5p | 11.88 | -0.56 | 0.414226396 | 0.999893527 |
| miR-153-2-3p | 11.86 | 0.02 | 0.980996038 | 0.999893527 |
| miR-598-5p | 11.85 | -0.14 | 0.840211191 | 0.999893527 |
| miR-6739-5p | 11.85 | 1.43 | 0.147692343 | 0.999893527 |
| miR-623-3p | 11.83 | -1.38 | 0.23003196 | 0.999893527 |
| miR-302b-3p | 11.79 | 0.18 | 0.814415965 | 0.999893527 |
| miR-6790-3p | 11.77 | 0.42 | 0.721817414 | 0.999893527 |
| miR-381-5p | 11.74 | 0.10 | 0.904675288 | 0.999893527 |
| miR-762-5p | 11.71 | 0.51 | 0.596806371 | 0.999893527 |
| miR-12121-3p | 11.65 | -0.20 | 0.78827932 | 0.999893527 |
| miR-2467-5p | 11.63 | -0.56 | 0.565150599 | 0.999893527 |
| miR-1199-3p | 11.58 | -0.03 | 0.960594266 | 0.999893527 |
| miR-379-3p | 11.58 | 1.52 | 0.19456111 | 0.999893527 |
| miR-196a-1-3p | 11.56 | -0.24 | 0.75032257 | 0.999893527 |
| miR-4492-3p | 11.53 | 0.32 | 0.802959842 | 0.999893527 |
| miR-4429-5p | 11.51 | -0.06 | 0.931569415 | 0.999893527 |
| miR-3663-3p | 11.50 | 0.03 | 0.969858779 | 0.999893527 |
| miR-636-5p | 11.48 | 0.69 | 0.359892543 | 0.999893527 |
| miR-6742-3p | 11.46 | 0.04 | 0.954595061 | 0.999893527 |
| miR-24-2-3p | 11.45 | -0.08 | 0.928718289 | 0.999893527 |
| miR-5687-5p | 11.42 | 0.53 | 0.563838751 | 0.999893527 |
| miR-5694-5p | 11.40 | 0.55 | 0.530980107 | 0.999893527 |
| miR-5587-3p | 11.40 | -0.78 | 0.489266837 | 0.999893527 |
| miR-4511-5p | 11.38 | -0.17 | 0.815400019 | 0.999893527 |
| miR-4634-5p | 11.38 | -0.67 | 0.670736919 | 0.999893527 |
| miR-6090-3p | 11.31 | 0.53 | 0.635435109 | 0.999893527 |
| miR-5579-5p | 11.30 | -0.17 | 0.811373353 | 0.999893527 |
| miR-1255b-5p | 11.30 | 0.39 | 0.649605 | 0.999893527 |
| Arg_TCG_leader | 11.25 | 1.14 | 0.32631868 | 0.999893527 |
| miR-4259-5p | 11.22 | -0.05 | 0.94789538 | 0.999893527 |
| miR-320e-3p | 11.21 | 0.31 | 0.64646712 | 0.999893527 |
| miR-4446-3p | 11.21 | 0.60 | 0.431914949 | 0.999893527 |
| miR-3085-3p | 11.19 | -0.39 | 0.724639404 | 0.999893527 |
| miR-3130-1-5p | 11.14 | -0.43 | 0.566001139 | 0.999893527 |
| Thr_TGT_leader | 11.12 | 0.15 | 0.876535571 | 0.999893527 |
| miR-10399-5p | 11.12 | 1.19 | 0.249567198 | 0.999893527 |
| miR-3672-5p | 11.09 | -0.66 | 0.376024608 | 0.999893527 |
| miR-6888-3p | 11.09 | -0.17 | 0.811638715 | 0.999893527 |
| miR-4755-3p | 11.07 | 0.83 | 0.550429452 | 0.999893527 |
| miR-431-3p | 11.06 | 0.29 | 0.825972557 | 0.999893527 |
| miR-1229-3p | 11.03 | 0.35 | 0.701906851 | 0.999893527 |
| miR-1913-5p | 11.02 | 0.86 | 0.258997354 | 0.999893527 |

| | | | | |
|---|---|---|---|---|
| miR-1972-2-3p | 11.01 | 0.47 | 0.678041406 | 0.999893527 |
| miR-4767-5p | 10.98 | -0.09 | 0.908027805 | 0.999893527 |
| miR-1296-3p | 10.97 | -0.81 | 0.253505929 | 0.999893527 |
| miR-187-5p | 10.97 | -0.20 | 0.773563655 | 0.999893527 |
| miR-668-3p | 10.92 | -0.03 | 0.96613099 | 0.999893527 |
| miR-141-3p | 10.91 | -0.14 | 0.840615037 | 0.999893527 |
| miR-12116-3p | 10.91 | 0.27 | 0.763782747 | 0.999893527 |
| miR-6780b-3p | 10.85 | -1.24 | 0.224481721 | 0.999893527 |
| miR-1914-5p | 10.85 | 0.01 | 0.986303489 | 0.999893527 |
| miR-5689-3p | 10.85 | 0.63 | 0.675194132 | 0.999893527 |
| miR-6741-3p | 10.80 | -0.08 | 0.914876419 | 0.999893527 |
| miR-5195-3p | 10.78 | 0.66 | 0.463639989 | 0.999893527 |
| miR-1260a | 10.78 | -0.85 | 0.382702867 | 0.999893527 |
| miR-3944-5p | 10.75 | -0.58 | 0.453091535 | 0.999893527 |
| miR-548ad-5p | 10.71 | -0.42 | 0.644567976 | 0.999893527 |
| miR-8078-5p | 10.67 | -0.42 | 0.555460272 | 0.999893527 |
| miR-3939 | 10.66 | 0.02 | 0.983131995 | 0.999893527 |
| miR-205-5p | 10.62 | 1.40 | 0.259666147 | 0.999893527 |
| miR-548n-3p | 10.61 | -0.21 | 0.774987668 | 0.999893527 |
| miR-3164-5p | 10.61 | -0.39 | 0.599027122 | 0.999893527 |
| Phe_GAA_3half | 10.60 | 1.26 | 0.20149259 | 0.999893527 |
| miR-4424-3p | 10.57 | 0.10 | 0.91241791 | 0.999893527 |
| Tyr_GTA_5half | 10.56 | 0.36 | 0.65442634 | 0.999893527 |
| miR-3131-5p | 10.51 | 0.39 | 0.57116127 | 0.999893527 |
| miR-489-3p | 10.49 | -0.48 | 0.638178721 | 0.999893527 |
| miR-548at-3p | 10.47 | 0.36 | 0.615159165 | 0.999893527 |
| miR-7110-3p | 10.47 | -0.35 | 0.612756008 | 0.999893527 |
| miR-493-5p | 10.46 | 0.07 | 0.953387174 | 0.999893527 |
| miR-6878-3p | 10.45 | 0.11 | 0.899170045 | 0.999893527 |
| miR-4804-5p | 10.44 | 0.57 | 0.4741178 | 0.999893527 |
| miR-4472-1-5p | 10.44 | -0.41 | 0.748737149 | 0.999893527 |
| miR-3115-5p | 10.43 | -0.35 | 0.688834543 | 0.999893527 |
| miR-4443 | 10.42 | 0.98 | 0.350377951 | 0.999893527 |
| miR-1272-5p | 10.39 | -0.15 | 0.854855236 | 0.999893527 |
| miR-548a-3p | 10.33 | -0.50 | 0.521150878 | 0.999893527 |
| miR-1827-3p | 10.33 | -0.02 | 0.987899233 | 0.999893527 |
| miR-935-5p | 10.32 | 0.39 | 0.589187763 | 0.999893527 |
| miR-4472-2-3p | 10.30 | -0.44 | 0.63708518 | 0.999893527 |
| miR-6080-3p | 10.28 | 0.86 | 0.411743149 | 0.999893527 |
| miR-3664-3p | 10.28 | -0.40 | 0.594698446 | 0.999893527 |
| miR-124-1-5p | 10.20 | 0.06 | 0.953008509 | 0.999893527 |
| miR-2355-5p | 10.16 | 0.93 | 0.341282047 | 0.999893527 |
| miR-7111-3p | 10.14 | -0.29 | 0.754498503 | 0.999893527 |
| miR-144-5p | 10.12 | 0.55 | 0.622104162 | 0.999893527 |
| miR-6847-5p | 10.12 | 0.30 | 0.695625194 | 0.999893527 |
| miR-514a-1-3p | 10.10 | -0.31 | 0.757712359 | 0.999893527 |
| miR-29a-5p | 10.09 | -0.17 | 0.837186436 | 0.999893527 |
| miR-10397-5p | 10.09 | -0.58 | 0.599227332 | 0.999893527 |
| miR-4687-3p | 10.04 | -0.17 | 0.815837444 | 0.999893527 |
| miR-3124-3p | 10.04 | 0.34 | 0.699026265 | 0.999893527 |
| miR-4521 | 10.03 | -0.33 | 0.639903995 | 0.999893527 |
| miR-19a-5p | 10.02 | 0.34 | 0.789801162 | 0.999893527 |

Table S3. DESeq2 differential analysis of tRNA halves by SA stress in U2OS wild-type and ANG KO cells (2 biological replicates)

*Table sorted by adjusted p value in WT. To simplify the table, only tRNA halves with MeanExpression(WT) >10 are shown. Log2FoldChange is shaded from red (up-regulated by SA) to blue (down-regulated by SA). Red font highlights the two ANG-dependent tRNA halves validated by Northern blot.*

| | **Wild-type** | | | | **ANG KO C3** | | | | **ANG KO C5** | | | |
|---|---|---|---|---|---|---|---|---|---|---|---|---|
| | **MeanExpression(WT)** | **log2FoldChange(SA/ctrl)** | **pvalue** | **padj** | **MeanExpression(KO C3)** | **log2FoldChange(SA/ctrl)** | **pvalue** | **padj** | **MeanExpression(KO C5)** | **log2FoldChange(SA/ctrl)** | **pvalue** | **padj** |
| Gly_GCC_5half | 15998.89 | 3.56 | 1.76E-20 | 4.46E-18 | 17198.75 | 3.74 | 3.42E-12 | 4.60E-10 | 38650.51 | 4.64 | 9.52E-48 | 2.08E-44 |
| Val_CAC_5half | 3671.88 | 3.42 | 5.29E-18 | 9.40E-16 | 3547.61 | 3.62 | 2.73E-12 | 4.00E-10 | 9905.03 | 4.33 | 7.46E-38 | 5.44E-35 |
| Gly_CCC_5half | 3993.35 | 3.27 | 1.21E-17 | 1.95E-15 | 4523.12 | 2.23 | 1.27E-03 | 2.34E-02 | 10155.57 | 3.40 | 8.14E-20 | 1.37E-17 |
| Glu_CTC_5half | 9679.52 | 2.55 | 4.92E-11 | 3.36E-09 | 6991.62 | 1.96 | 1.90E-05 | 7.68E-04 | 15637.34 | 2.57 | 9.50E-16 | 9.45E-14 |
| Val_AAC_5half | 506.89 | 1.96 | 1.19E-05 | 4.23E-04 | 502.05 | 2.09 | 4.23E-05 | 1.51E-03 | 1100.44 | 2.38 | 9.25E-12 | 5.79E-10 |
| Met_CAT_3half | 102.26 | 2.30 | 2.04E-05 | 6.70E-04 | 122.47 | 3.11 | 3.89E-07 | 2.24E-05 | 385.04 | 3.50 | 1.06E-14 | 9.13E-13 |
| His_GTG_5half | 339.91 | 2.16 | 3.96E-05 | 1.24E-03 | 131.58 | 0.60 | 3.25E-01 | 7.27E-01 | 266.63 | 0.23 | 5.37E-01 | 9.97E-01 |
| iMet_CAT_5half | 24.95 | 3.26 | 3.99E-05 | 1.24E-03 | 29.21 | 3.16 | 3.39E-04 | 8.55E-03 | 82.21 | 3.70 | 2.24E-11 | 1.36E-09 |
| Glu_CTC_3half | 735.27 | 2.02 | 7.50E-05 | 2.22E-03 | 1139.74 | 2.86 | 1.14E-07 | 7.35E-06 | 2180.26 | 2.83 | 6.14E-12 | 3.95E-10 |
| Lys_TTT_5half | 680.88 | 1.92 | 2.18E-04 | 5.49E-03 | 728.96 | 1.45 | 4.78E-03 | 6.27E-02 | 1451.04 | 2.57 | 1.82E-13 | 1.38E-11 |
| Asp_GTC_3half | 7501.37 | 1.52 | 5.13E-04 | 1.12E-02 | 4472.76 | -1.26 | 1.17E-02 | 1.13E-01 | 7478.67 | -0.51 | 1.33E-01 | 6.11E-01 |
| Arg_CCG_5half | 15.80 | 3.10 | 7.17E-04 | 1.46E-02 | 18.58 | 2.97 | 2.24E-03 | 3.64E-02 | 29.90 | 3.01 | 3.00E-05 | 5.62E-04 |
| Lys_CTT_5half | 5079.14 | 1.39 | 1.09E-03 | 2.04E-02 | 5549.23 | 1.03 | 1.38E-01 | 5.68E-01 | 9221.17 | 1.93 | 7.65E-09 | 3.10E-07 |
| Gly_CCC_3half | 1133.67 | 1.42 | 1.58E-03 | 2.65E-02 | 982.00 | 1.17 | 9.20E-02 | 4.43E-01 | 3168.88 | 1.17 | 1.79E-03 | 2.06E-02 |
| Gly_GCC_3half | 98.95 | 1.88 | 1.89E-03 | 3.05E-02 | 131.47 | 1.21 | 4.81E-02 | 3.19E-01 | 315.65 | 1.99 | 6.39E-05 | 1.06E-03 |
| Gly_TCC_5half | 973.66 | 1.14 | 4.04E-03 | 5.98E-02 | 458.00 | -1.30 | 6.50E-03 | 7.59E-02 | 887.55 | -0.77 | 2.26E-02 | 1.75E-01 |
| Pro_CGG_5half | 19.65 | 1.91 | 9.73E-03 | 1.20E-01 | 18.59 | 1.71 | 4.25E-02 | 2.92E-01 | 92.21 | 1.31 | 9.82E-03 | 9.00E-02 |
| Glu_TTC_3half | 470.72 | 1.20 | 1.12E-02 | 1.32E-01 | 458.70 | 1.75 | 2.60E-03 | 4.12E-02 | 997.69 | 1.35 | 4.18E-04 | 5.68E-03 |
| Leu_CAG_5half | 11.41 | 2.51 | 1.45E-02 | 1.56E-01 | 13.53 | 2.60 | 3.05E-02 | 2.32E-01 | 45.71 | 2.98 | 1.66E-05 | 3.39E-04 |
| Asp_GTC_5half | 237.14 | 0.91 | 2.81E-02 | 2.58E-01 | 201.93 | -0.92 | 7.88E-02 | 4.09E-01 | 338.31 | -0.43 | 2.67E-01 | 8.62E-01 |
| Val_TAC_5half | 1455.02 | 0.91 | 4.04E-02 | 3.27E-01 | 1030.64 | -0.52 | 2.81E-01 | 7.04E-01 | 2618.68 | -0.59 | 6.62E-02 | 4.11E-01 |
| Gln_CTG_3half | 123.75 | 1.08 | 5.74E-02 | 4.11E-01 | 80.30 | -1.37 | 3.85E-02 | 2.72E-01 | 160.99 | -1.37 | 1.61E-03 | 1.88E-02 |
| SeC_TCA_5half | 11.05 | 1.75 | 7.13E-02 | 4.52E-01 | 9.77 | 0.56 | 6.42E-01 | 8.70E-01 | 9.62 | 1.77 | 8.64E-02 | 4.74E-01 |
| Glu_TTC_5half | 8864.09 | 0.70 | 7.44E-02 | 4.62E-01 | 3056.80 | -0.55 | 2.68E-01 | 7.03E-01 | 11086.36 | -0.87 | 1.45E-02 | 1.23E-01 |
| Val_TAC_3half | 197.55 | 0.75 | 1.86E-01 | 7.22E-01 | 128.30 | -1.41 | 6.75E-03 | 7.69E-02 | 166.46 | -0.35 | 4.29E-01 | 9.86E-01 |
| Gln_TTG_3half | 35.79 | 0.88 | 1.98E-01 | 7.35E-01 | 18.14 | -1.44 | 1.27E-01 | 5.42E-01 | 59.02 | -2.75 | 1.95E-06 | 4.84E-05 |
| Pro_AGG_5half | 15.33 | -0.90 | 2.70E-01 | 8.01E-01 | 19.47 | -0.29 | 7.47E-01 | 9.11E-01 | 93.49 | -0.78 | 1.20E-01 | 5.84E-01 |
| Pro_TGG_3half | 20.55 | 0.70 | 3.14E-01 | 8.34E-01 | 15.29 | -0.97 | 3.89E-01 | 7.51E-01 | 48.60 | -1.75 | 2.53E-03 | 2.77E-02 |
| Leu_CAA_3half | 10.06 | -0.97 | 3.14E-01 | 8.34E-01 | 14.76 | -0.61 | 4.88E-01 | 8.17E-01 | 22.77 | 0.50 | 4.73E-01 | 9.86E-01 |
| Gln_CTG_5half | 347.16 | 0.41 | 3.33E-01 | 8.36E-01 | 315.54 | -0.34 | 5.84E-01 | 8.59E-01 | 710.99 | -0.57 | 1.14E-01 | 5.70E-01 |
| Ala_TGC_3half | 11.10 | 0.78 | 3.78E-01 | 8.45E-01 | 4.66 | -1.79 | 1.93E-01 | NA | 9.16 | -1.76 | 8.58E-02 | 4.73E-01 |
| Pro_TGG_5half | 19.40 | 0.58 | 4.28E-01 | 8.62E-01 | 18.05 | 0.59 | 4.92E-01 | 8.19E-01 | 89.20 | 0.03 | 9.49E-01 | 9.97E-01 |
| Gly_TCC_3half | 23.20 | 0.66 | 4.41E-01 | 8.65E-01 | 25.12 | -1.64 | 3.72E-02 | 2.67E-01 | 46.62 | -0.81 | 1.60E-01 | 6.88E-01 |
| Pro_CGG_3half | 13.22 | 0.32 | 6.98E-01 | 9.47E-01 | 4.39 | 0.48 | 7.32E-01 | NA | 27.06 | -1.30 | 5.60E-02 | 3.62E-01 |
| Arg_CCG_3half | 17.74 | 0.21 | 7.97E-01 | 9.76E-01 | 12.86 | -0.06 | 9.46E-01 | 9.77E-01 | 40.48 | -1.72 | 3.66E-03 | 3.89E-02 |
| Asn_GTT_3half | 15.70 | 0.18 | 8.23E-01 | 9.78E-01 | 14.46 | -0.92 | 3.23E-01 | 7.27E-01 | 27.18 | -0.56 | 3.80E-01 | 9.60E-01 |
| Ile_AAT_3half | 13.97 | -0.14 | 8.66E-01 | 9.90E-01 | 14.71 | -2.63 | 2.56E-03 | 4.08E-02 | 18.34 | -1.37 | 7.45E-02 | 4.40E-01 |
| Lys_CTT_3half | 11.87 | 0.07 | 9.32E-01 | 9.94E-01 | 13.28 | 0.49 | 5.66E-01 | 8.49E-01 | 33.40 | 0.60 | 3.16E-01 | 8.95E-01 |
| Gln_TTG_5half | 65.04 | -0.01 | 9.82E-01 | 9.98E-01 | 38.64 | 0.52 | 4.59E-01 | 7.96E-01 | 133.90 | -1.21 | 7.75E-03 | 7.41E-02 |
| Lys_TTT_3half | 19.41 | 0.01 | 9.86E-01 | 9.98E-01 | 16.72 | -0.52 | 5.54E-01 | 8.45E-01 | 42.60 | -1.09 | 9.00E-02 | 4.88E-01 |
